# Design, Synthesis, Molecular Docking and Biological Activity of Pyrazolo[3,4-*b*]pyridines as Promising Lead Candidates Against *Mycobacterium tuberculosis*

**DOI:** 10.1101/2023.06.26.546517

**Authors:** H. Surya Prakash Rao, R. Gunasundari, Lakshmi Narayana Adigopula, Jayaraman Muthukumaran

**Affiliations:** Department of Chemistry, Pondicherry University, Puducherry 605 014. INDIA; Department of Biotechnology, Sharda School of Engineering and Technology, Sharda University, Greater Noida. INDIA; Vasista Pharma Chem, ALEAP, Pragathi Nagar, Kukatpalli, Hyderabad 500 090. INDIA

## Abstract

Pyrazolo[3,4-*b*]pyridine is a medicinally privileged structure. We have achieved a new and facile synthesis of a combinatorial library of its tetra- and persubstituted derivatives by trifluoracetic acid catalysed condensation of a group of 5-aminopyrazoles and a group of α-oxoketene dithioacetals. Furthermore, we demonstrated structural modification of the products via reductive desulfurization, hydrolysis of the ester and Suzuki coupling of the bromo derivative with aryl bornoic acids. Some of the products were subjected to *in vitro* MABA assay against *M. tuberclulosis* H37Rv strain and *in silico* analysis by binding to pantothenate synthetase from *M. tuberclulosis* (MTBPS). The results indicated that the pyazolo[3,4-*b*]pyridine with N(1)CH_3_, C(3)C_6_H_5_, C(4) *p*CH_3_C_6_H_5_, C(5)CO_2_Et, C(6)SMe substitutions exhibits promising antituberculotic activity.

## 1. Introduction

Synthesis and biological evaluation of a combinatorial library of heterocyclic compounds built on a medicinally privileged scaffold is a well-accepted practice in the drug discovery regimen.^1^ Amongst them, heterocyclic aromatic compounds with multiple nitrogen atoms have garnered greater attention as they exhibit extremely useful biological properties.^2^ Pyrazolo[3,4-*b*]pyridine **1** (Figure 1), a bicyclic heterocyclic aromatic compound with three nitrogen atoms present at 1, 2 and 7 positions, is one such scaffold.^3^ It is embellished with two basic and hydrogen bond acceptor (HBA) nitrogen atoms, and one hydrogen bond donor (HBD) NH (See structure **1**, Figure 1). Structurally, the scaffold is aromatic, stable, flat and is an isostere of naphthalene. Of the two fused aromatic rings, the pyrazole portion in **1** is electron rich, and the pyridine portion is electron deficient; hence, unlike naphthalene it is polarized. The ring allows structural modifications at five positions namely *N*(1), *C*(3), *C*(4), *C*(5) and *C*(6), either during the ring formation or at the later stage of functional group modification.^4^ The scaffold, thus, provides enormous opportunities for the synthesis of a combinatorial library of new chemical entities (NCEs) for exploring structure activity relationships (SAR).^5^ Versatility of the scaffold and diverse biological activity of its derivatives make **1** a medicinally privileged.^6^ Molecules built on **1** exhibit broad spectrum of useful medicinal properties. For example, the drugs like etazolate **2**, tracazolate **3**, cartazolate **4**, built on the scaffold **1**, are used for reduction of anxiety and convulsions.^7^ The drugs enhance binding of γ-aminobutyric acid (GABA), a neurotransmitter, to the GABA specific receptors.^8^ They are less potent than benzodiazepines but are much less sedative, hence preferred. Glicaramide **5** is a second-generation sulfonylurea-based antidiabetic drug.^9^ Vericiguat **6** is a guanylate cyclase receptor agonist. It has been approved recently (2021) for treatment of the patients with the possibility of chronic heart failure.^10^ Its sibling riociguat is used extensively as an independent of NO simulator of soluble guanylate cyclase (sGC).^11^ Besides above examples, molecules with the pyrazolo[3,4-*b*]pyridine core have been used for the treatment of gastro-intestinal diseases,^12^ Alzheimer disease^13^ alcohol withdrawal symptoms^14^ and erectile dysfunction.^15^ Specifically, molecules built on this scaffold have been reported as selective inhibitors of A1 adenosine receptors,^16^ phosphodieste<u>rase</u> 4 (PDE 4) inhibitors in immune and inflammatory cells,^17^ glycogen synthase kinase-3 (GSK-3) inhibitors,^18^ and kinase inhibitors of p38α.^19^ Naturally, medicinal chemists in both industry and academia, have put high focus to develop newer methods of its synthesis.^20^

**Figure 1.**
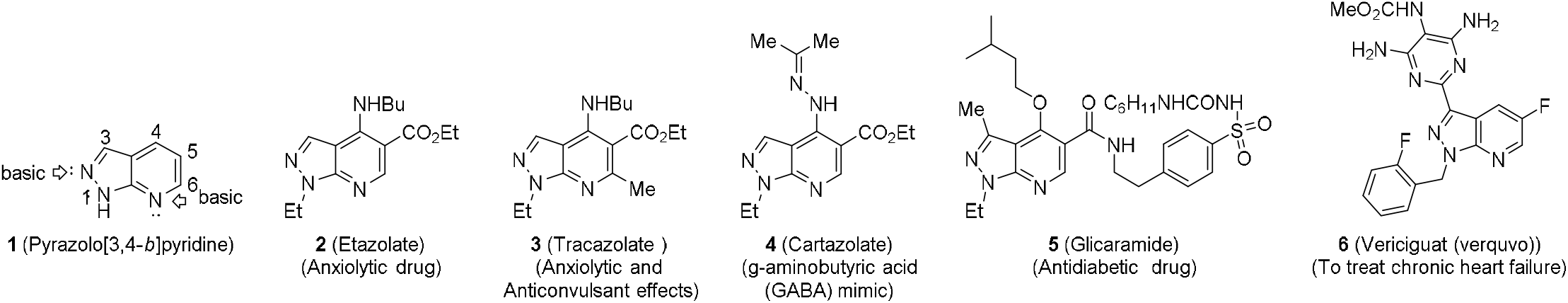
Structures of pyrazolo[3,4-*b*]pyridine **1**, etazolate **2**, tracazolate **3**, cartazolate **4**, glicaramide **5** and vericiguat **6**.

Although the pyrazolo[3,4-*b*]pyridine structure can be assembled by a variety of methods, condensation of *N*(1) substituted 5-aminopyrazoles with 1,3-bis-electrophilic three-carbon synthons is the popular choice (Equation 1, Scheme 1).^21^ As an example, first step in the Bare’s synthesis of etalozolate **2** and cartazolate **4** went through a condensation of 5-aminopyrazole **7** with the masked 1,3-carbonyl compound **8**. The condensation provided **9** which upon cyclization under acidic conditions furnished the pyrazolo[3,4-*b*]pyridine **10**.^22^ Alternatively, the *N*(1) substituted 5-aminopyrazoles **7** can be generated *in situ* from ethyl hydrazine and ethyl cyanoacetate.^23^ We have previously reported that the condensation of 5-amino-3-phenylpyrazoles **11** with 4*H*-chromenes **12** in EtOH reflux provided dihydropyazolo[3,4-*b*]pyridines **13** (Equation 2, Scheme 1).^24^ In this remarkable transformation, initially formed C-alkylated 4*H*-chromene rearranged to **13** via chromene ring opening followed by recyclization. Dehydrogenation of **13** with 2,3-dichloro-5,6-dicyano-1,4-benzoquinone (DDQ) provided pyrazolo[3,4-*b*]pyridines **14**. This two-step method for the synthesis of a combinatorial library pyazolo[3,4-*b*]pyridines **13** was facile and high yielding, but its applicability was limited to those with an ortho-substituted phenolic group at C(4). In our quest to unearth facile synthesis of analogues of etazolate **2**, we planned the condensation of 5-aminopyrazoles **7** with α-oxo ketene dithioacetals **15** (OKDTAs) as an alternate and hitherto unknown method for the synthesis of pyrazolo[3,4-*b*]pyridines (Equation 3, Scheme 1).

**Scheme 1.**
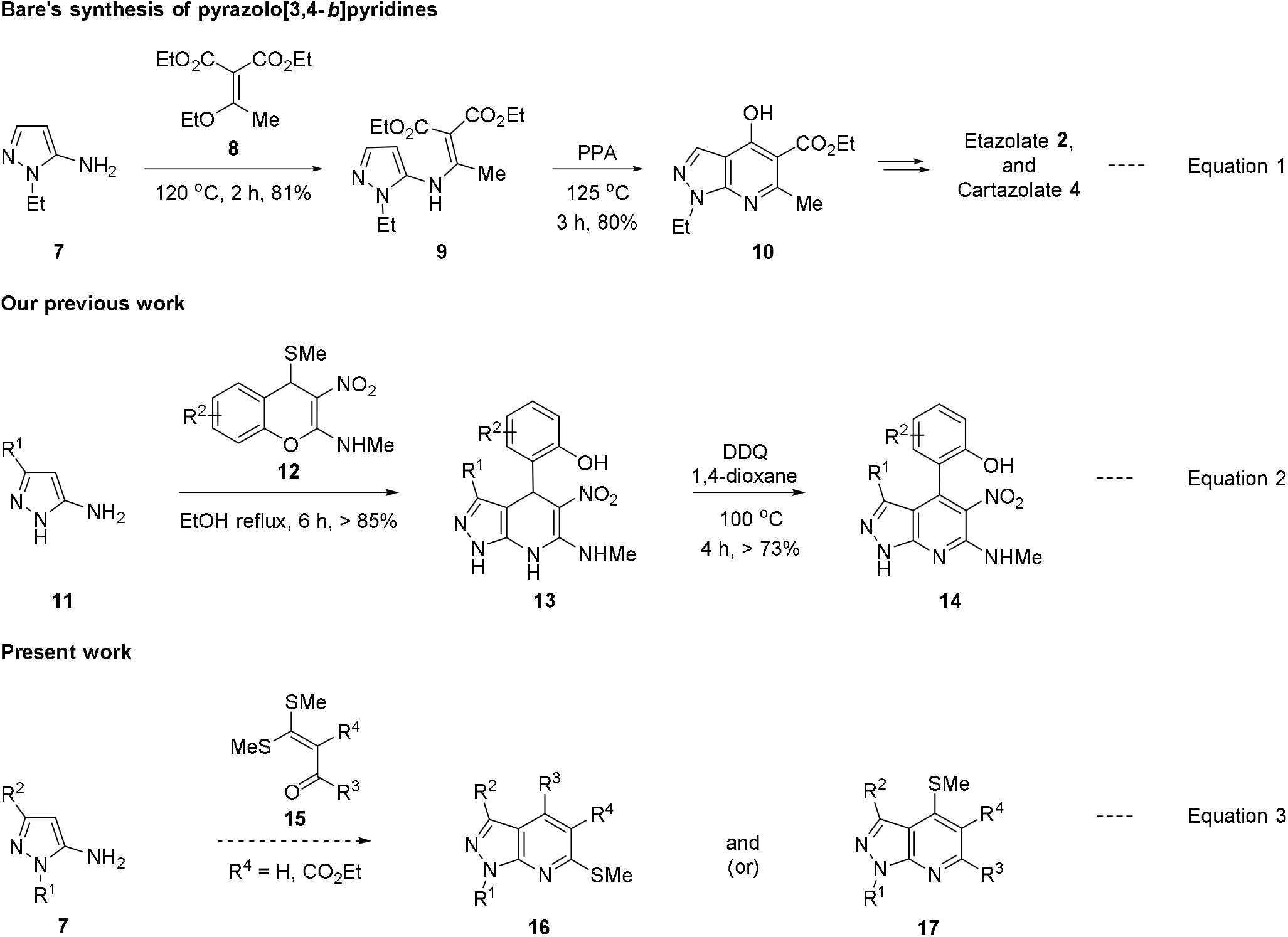
Synthetic studies towards pyrazolo[3,4-*b*]pyridines.

The OKDTAs **15** (Figure 2) are three-carbon synthons, extremely useful in the synthesis of a wide variety of heterocyclic compounds.^25^ The ketene dithioacetal functional group in OKDTAs **15** is a polarized push-pull alkene, primarily, due to the electron withdrawing carbonyl and the electron donating SMe, located in 1,2-positions of ethylene.^26^ Owing to the presence of an α,β-unsaturated ketone sub-structure, **15** is an excellent Michael acceptor. The alkylsulfanyl group in it is open for substitution with nucleophiles *via* substitution nucleophilic-vinylic pathway (S_N_V). We reasoned that since 5-aminopyrazoles **7** possess two nucleophilic centers (NH_2_ and C(4)) and the OKDTAs **15** possess two electrophilic centers (C(1) and C(3)) the reaction between them could lead to either or both of the regio-isomeric pyrazolo[3,4-*b*]pyridines **16** and (or) **17** (Equation 3, Scheme 1). The product pyrazolo[3,4-*b*]pyridines **16** and (or) **17** will still have one more SMe group. Displacement of the residual SMe with another nucleophile will provide a new set of tetra-substituted pyrazolo[3,4-*b*]pyridines. Thus, our plan (Equation 3, Scheme 1) will provide a combinatorial library of pyrazolo[3,4-*b*]pyridines with at least four points of diversity within a two-step reaction sequence from easily and readily available starting compounds. Furthermore, we planned to conduct functional group manipulation of **16**/**17** and evaluate NCEs for their biological activity in silico and in vitro for anti-tuberculosis (anti-TB) activity.

**Figure 2.**
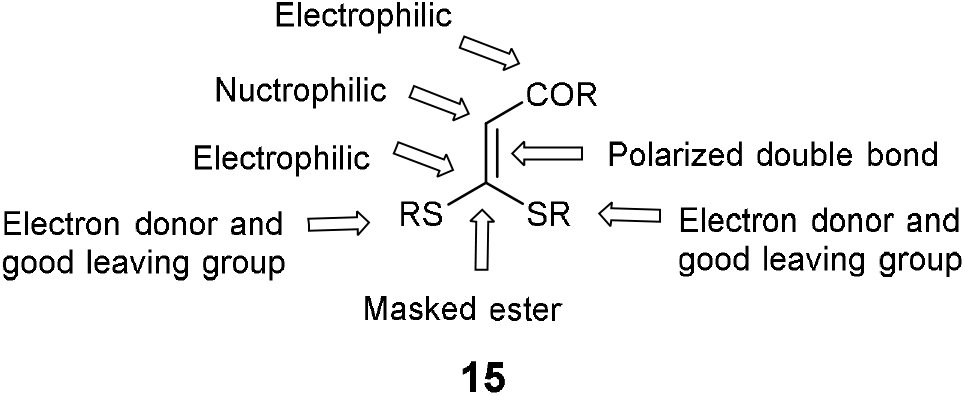
Structural features and reactivity pattern of α-oxo ketene dithioacetals (OKDTAs).

*Mycobacterium tuberculosis (*family *Mycobacteriaceae)* is a highly infectious bacterium. It causes tuberculosis (TB), a debilitating disease affecting more than 2 billion people a year, out of which many are passive carriers.^27^ The bacterium is an opportunistic pathogen specially affecting those with co-morbidities like HIV or AIDS.^28^ It has an unusual waxy coating on its cell surface, primarily due to the presence of mycolic acids as one of the interlinking scaffolds.^29^ Mycolic acids are 60-90 carbon long fatty acids with α-hyroxycarboxlic acid functional group. Presence of mycolic acids in the cell wall makes the bacterium hydrophobic, sturdy, and resistant to desiccation. Interfering in the biosynthesis of mycolic acids may be a key to arresting its virulence. Pantothenate synthetase (MTBPS), an enzyme involved in the biosynthesis of pantothenate (pantothenic acid; vitamin B5; water soluble and essential nutrient) and fatty acids is present in the bacterium but not in humans. The enzyme catalyses ATP-dependent condensation of D-pantoate and β-alanine to form the amide bond of pantothenate. Pantothenate is a precursor for Coenzyme A (CoA). The enzyme participates in the biosynthesis of fatty acids that lead to mycolic acids.^30^ Thus, MTBPS inhibitors could act as anti-TB drugs. Presently, multi-drug therapy in the first line of treatment for TB consists of a combination of antibacterial agents like isoniazid, rifampin, ethambutol, pyrazinamide. If the first line of treatment fails, drugs like bedaquiline, linezolid are used. Still MDR (multi-drug resistant), XDR (extremely drug resistant) and totally drug-resistant (TDR) strains are emerging for which new drugs are required. We believe that suitably substituted NCEs like pyrazolo[3,4-*b*]pyridines **16** hold promise by targeting MTBPS. Herein we report a new method for the synthesis of coveted pyrazolo[3,4-*b*]pyridine scaffold and characterization of several NCEs built on it. Furthermore, we report *in silico* evaluation of the of the best NCEs against MTBPS and their *in vitro* evaluation to unearth a lead for the treatment of TB.

## 2. Results and Discussion

### 2.1. Synthesis of pyrazolo[3,4-b]pyridines

We selected the condensation of 1,3-diphenyl-5-aminopyrazole **7a** with OKDTA **15a** to evaluate feasibility of our proposal and to optimize the reaction conditions (Scheme 2, Table1). We premised that the first and second steps in the condensation, namely, nucleophilic displacement of SMe in OKDTA by the C(5) primary amine or C(4) of the pyrazole followed by cyclization to generate the annulated pyridine ring, will be promoted by an acid catalyst. Singh and co-workers employed acetic acid in the condensation of 1,3-diphenyl-5-aminopyrazole with 1,3-bis-electrophilic 1,3-diketones.^31^ In the first attempt, we conducted the condensation of **7a** and **15a** in acetic acid (pka = 4.76 and b.p = 118 °C) medium. The reaction did not take place at rt or at reflux temperature of acetic acid. The reaction, regrettably, led to decomposition of the OKDTA **15a**. Next, we tried the reaction in toluene reflux while employing acetic acid in stoichiometric amount, but the reaction did not proceed (Table 1, entry 1). We reasoned that activation of the carbonyl group in OKDTA requires stronger acid. Accordingly, we sequentially employed stoichiometric amounts of chloroacetic acid (pKa = 2.86; entry 2)^32^, fluoroacetic acid (pKa = 2.6, entry 3), trichloroacetic acid (pKa = 0.65, entry 4), and trifluoroacetic acid (TFA, pKa = 0.23, entry 5). The condensation reaction worked with trifluoroacetic acid to provide the regio-isomeric pyrazolo[3,4-*b*]pyridines **16a** and **17a** in about 85% yield in the ratio of about 3:1 (entry 5). By conducting different experiments with varying the amount of trifluoroacetic acid in toluene reflux, we discovered that 30 mol% of trifluoroacetic acid is the optimal for the best yield (89%), shortest time (6 h) and cleanest product (entry 6). The reaction with 10 mol% of TFA led to lowered yield (80%) and longer time (12 h; entry 7) without change in ratio. To evaluate if the ratio of the isomers changed with the acid employed, we attempted several reactions in toluene reflux with an organic solvent soluble strong protic acid and Lewis acids. The reaction with 30 mol% of CF_3_SO_3_H (pKa = −14), a very strong protic acid, resulted in lowering of the yield with minor change in the ratio (entry 8). The very strong acid possibly protonated the primary amine and converted it into its ammonium salt, which prevented the nucleophilic attack on the OKDTA. The reaction conducted with 1 equivalent of *p*-toluenesulfonicacid (PTSA, entry 9) resulted in regio-isomers **16a** and **17a** in 1:1 ratio in 82% of yield. However, the reaction took 12 h for completion and products were not clean even after chromatographic purification. The reaction proceeded partially (50%) when we employed 0.5 equivalent of PTSA (entry 10). Finally, we conducted the reaction in presence of the Lewis acids like FeCl_3_ (entry 11), ZnCl_2_ (entry 12), SnCl_4_ (entry 13), CuI (entry 14), Cu(OAc)_2_ (entry 15), and Cu(OTf)_2_ (entry 16) in refluxing toluene. While the reaction did not take place with ZnCl_2_, SnCl_4_, SnCl_2_, CuI, and Cu(OTf)_2_ it proceeded partially at a slower rate in FeCl_3_ and Cu(OAc)_2_. The work-up of the reaction with TFA after its completion was simple. Toluene and TFA (b.p. = 72 °C) were removed in a rotavap under reduced pressure and re-used for the next run.

**Scheme 2.**
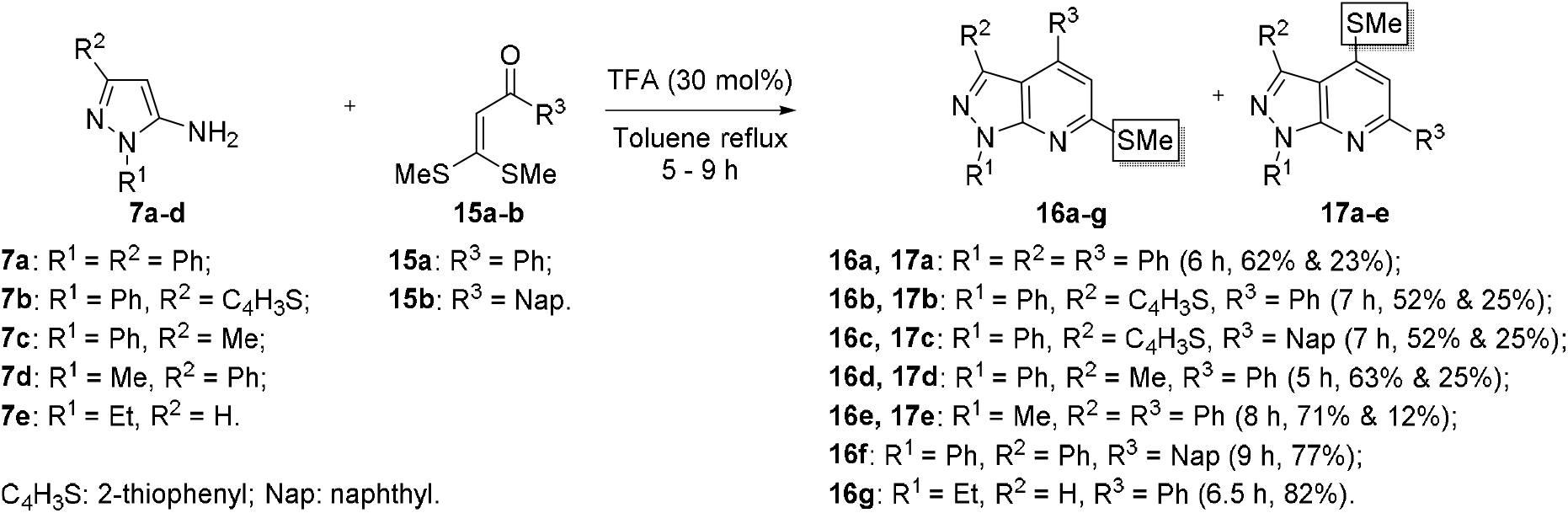
Synthesis of pyrazolo[3,4-*b*]pyridines **16a-g** and **17a-e** from 5-aminopyrazol **7a-e**, and OKDTAs **15a-b.**

**Table 1.** Screening of acid catalysts for the condensation of 5-aminopyrazol **7a** and OKDTA **15a**.

| Entry | Acid | Equiv | Time (h) | 15a:16a | Yield (%) |
| --- | --- | --- | --- | --- | --- |
| 1 | AcOH | 1.0 | 24 |  | nr |
| 2 | ClCH<sub>2</sub>CO<sub>2</sub>H | 1.0 | 24 |  | nr |
| 3 | FCH<sub>2</sub>CO<sub>2</sub>H | 1.0 | 18 |  | nr |
| 4 | Cl<sub>3</sub>CCO<sub>2</sub>H | 1.0 | 24 |  | nr |
| 5 | F<sub>3</sub>CCO<sub>2</sub>H | 1.0 | 06 | 3:1 | 85 |
| **6** | **F<sub>3</sub>CCO<sub>2</sub>H** | **0.3** | **06** | **3:1** | **89** |
| 7 | F<sub>3</sub>CCO<sub>2</sub>H | 0.1 | 12 | 3:1 | 80 |
| 8 | CF<sub>3</sub>SO<sub>3</sub>H | 0.3 | 05 | 3:2 | 61 |
| 9 | PTSA | 1.0 | 12 | 1:1 | 82 |
| 10 | PTSA | 0.5 | 16 | 1:1 | 32* |
| 11 | FeCl<sub>3</sub> | 0.3 | 10 | 3:2 | 42<sup>\$</sup> |
| 12 | ZnCl<sub>2</sub> | 0.3 | 12 |  | nr |
| 13 | SnCl<sub>4</sub> | 0.3 | 12 |  | nr |
| 14 | CuI | 0.3 | 24 |  | nr |
| 15 | Cu(OAc)<sub>2</sub> | 0.3 | 20 | 1:1 | 40 |
| 16 | Cu(OTf)<sub>3</sub> | 0.3 | 24 |  | nr |

nr = No-reaction; \* 50% conversion; §55% conversion

We separated the isomers **16a** and **17a** by column chromatography and characterized them from the analysis of the spectroscopic (IR, ^1^H NMR, ^13^C NMR, DEPT-135 and 2D NMR) and high-resolution mass spectral data. Their 2D NMR (HSQC and HMBC) spectra were particularly useful for structural assignment of the regio-isomers. The HMBC 2D NMR spectrum of **16a** showed cross-peak for hydrogens of C(6)SCH_3_ (2.7 ppm) to the C(6) carbon (160.7 ppm) which was the most down-field shifted signal in the ^13^C NMR spectrum. As anticipated, the lone singlet in the ^1^H NMR spectrum for C(5)H and located at 7.05 ppm also displayed cross-peak for C(6) located at 160.7 ppm. Final confirmation of the structural assignment came from the analysis of the single crystal X-ray crystallographic diffraction data of **16a** (Figure 3). The solid state structures showed that the C(4) phenyl group is twisted out of mean plane of the pyrazolo[3,4-*b*]pyridine to avoid peri-interactions with the phenyl / methyl group at C(3). On the other hand, the N(1) phenyl is in plane with the core structure of pyrazolo[3,4-*b*]pyridine, possibly because of weak but stabilizing interactions between CH and lone pair on pyridine nitrogen.

**Figure 3.**
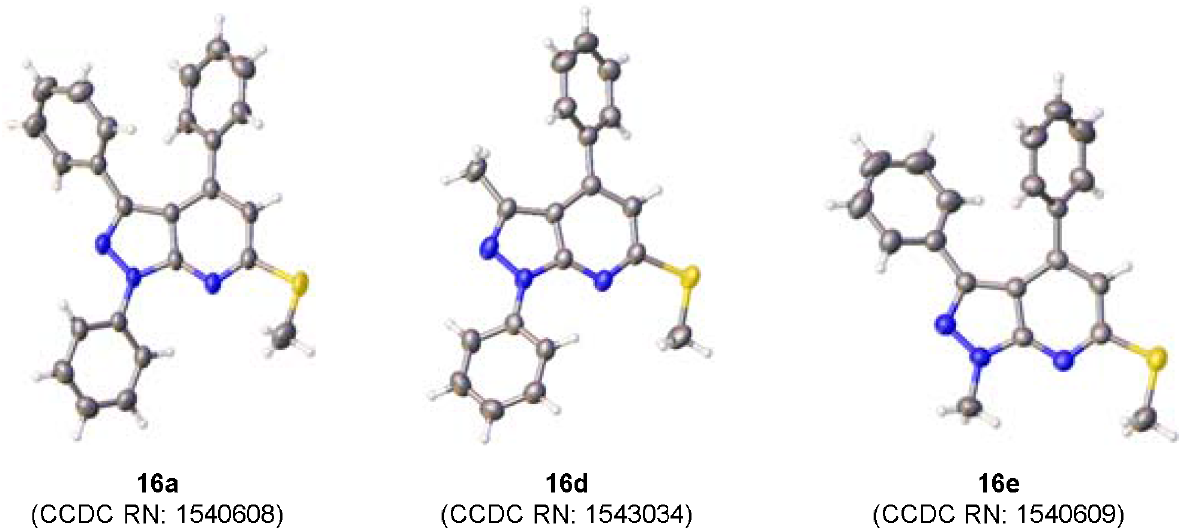
The X-ray crystal structures of the pyrazolo[3,4-*b*]pyridines **16a**, **16d** and **16e**; thermal ellipsoids are set at 50% probability.

Next, we sought to generalize the synthesis of coveted pyrazolo[3,4-*b*]pyridines **16** by reacting a set of five *N*(1)substituted-5-aminopyrazoles **7a-e** and two OKDTAs **15a-b** (Scheme 2). We intended to evaluate influence of the substituents at the *N*(1) and C(3) positions of the pyrazoles and steric bulk of the aryl groups in OKDTAs on the regiochemical outcome of the reaction. Furthermore, **7e** with (N(1) Et) was selected based on the likeness to the drugs **2**-**4**. The condensation of the pyrazole **7b,** which has C(3) thiophenyl group, with OKDTAs **15a-b** delivered pyrazolopyridines **16b-c** and **17b-c** in the ratio of 17:8 in 77% combined yield in both the cases. Higher percentage of **17b** compared to **17a** could be due to the electron donating abilities and lower steric bulk of the C(3) thiophenyl group in **7b** compared to the C(3) phenyl group in **7a**. The condensation of the pyrazole **7c**, which has the C(3) methyl group, with the OKDTA **15a** provided pyrazolopyridines **16d** and **17d** in the ratio of 18:7 in 88% combined yield. The condensation of the pyrazole **7d**, which has N(1)Me, with the OKDTA **15a** furnished **16e** and **17e** in the ratio of 43:7 in 83% combined yield. Higher percentage of the isomer **16e** compared to the earlier experiments indicated that the electron-donating methyl group present on *N*(1) greatly influenced the reaction. The reactions of the pyrazole **7a** and the pyrazole **7e** with the OKDTA **15a** and **15b** respectively provided the regio-isomers **16f-g** exclusively in excellent yield. Above results indicated that the condensation of the pyrazoles with an electron-donating alkyl group on *N*(1) and OKDTAs with a bulky group furnish the regio-isomer **16**. In each case the isomers were separated and characterized. The ^1^H, ^13^C NMR and DEPT-135 of **16b-g** and **17b-e** matched well with the parent pyrazolopyridines **16a** and **17a**. Additionally, X-ray crystal structures of **16d** and **16e** further confirmed the assigned structures (Figure 3).

Non-arrival of a single isomer of the tetrasubstituted pyrazolo[3,4-*b*]pyridines in the above reactions, prompted us to look for modifications in the parent OKDTA **15**. We designed OKDTA **15c** (R^3^ = aryl / alkyl; R^4^ = CO_2_Et; Scheme 1, equation 3) with an ester group on C(2) to increase the electrophilic character of C(3). Furthermore, we reasoned that the product from the reaction of 5-aminopyrazoles with the OKDTA **15c** will provide the pyrazolo[3,4-*b*]pyridines with an ester group at C(5) as seen in drug molecules like etazolate **2**.

In contrast to the condensation reaction of the 5-aminopyrazole **7d** with OKDTA **15a**, its reaction with **15c** delivered the pyrazolo[3,4-*b*]pyridine **16h** exclusively (Scheme 3); without any trace of its isomer **17h**. The structure of **16h** was established by spectroscopic (^1^H, ^13^C, DEPT-135 and 2D NMR) and analytical (HRMS) methods. The 2D NMR spectral data, particularly, HMBC and HSQC were used in assigning the structure. Arrival of a single regio-isomer **16h** in the reaction prompted us to generalize the reaction for the synthesis of etazolate **2** analogues. In this effort, we prepared a library of twenty two pyrazolo[3,4-*b*]pyridines **16h-aa** by condensing a set of five 5-aminopyrazoles **7a**, **7d-h** and a set of five OKDTAs **15c-g** in a combinatorial fashion (Scheme 3). Each reaction worked well to provide pyrazolo[3,4-*b*]pyridines **16** as a single regioisomer in more than 80% yield. Structure of the products emerged from the analysis of spectroscopic and analytical data. In addition, we determined the single crystal X-ray structures of pyrazolo[3,4-*b*]pyridines **16k** and **16aa**, by which we unequivocally confirmed the structures (inset in Scheme 3). Substitution in the N(1) of the 5-aminopyrazole **7** included aliphatic (Me) and aromatic (Ph) groups. Substitution in the C(3) aryl ring included Cl (electron withdrawing, ortho-, para-directing, ability to form weak-hydrogen bonds), Br (electron withdrawing and facilitator in the coupling reactions) and MeO (electron donating and hydrogen bond acceptor group). Substitutions in the OKDTAs **15** included phenyl, 4-chlorophenyl (electron deficient aromatic ring, Cl group forms weak hydrogen bonds), 4-bromophenyl (electron deficient aromatic ring, Br group facilitates coupling reactions), 4-methoxyphenyl (electron rich aromatic ring and OMe is a hydrogen bond acceptor) and methyl (electron donor).

**Scheme 3.**
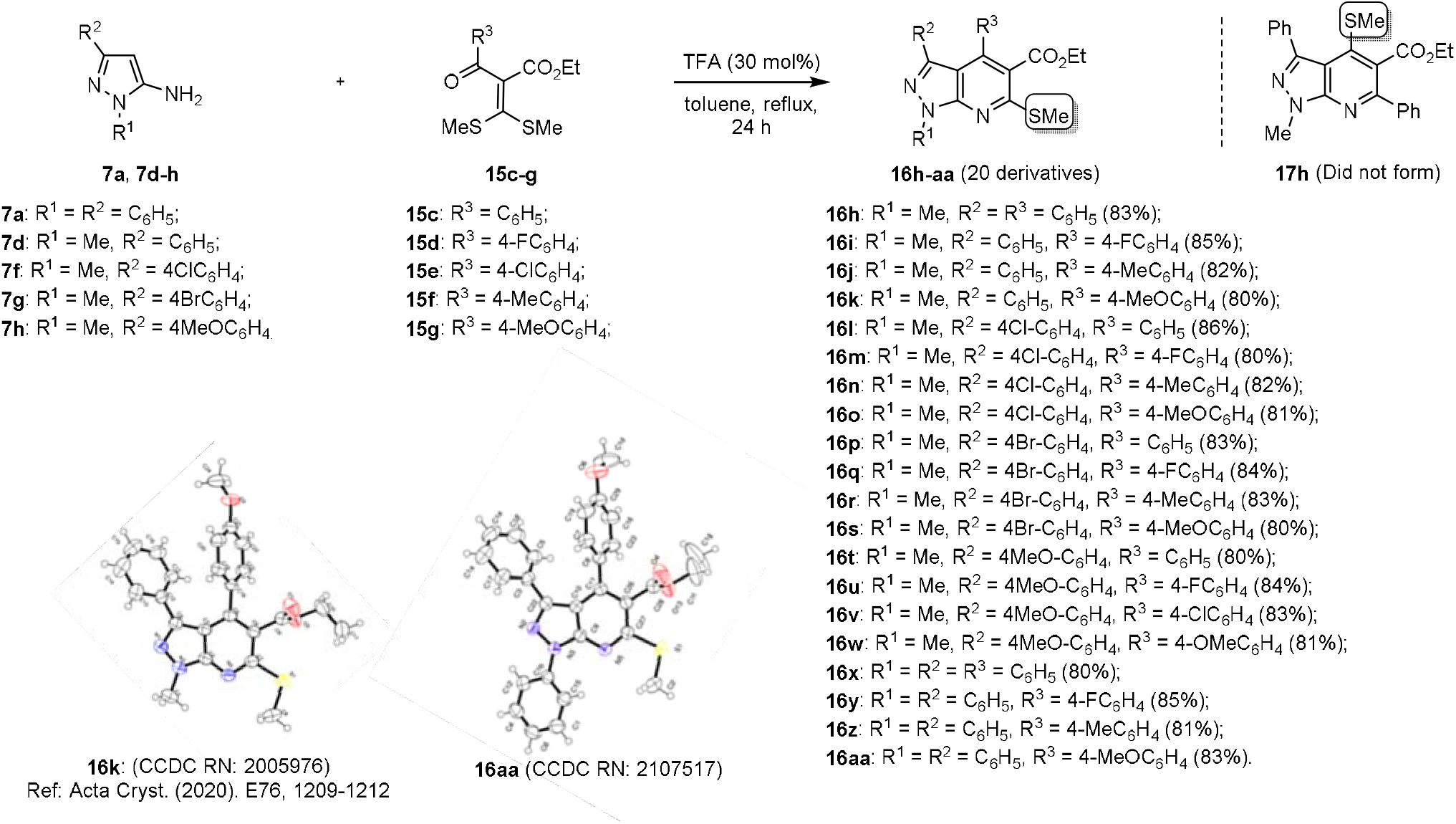
Synthesis of a combinatorial library of pyrazolopyridines **16h-aa** with C(6)SMe and single crystal X-ray structures of pyrazolo[3,4-*b*]pyridines **16k** and **16aa** (thermal ellipsoids are set at 50% probability).

Next, we considered condensation of the 5-aminopyrazoles **7f** and **7h** and with the OKDTA 1**5h** (Scheme 4). We prepared the OKDTA **15h** in good yield from ethyl acetoacetate, carbon disulphide and methyl iodide using NaH as the base.^33^ Condensation of the 5-aminopyrazole **7f** with OKDTA 1**5h**, surprisingly provided the **17f** exclusively. It has the SMe group at C(4) position instead of C(6) as seen in the cases of **16h-aa** - that is, **16ab** did not form, indicating a switch in the regio-chemistry when OKDTA has an alkyl group at C(1). The ^1^H NMR spectrum of **17f** displayed a singlet for SMe at 1.91 ppm, about 0.8 ppm up field from the one seen for C(6)SMe in 1**6h**. The CH_2_ group for the ethyl ester displayed a quartet at 4.47 ppm and this signal is downfield shifted to the corresponding signal in 1**6h** by about 0.4 ppm. In the ^13^C NMR spectrum the signal for C(6) appeared at about δ 155 ppm, instead of 159 for the series in which the SMe was on C(6). Its 2D HMBC NMR spectrum displayed cross peaks for C(6)Me to C(6) and N(Me) to C(7a), thereby confirming assigned structure. We repeated the condensation on **7h** and **15h** and found that the reaction also provided a single product **17g**. Thus, it appears that the subtle changes made in the electrophilicity on the carbonyl carbon of **15** (Ph vs Me) lead to drastic changes in the regio-chemical outcome in its condensation with 5-aminopyrazole **7**.

**Scheme 4.**
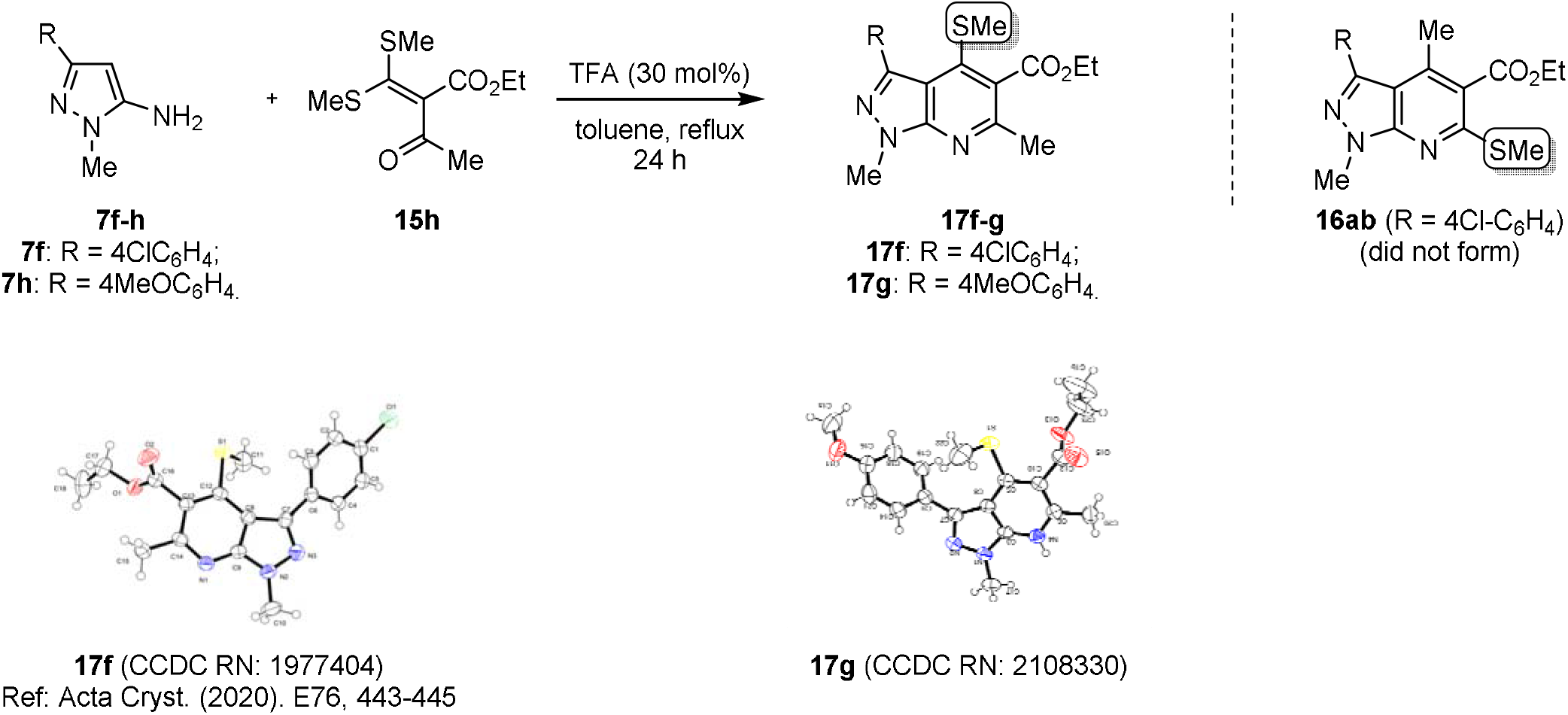
Condensation of 5-aminopyrazoles **7f-h** and with OKDTA **15h** to form the regioisomeric pyrazolopyridines **17f-g** with C(4)SMe and single crystal X-ray structures of pyrazolo[3,4-*b*]pyridines **17f** and **17g** (thermal ellipsoids are set at 50% probability).

Proposed mechanism for the condensation of 5-aminopyrazoles **7** and OKDTA **15c**/**15h** to form pyrazolo[3,4-*b*]pyridines **16**/**17** is given in Scheme 5. Compared to the OKDTAs **15a-b** the C(3) carbon of the OKDTAs 1**5c** is more electrophilic and harder owing to presence of the ester (COOEt) on C(2) carbon. Moreover, the acid catalyst enhances the electrophilic nature of C(3) by activating the carbonyl oxygen. Hence, first step in the reaction between **7** and **15c** is the nucleophilic attack of the harder base, the primary amine at C(5) of **7** on the C(3) carbon of OKDTA **15c** in conjugate fashion to provide the intermediate **18** (Equation 1, Scheme 5). Loss of MeSH led to **19** and cyclization via nucleophilic attack of C(4) of pyrazole to the carbonyl carbon of OKDTA leads to **20**. Finally, loss of water from **20** furnished regio-isomer pyrazolo[3,4-*b*]pyridines **16**. On the other hand, compared to the OKDTA **15c** the C(3) carbon of the OKDTA **15h** is less electrophilic owing to electron donating Me group present on the carbonyl carbon and thus behaves like a soft electrophile. Consequently, first step in the reaction between the pyrazole **7** and **15c** was the attack of the soft nuclophile - that is C(4) carbon of the 5-aminopyrazole - on the C(3) carbon of **15h** in conjugate fashion to form the intermediate **21** (Equation 2, Scheme 5). Loss of MeSH led to **22**. Re-aromatization of the pyrazole ring through tautomerism provided **23**. Both these steps, namely, loss of MeSH and the tautomerization are expected to be very fast and concomitant. The intramolecular attack of the primary amine in **23** to the carbonyl carbon led to cyclization and form **24**. Finally, loss of protonated water delivered the pyrazolo[3,4-*b*]pyridines **17**.

**Scheme 5.**
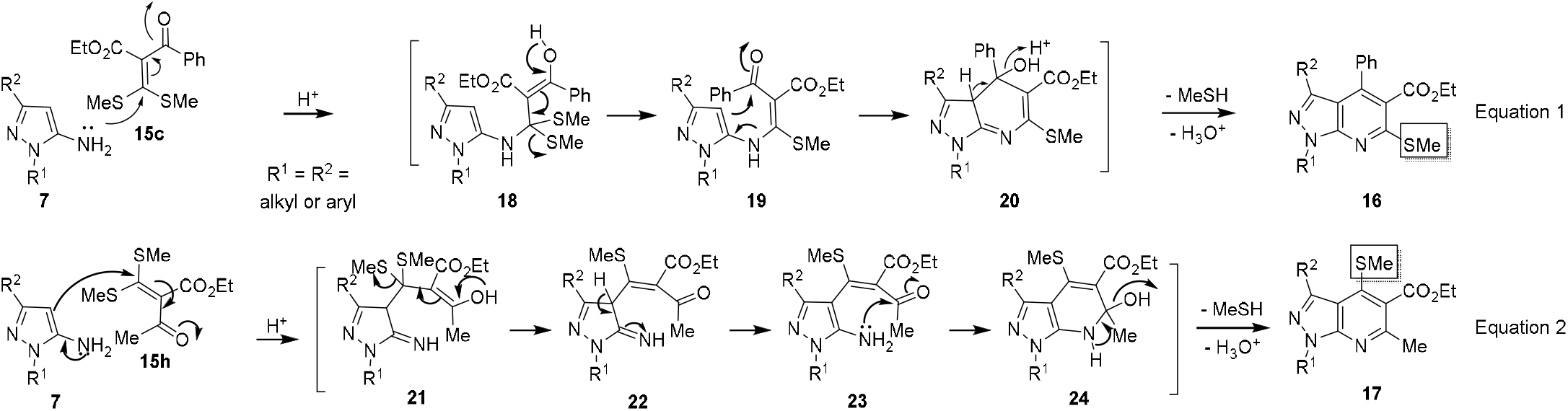
Proposed mechanism for the condensation of 5-aminopyrazoles **7** and OKDTAs **15c**/**15h** to form pyrazolopyridines **16** or **17**.

### 2.2. Modifications in the structures of persubstituted pyrazolo[3,4-b]pyridines

The pyrazolopyridines like **16** are amenable for further structural modifications (Figure 4). For example, (i) the C(4) SMe group can be reductively removed or substituted with C/N/O nucleophiles, (ii) the C(5) ester can be hydrolyzed to acid which can be further converted into esters / amides, and (iii) the 4-BrC_6_H_4_ substitution at C(1), C(3) and C(6) positions can be subjected to Suzuki coupling with a wide variety of aryl/alkenyl boronic acids.

**Figure 4.**
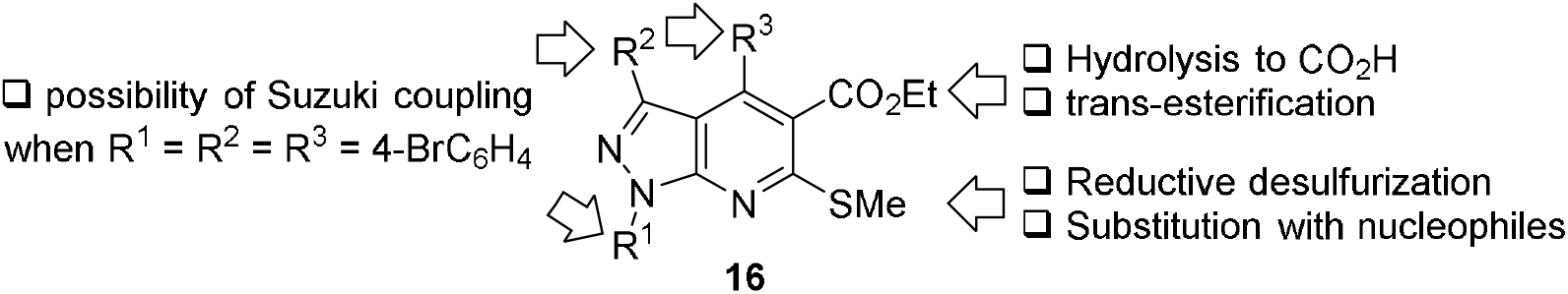
Possibilities for structural modification of persubstituted pyrazolo[3,4-*b*]pyridines **16**.

First among our plans for structural modifications of **16** was the reductive desulfuraization^34^ to furnish pyrazolo[3,4-*b*]pyridines **25** (Scheme 6). The reaction of **16h** with Raney Ni under hydrogen atmosphere in the mixture of EtOH and ethyl acetate (4:1) at 50 °C provided **25a** in 80% yield. Yield of the product **25a** was low when the reaction was conducted without hydrogen at rt (10%) or at 50 °C (25%). The reactant **16h** was not readily soluble in EtOH, hence, we used the solvent mixture of EtOH and EtOAc (4:1). Even in this solvent mixture the reaction was sluggish at rt (50% yield after 24 h). Structure of the product was confirmed based on spectroscopic and analytical data. The reaction was generalized by subjecting two more thio-ethers **16i** and **16p** for reductive desulfurization to furnish the pyrazolo[3,4-*b*]pyridines **25b-c**.

**Scheme 6.**
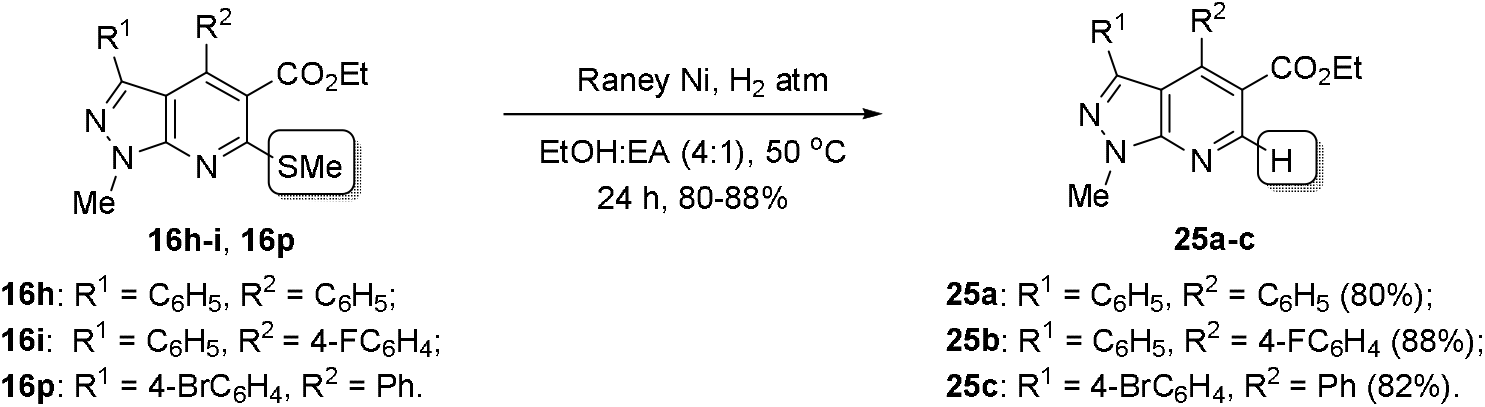
Reductive desulfurization of **16h-i**, **16p** to furnish pyrazolo[3,4-*b*]pyridines **25a-c**.

Next, we conducted hydrolysis of the ester in **16h** to provide the carboxylic acid **26a** (Scheme 7). The acid functional group was intended to increase hydrophlicity of the product and for preparation of different amides / esters. We tried several conditions, among them, the hydrolysis was facile with LiOH (3 equivalents) in a mixture of 1,4-dioxane and water (4:1) worked to provide the acid **26a** in 80% yield.^35^ With LiOH in MeOH or KOH in MeOH the reaction was not clean, and any tangible product did not form. We generalized the reaction of the ester hydrolysis by subjecting three more pyrazolopyridines **16i-k** to furnish the carboxylic acids **26b-d** in good yield.

**Scheme 7.**
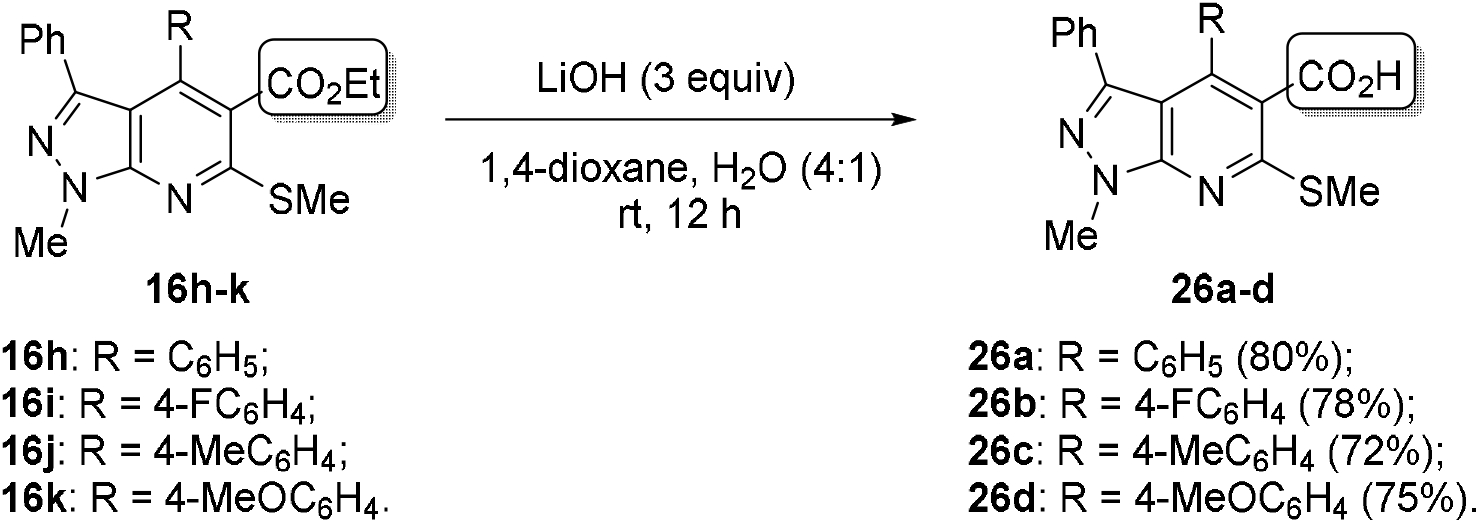
Hydrolysis of **16h-k** to furnish pyrazolo[3,4-*b*]pyridines **26a-d**.

Finally, we planned a palladium mediated Suzuki coupling of pyrazolopyridine **16p**, which has 4-bromophenyl substitution with phenylboronic acid to prepare pyazolopyridines **27** to demonstrate the proof-of-the-principle (Scheme 8). With additional aromatic ring in place, **27** is expected to display increased hydrophobic / lipophilic characteristics. The Suzuki coupling worked best with Pd(OAc)_2_ (10 mol%, catalyst), PPh_3_ (3 equivalents, ligand) and K_3_PO_4_ (3 equivalents, base), in dichloroethane (DCE, solvent) reflux. Before arriving at the optimal conditions, we tried Pd(PPh_3_)_4_ (10 mol% or 30 mole%) as the catalyst, toluene or dioxane as a solvent but the reaction did not take place. With Pd(OAc)_2_ (10 mol%) and PPh_3_ (3 equivalents) K_2_CO_3_ (3 equivalents, base) in 1,4-dioxane as solvent provided the product **27** in only 20% yield. Switching to K_3_PO_4_ as the base and 1,4-dioxane (reflux) as solvent improved the yield to 50%.

**Scheme 8.**
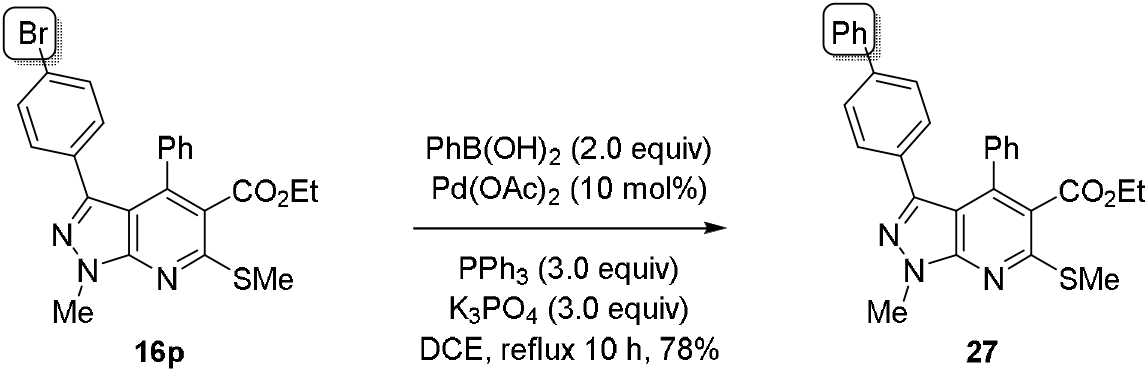
Suzuki Coupling of aryl boronic acids and aryl bromides **16p** to furnish pyrazolo[3,4-*b*]pyridine **27**.

### 2.3. In silico screening, in-vitro MTB MABA assay and molecular docking studies

Initially, newly prepared the pyrazolo[3,4-b]pyridines were subjected to *in silico* screening for drug likeliness, such as adherence to Lipinski’s Rule of Five and for Absorption, Distribution, Metabolism, Excretion and Toxicity (ADMET) properties. The analysis showed that all of them obeyed Lipinski’s Rule of Five and possess reasonable ADMET properties. The predicted values for CNS penetration, number of HB acceptors, number of HB donors and solubility were found to be within the limits, which infer that the compounds have good transport and bioavailability, following which we evaluated the NCEs for anti-tuberculotic activity.^36^

We selected thirteen best pyrazolo[3,4-*b*]pyridines from the pool for MABA assay against *M. tuberclulosis* H37Rv. Apart from adherence to the Lipinski’s Rule of Five and favourable ADMET properties, the screening criteria included structural diversity. Out of those subjected to the MABA assay, the pyrazolopyridines, **16k**, **16w and 17b** were the best to inhibit the growth of *M. tuberculosis* at IC_50_ of 3.125 µg/mL. This value compared favourably with that of isoniazid (IC_50_ = 0.36 µg/mL).

Based on the results obtained from *in vitro* studies (Tables 2-3) and *in silico* screening of drug likeliness (Supplementary Data, Tables 1-2), we selected the **16k** (both the regioisomers) for molecular docking studies with MTBPS (PDB ID: 1MOP). The results (Figures 5-6) showed that **16kA** and **16kB** (isomers **A** and **B** of **16k**) exhibit significant binding affinity (−10.15 kcal/mol (Ki: 36.29 nM) and −8.70 kcal/mol (Ki: 419.5 nM), which are appreciable and comparable with the best compound of the previously reported studies of SOMs from taken from chemical libraries.^37^ The docking results (Figures 3-6) corroborate with *in vitro* MIC assay. The residue contribution analysis explained that **16kA** fits better in the active site of MTBPS compared to its isomer **16kB**. In the active site the pyrazolo[3,4-*b*]pyridine **16kA** exhibited three intermolecular hydrogen bonding interactions with His44, Gly158 and Gln164, with the bond distance of 2.93 Å, 2.53 Å and 3.13 Å, respectively. In addition, the ligand also displayed hydrophobic interactions with the residues of Met195, Lys160, Gly146, Asp161, Phe157, Val142, Gln72, Thr39, Pro38, Met40, Ser197, Tyr82 and His47. Further, the electrostatic surface potential map of protein-ligand complexes, computed using APBS plugin of PyMOL, exhibited that the isomer **16kA** oriented towards positively charged amino acid residues in the active site of MTBPS. Like **16kA** its isomer **16kB**, also displayed hydrogen bonding with Ser197, Lys160, Glu72 located around the active site with the bond distances of 2.87 Å, 2.88 Å and 2.86 Å respectively. Moreover, it also showed hydrophobic interactions with the Arg198, Ser196, Gln164, Asp161, Phe157, Phe156, Tyr82, Leu50, His47, Gly41, Met40, Thr39, and Pro38 residues. The residues Gln164, Lys160, Asp161, Phe157, Glu72, and Thr39 were common for both the isomers.

**Figure 5.**
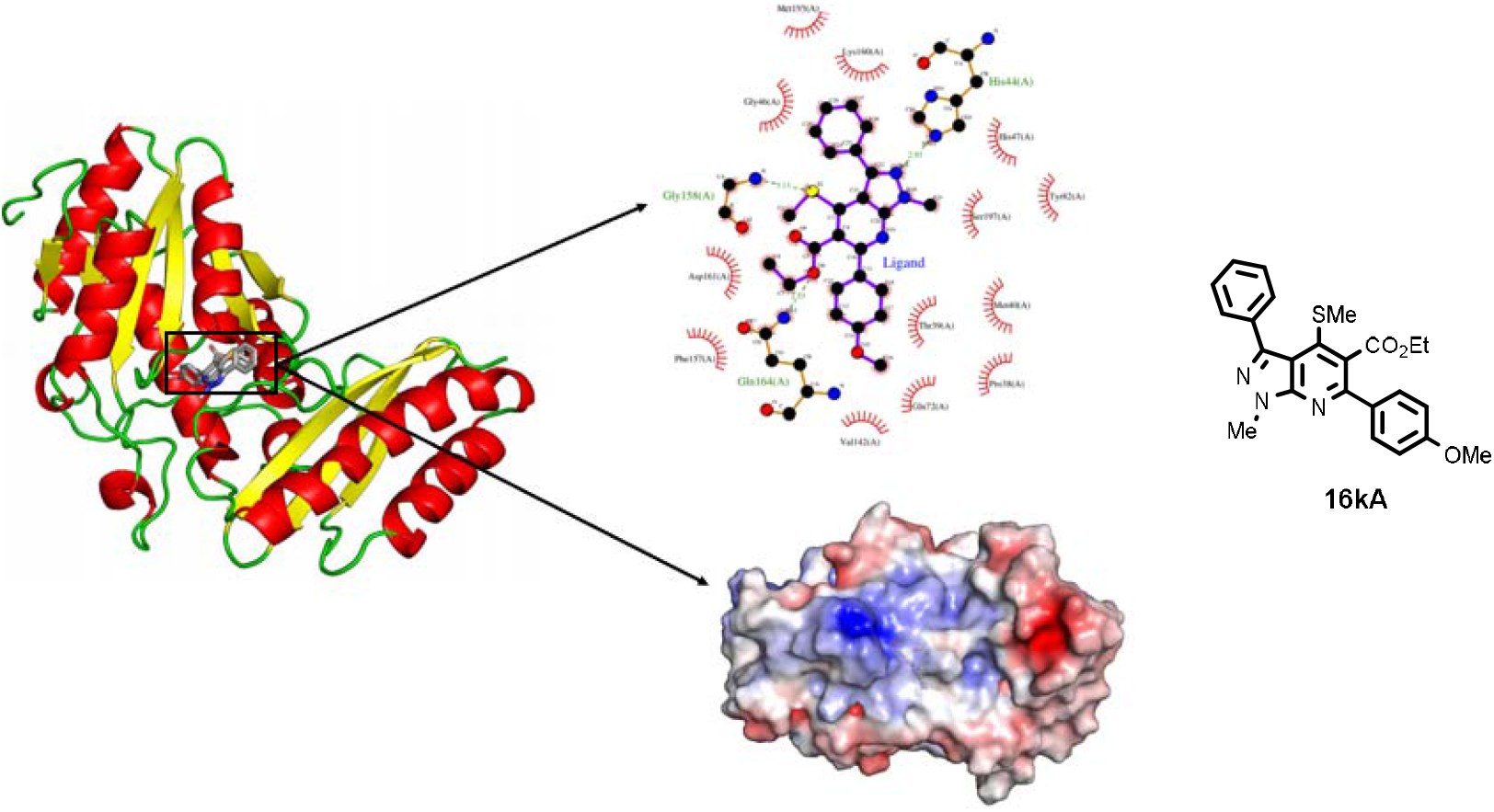
Molecular docking results of the pyrazolo[3,4-b]pyridine **16kA** (isomer A with C(4)SMe) with MTBPS.

**Figure 6.**
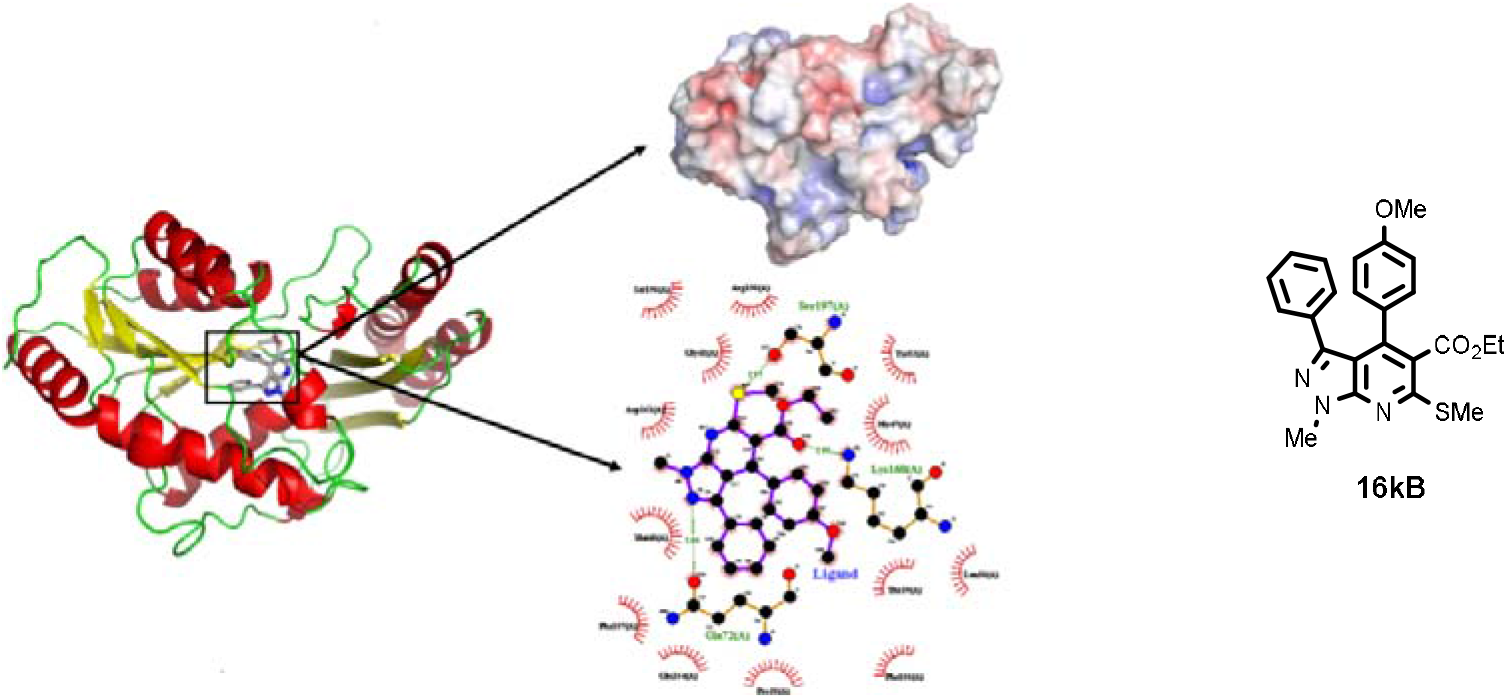
Molecular docking results of the pyrazolo[3,4-b]pyridine **16kB** (regioisomer **B** with C(6)SMe) with MTBPS.

**Table 2.**
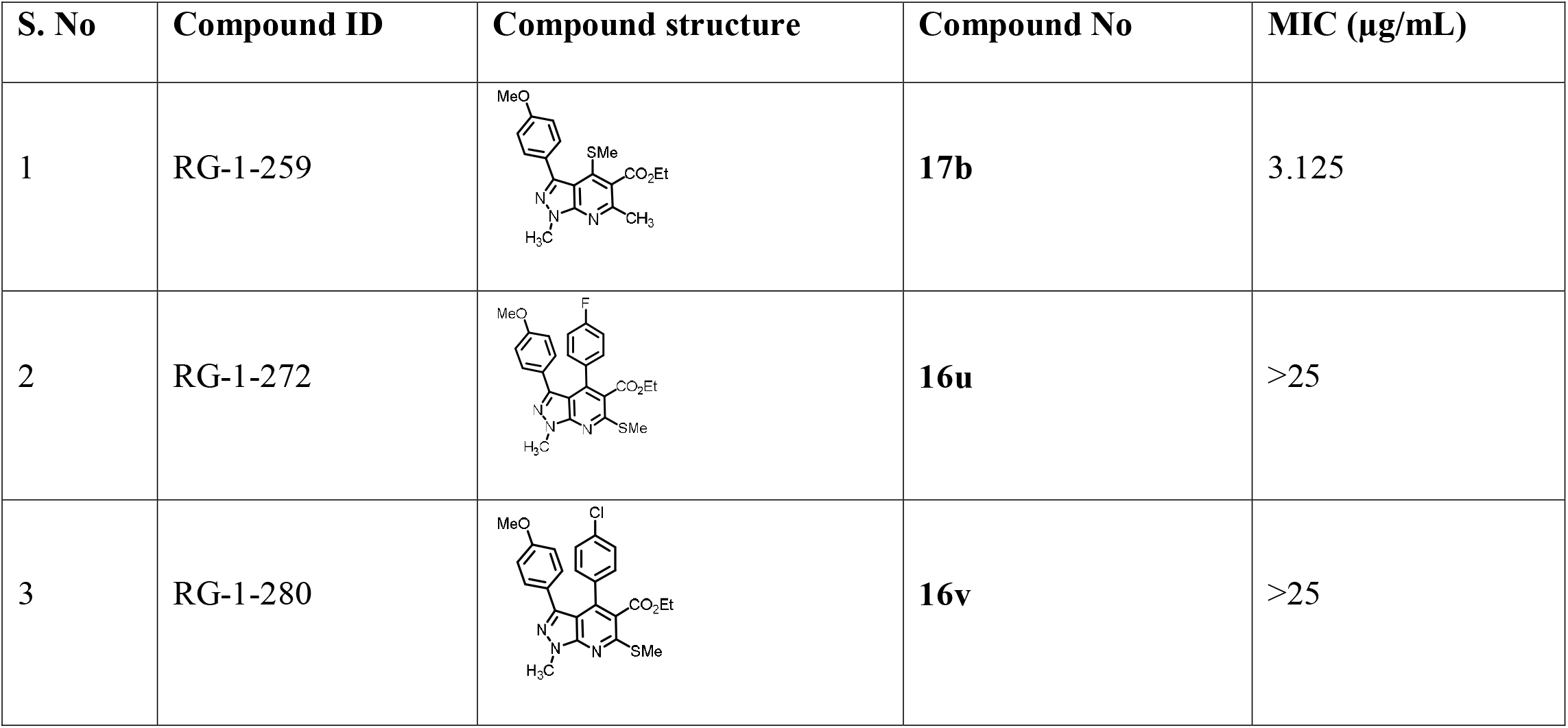
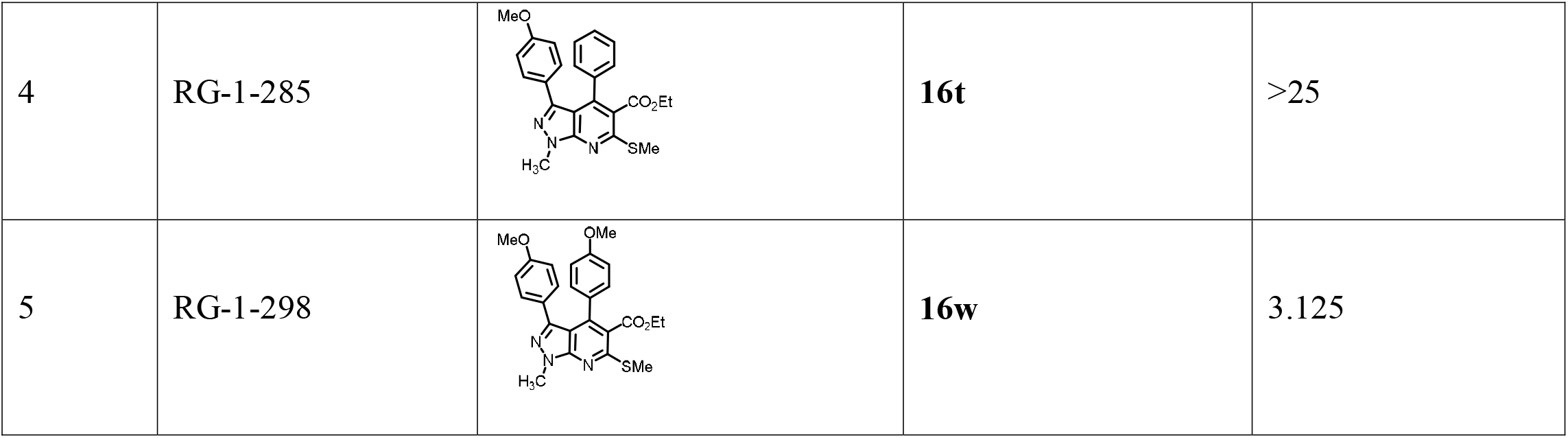
MABA assay of pyrazolo[3,4-*b*]pyridines against *M. tuberclulosis* H37Rv.

| S. No | Compound ID | Compound structure | Compound No | MIC (µg/mL) |
| --- | --- | --- | --- | --- |
| 1 | RG-1-259 |  | **17b** | 3.125 |
| 2 | RG-1-272 |  | **16u** | >25 |
| 3 | RG-1-280 |  | **16v** | >25 |
| 4 | RG-1-285 |  | **16t** | >25 |
| 5 | RG-1-298 |  | **16w** | 3.125 |

**Table 3.**
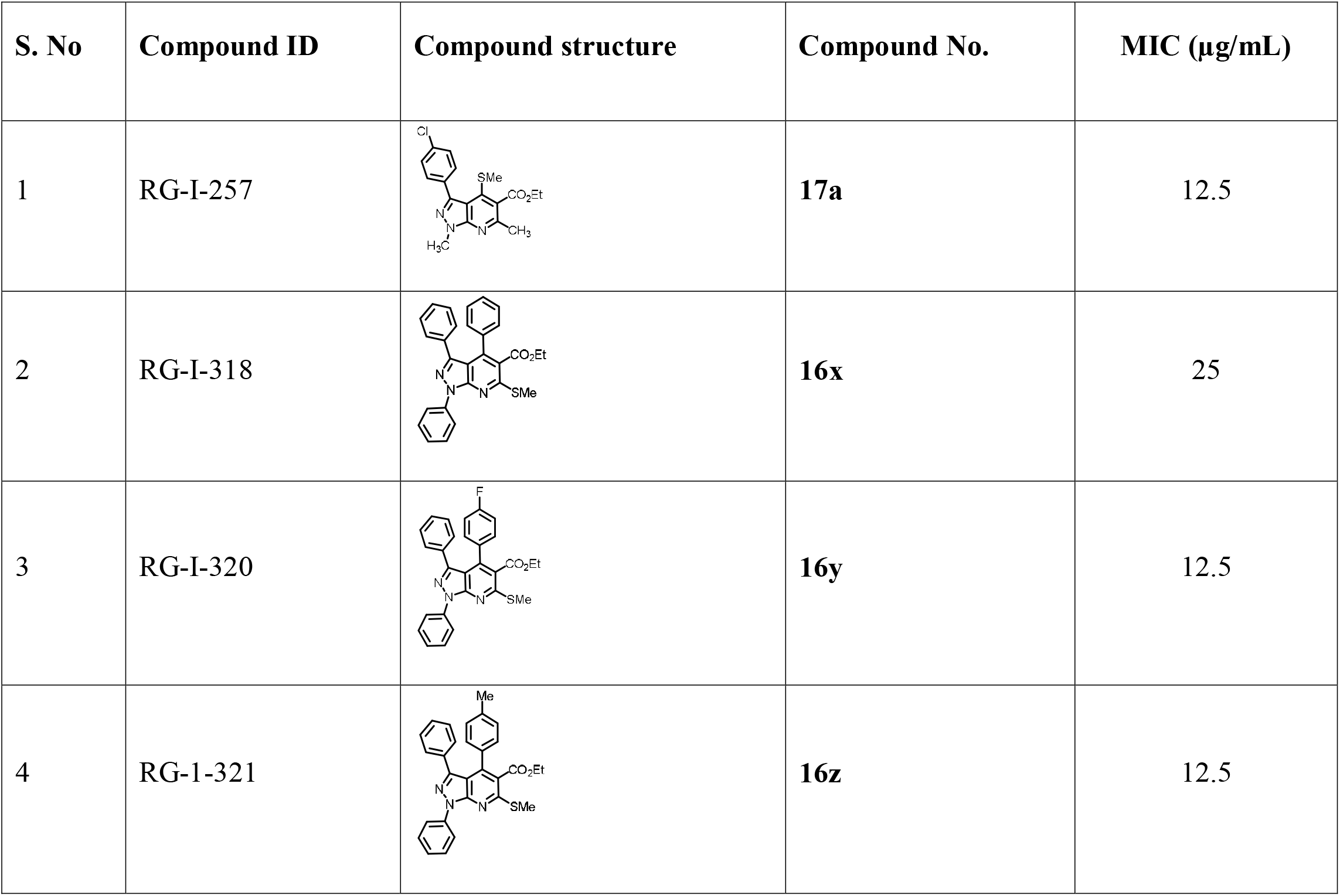
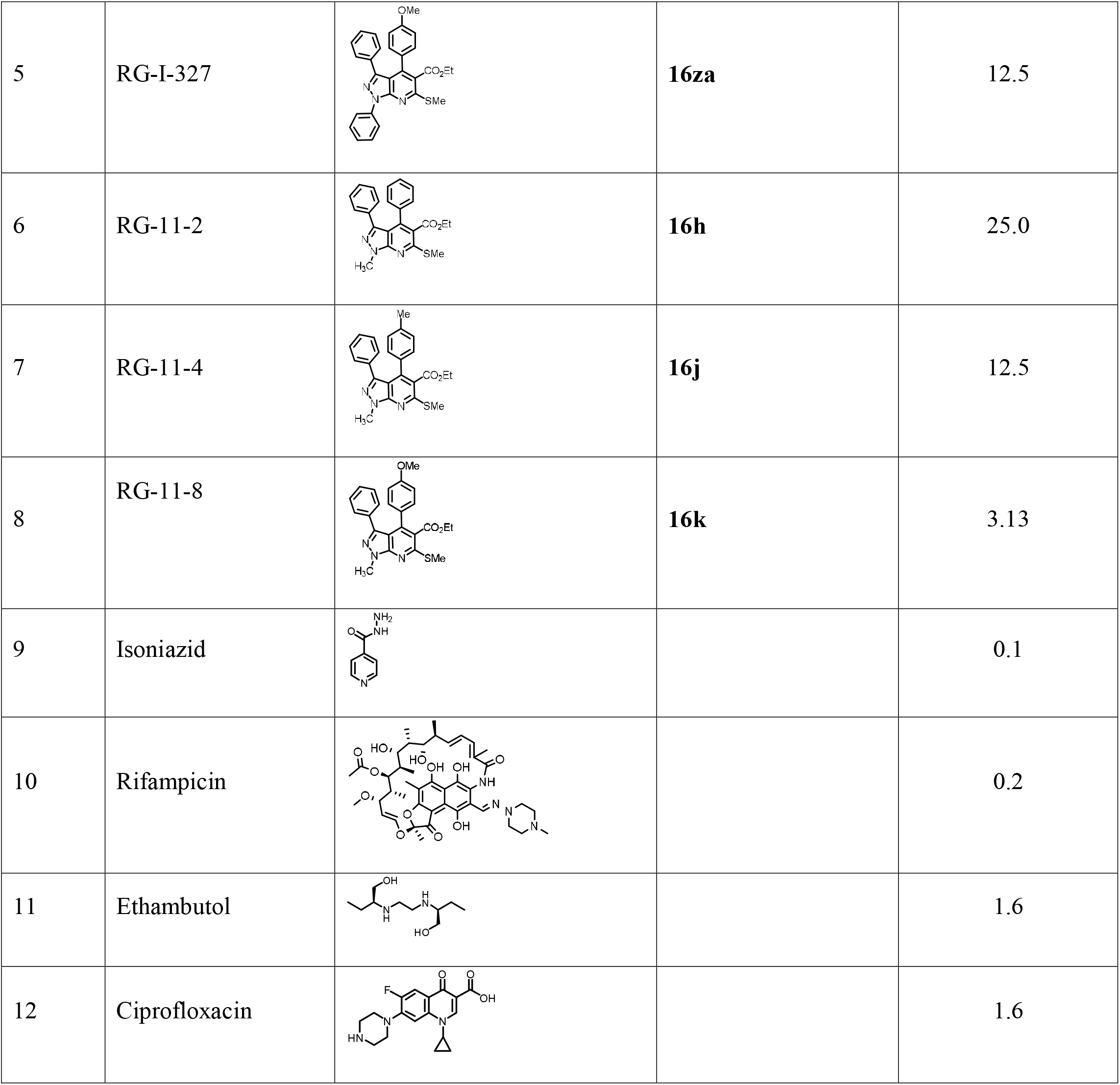
MABA assay of pyrazolo[3,4-*b*]pyridines against *M. tuberclulosis* H37Rv.

| S. No | Compound ID | Compound structure | Compound No. | MIC (µg/mL) |
| --- | --- | --- | --- | --- |
| 1 | RG-I-257 |  | **17a** | 12.5 |
| 2 | RG-I-318 |  | **16x** | 25 |
| 3 | RG-I-320 |  | **16y** | 12.5 |
| 4 | RG-1-321 |  | **16z** | 12.5 |
| 5 | RG-I-327 |  | **16za** | 12.5 |
| 6 | RG-11-2 |  | **16h** | 25.0 |
| 7 | RG-11-4 |  | **16j** | 12.5 |
| 8 | RG-11-8 |  | **16k** | 3.13 |
| 9 | Isoniazid |  |  | 0.1 |
| 10 | Rifampicin |  |  | 0.2 |
| 11 | Ethambutol |  |  | 1.6 |
| 12 | Ciprofloxacin |  |  | 1.6 |

## 3. Conclusion

In conclusion, we have demonstrated a facile synthesis of a combinatorial library of pyrazolo[3,4-*b*]pyridines **16** and in some cases **17** by the condensation of 5-aminopyrazoles **7** and OKDTAs **15**. In contrast to the reaction of **7** with OKDTAs without C(2)-ester, in which a mixture of two regio-isomeric pyrazolo[3,4-*b*]pyridines namely **16** (major) and **17** (minor) formed, the reaction with C(2) ester furnished the regio-isomer **16** exclusively. We have achieved reductive desulfurization, hydrolysis of the ester and Suzuki coupling with a few aryl bornoic acids on **16**. *In vitro* MABA assay against *M. tuberclulosis* H37Rv^38^ indicated that **16k** displayed best activity. *In silico* analysis of binding of **16k** with the bacterial MTBPS corroborated the *in vitro* results. We believe that our results will pave way for the discovery of a pyazolo[3,4-*b*]pyridine based anti-TB drug.

## 4. Experimental section

### 4.1. General

The solvents were purified and dried using conventional methods.^39^ Thin layer chromatography (TLC; SiO_2_) was used to monitor all the reactions. Column chromatography (CC) on silica gel (100-200 mesh, AVRA Synthesis Private Limited) was performed with an increasing amount of ethyl acetate (EtOAc) in hexanes. The melting points were determined using open-ended capillary tubes on BUCHI M-560 equipment. The IR spectra were recorded as KBr pellets on a Nicolet-6700 spectrometer. The ^1^H NMR (400 MHz), ^13^C NMR (100 MHz), DEPT-135, HMBC, and HSQC 2D NMR spectra were acquired on a Bruker - Avance 400 MHz spectrometer using tetramethylsilane (TMS) as the internal standard for (CDCl_3_ or DMSO-*d_6_*) solutions; *J*-values are given in Hz. Broad band ^1^H decoupling was used to record the ^13^C NMR spectra. The DEPT-135 NMR spectra were recorded for each sample to assign if the carbons were CH_3_/CH or CH_2_ or C (absence in the spectrum by matching with ^13^C NMR spectrum) in support the given structure. High-resolution mass spectra were obtained in electrospray ionization mode on a Water Q-TOF micro mass spectrometer and an Agilent 6350 B Q-TOF mass spectrometer. Reagents, catalysts, acids, and bases were procured from commercial sources and used as such. The pyrazol-5-amines and OKDTAs were synthesized as per the literature procedures as indicated in the text.

### 4.2. Synthesis of pyrazolo[3,4-b]pyridines: General procedure

Synthesis of 6-(methylthio)-1,3,4-triphenyl-1*H*-pyrazolo[3,4-*b*]pyridine **16a** and 4-(methylthio)-1,3,6-triphenyl-1*H*-pyrazolo[3,4-*b*]pyridine **17a**.

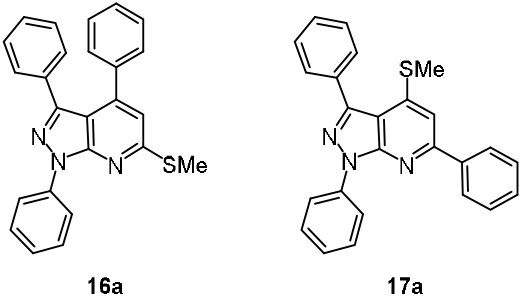

A solution of 1,3-diphenyl-1*H*-pyrazol-5-amine (224 mg, 1.00 mmol) **7a** 3,3-bis(methylthio)-1-phenylprop-2-en-1-one **15a** (224 mg, 1.00 mmol) and trifluoroacetic acid (TFA) (36 mg, 0.03 mmol) in anhydrous toluene (6 mL) was heated to reflux for 6 h by which time both the amine and OKDTA were absent (TLC, 2% EtOAc in hexanes). The acid in the reaction mixture after cooling was quenched with Na_2_CO_3_ and charged on a silica gel column and eluted with 1% ethyl acetate in hexanes. Pooling of fractions having **16a**/**17a** and evaporation of the solvent under reduced pressure provided pure products.

6-(Methylthio)-1,3,4-triphenyl-1*H-*pyrazolo[3,4-*b*]pyridine **16a**, white solid. Yield: 275 mg (62%). MP: 161°C. IR (KBr, cm^-1^) 3058, 2923, 1595, 1563, 1303, 1095, 756. ^1^H NMR (400 MHz, CDCl_3_ + CCl_4_ (1:1)) δ 8.50 - 8.35 (d, 2H), 7.50 (t, *J* = 8.0 Hz, 2H), 7.32 - 7.21 (m, 2H), 7.19 - 7.09 (m, 8H), 7.09 - 7.04 (m, 2H), 2.73 (s, 3H) ppm. ^13^C NMR (100 MHz, CDCl_3_ + CCl_4_ (1:1)) δ 160.7 (C), 151.6 (C), 146.2 (C), 145.4 (C), 139.7 (C), 136.8 (C), 133.0 (C), 129.5 (CH), 129.2 (CH), 129.0 (CH), 128.5 (CH), 128.0 (CH), 127.8 (CH), 127.6 (CH), 125.9 (CH), 121.4 (CH), 117.1 (CH), 110.2 (C), 13.5 (CH_3_) ppm. HRMS (ESI+) Calcd for C_25_H_19_N_3_S [M+H]^+^ 394.1372, found 394.1410.

4-(Methylthio)-1,3,6-triphenyl-1*H*-pyrazolo[3,4-*b*]pyridine **17a**, white solid. Yield: 52 mg (23%). MP: 168 °C. IR (KBr, cm^-1^) 3055, 2920, 1596, 1560, 1542, 1497, 1359, 974, 748. ^1^H NMR (400 MHz, CDCl_3_ + CCl_4_ (1:1)) δ 8.45 (dd, *J* = 8.7, 1.1 Hz, 2H), 8.11 (dd, *J* = 8.2, 1.3 Hz, 2H), 7.77 (ddd, *J* = 5.3, 2.9, 1.5 Hz, 2H), 7.56 – 7.38 (m, 8H), 7.34 - 7.18 (m, 2H), 2.53 (s, 3H) ppm. ^13^C NMR (100 MHz, CDCl_3_ + CCl_4_ (1:1)) δ 156.6 (C), 151.0 (C), 147.0 (C), 146.0 (C), 139.9 (C), 139.5 (C), 133.5 (CH), 130.5 (CH), 129.6 (CH), 129.0 (CH), 128.99 (CH), 128.96 (CH), 128.1 (CH), 127.8 (CH), 125.9 (CH), 121.6 120 (CH), 112.6 (C), 109.0 (CH), 14.3 (CH_3_) ppm. HRMS (ESI+) Calcd for C_25_H_19_N_3_S [M+H]^+^ 394.1372 amu, found 394.1410 amu.

Synthesis of 6-(methylthio)-1,4-diphenyl-3-(thiophen-2-yl)-1*H*-pyrazolo[3,4-*b*]pyridine **16b**, and 4-(methylthio)-1,6-diphenyl-3-(thiophen-2-yl)-1*H*-pyrazolo[3,4-*b*]pyridine **17b**.

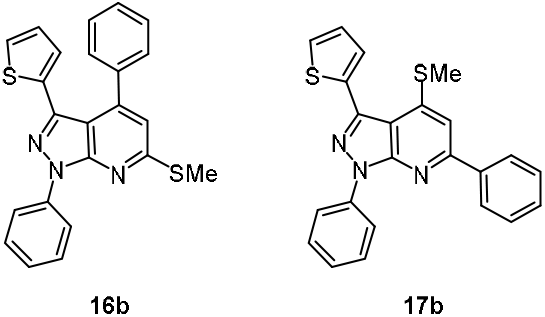

The condensation reaction of 1-phenyl-3-(thiophen-2-yl)-1*H*-pyrazol-5-amine **7b** (241 mg, 0.10 mmol), 3,3-bis(methylthio)-1-phenylprop-2-en-1-one **15a** (224 mg, 0.10 mmol) and TFA (42 mg, 0.03 mmol) in toluene (3 mL) reflux for 7 h by following the general procedure provided **16b** and **17b** in combined yield of 77%. 6-(Methylthio)-1,4-diphenyl-3-(thiophen-2-yl)-1*H*-pyrazolo[3,4-*b*]pyridine **16b**, white solid. Yield: 209 mg (52%). MP: 146 °C. IR (KBr, cm^-1^) 3053, 2917, 1631, 1539, 1349, 1213, 765. ^1^H NMR (400 MHz, CDCl_3_ + CCl_4_ (1:1)) δ 8.40 (dd, *J* = 8.7, 1.1 Hz, 2H), 7.50 (dd, *J* = 8.5, 7.5 Hz, 2H), 7.39 - 7.25 (m, 6H), 7.17 (dd, *J* = 5.1, 1.1 Hz, 1H), 7.03 (s, 1H), 6.67 (dd, *J* = 5.1, 3.6 Hz, 1H), 6.17 (dd, *J* = 3.6, 1.1 Hz, 1H), 2.71 (d, *J* = 3.2 Hz, 3H) ppm. ^13^C NMR (100 MHz, CDCl_3_ + CCl_4_ (1:1)) δ 160.9 (C), 151.6 (C), 145.4 (C), 140.1 (C), 139.6 (C), 137.3 (C), 134.5 (C), 129.2 (C), 129.0 (CH), 128.7 (CH), 128.7 (CH), 128.2 (CH), 126.8 (CH), 126.1 (CH), 126.0 (CH), 121.5 (CH), 117.6 (CH), 110.2 (C), 13.6 (CH_3_) ppm. HRMS (ESI+) Calcd for C_23_H_17_N_3_S [M+H]^+^ 400.0864, found 400.0947.

4-(Methylthio)-1,6-diphenyl-3-(thiophen-2-yl)-1*H*-pyrazolo[3,4-*b*]pyridine **17b**, white solid. Yield: 101 mg (25%). MP: 146 °C. IR (KBr, cm^-1^) 3055, 2921, 1552, 1521, 1493, 1414, 1355, 1298, 758. ^1^H NMR (400 MHz, CDCl3) δ 8.36 (dd, *J* = 8.6, 1.0 Hz, 2H), 8.07 - 8.00 (m, 2H), 7.55 (dd, *J* = 3.6, 1.1 Hz, 1H), 7.50 – 7.38 (m, 6H), 7.28 - 7.18 (m, 2H), 7.11 (dd, *J* = 5.1, 3.6 Hz, 1H), 2.53 (s, 3H) ppm. ^13^C NMR (100 MHz, CDCl_3_) δ 156.7 (C), 150.9 (C), 147.0 (C), 139.6 (C), 139.5 (C), 139.3 (C), 134.2 (C), 129.9 (C), 129.7 (CH), 129.0 (CH), 128.9 (CH), 127.8 (CH), 127.3, 127.2 (CH), 126.1 (CH), 121.7 (CH), 112.6 (C), 109.2 (CH), 14.3 (CH_3_) ppm. HRMS (ESI+) Calcd for C_23_H_17_N_3_S [M+H]^+^ 400.0864, found 400.0954.

Synthesis of 6-(methylthio)-4-(naphthalen-1-yl)-1-phenyl-3-(thiophen-2-yl)-1*H*-pyrazolo[3,4-*b*]pyridine **16c** and 4-(methylthio)-6-(naphthalen-1-yl)-1-phenyl-3-(thiophen-2-yl)-1*H*-pyrazolo[3,4-*b*]pyridine **17c**.

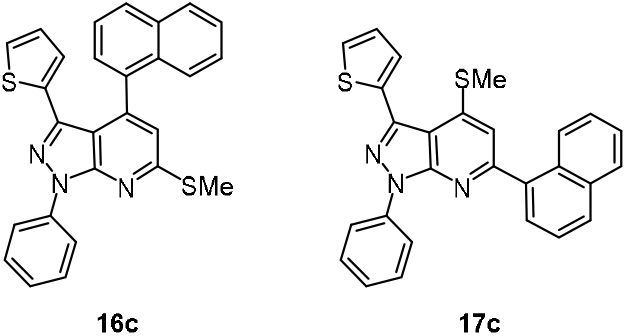

The condensation of 1-phenyl-3-(thiophen-2-yl)-1*H*-pyrazol-5-amine **7b** (241 mg, 0.10 mmol) 3,3-bis(methylthio)-1-(naphthalen-1-yl)prop-2-en-1-one **15b** (274 mg, 0.10 mmol) and trifluoroacetic acid (TFA) (42 mg, 0.03 mmol) in toluene (5 mL) reflux for 6 h by following the general procedure provided **16c** and **17c** in combined yield of 77%.

6-(Methylthio)-4-(naphthalen-1-yl)-1-phenyl-3-(thiophen-2-yl)-1*H-*pyrazolo[3,4-*b*]pyridine **16c**, white solid. Yield: 301 mg (67%). MP: 214 °C. IR (KBr, cm^-1^) 2922, 2856, 1629, 1558, 1495, 1413, 1365, 1299, 936, 771. ^1^H NMR (400 MHz, CDCl_3_) δ 8.44 (d, *J* = 7.9 Hz, 2H), 7.86 (d, *J* = 8.1 Hz, 1H), 7.79 (d, *J* = 8.2 Hz, 1H), 7.52 (t, *J* = 8.0 Hz, 3H), 7.45 - 7.21 (m, 6H), 7.13 (s, 1H), 6.89 (d, *J* = 4.9 Hz, 1H), 6.33 - 6.25 (m, 1H), 5.69 (d, *J* = 3.1 Hz, 1H), 2.74 (s, 3H) ppm. ^13^C NMR (100 MHz, CDCl_3_) δ 161.0 (C), 151.4 (C), 143.7 (C), 140.4 (C), 139.7 (C), 135.4 (C), 134.0 (C), 133.3 (C), 132.0 (C), 129.1 (CH), 129.0 (CH), 128.1 (CH), 127.4 (CH), 127.2 (CH), 126.7 (CH), 126.5 (CH), 126.3 (CH), 126.0 (C), 125.7 (CH), 125.5 (CH), 125.1 (CH), 121.4 (CH), 118.8 (CH), 111.7 (C), 13.6 (CH_3_) ppm. HRMS (ESI+) Calcd for C_27_H_19_N_3_S_2_ [M +H]^+^ 450.1020, found 450.1102.

4-(Methylthio)-6-(naphthalen-1-yl)-1-phenyl-3-(thiophen-2-yl)-1*H* pyrazolo[3,4-*b*]pyridine **17c**, white solid. Yield: 63 mg (14%). MP: 208 °C. IR (KBr, cm^-1^) 3053, 2917, 1539, 1497, 1407, 1349, 1213, 930, 765. ^1^H NMR (400 MHz, CDCl_3_) δ 8.41 (dd, *J* = 8.7, 1.1 Hz, 2H), 8.32 – 8.26 (m, 1H), 7.97 (t, *J* = 8.0 Hz, 2H), 7.75 (dd, *J* = 7.1, 1.1 Hz, 1H), 7.69 (dd, *J* = 3.6, 1.1 Hz, 1H), 7.60 (dd, *J* = 8.2, 7.2 Hz, 1H), 7.53 (ddd, *J* = 7.7, 5.0, 1.4 Hz, 3H), 7.49 - 7.43 (m, 2H), 7.30 - 7.21 (m, 4H), 2.53 (s, 3H) ppm. ^13^C NMR (100 MHz, CDCl_3_) δ 158.5 (C), 150.6 (C), 146.8 (C), 139.59 (C), 139.52 (C), 138.5 (C), 134.0 (C), 134.0 (C), 131.4 (C), 130.0 (CH), 129.5 (CH), 129.1 (CH), 128.6 (CH), 128.0 (CH), 127.4 (CH), 127.3 (CH), 126.7 (CH), 126.1 (CH), 125.8 (CH), 125.4 (CH), 121.6 (CH), 113.8 (CH), 112.4 (C), 14.3 (CH_3_) ppm. HRMS (ESI+) Calcd for C_27_H_19_N_3_S_2_ [M+H]^+^ 450.1020, found 450.1097.

Synthesis of 3-methyl-6-(methylthio)-1,4-diphenyl-1*H*-pyrazolo[3,4-*b*]pyridine **16d** and 3-methyl-4-(methylthio)-1,6-diphenyl-1*H*-pyrazolo[3,4-*b*]pyridine **17d**.

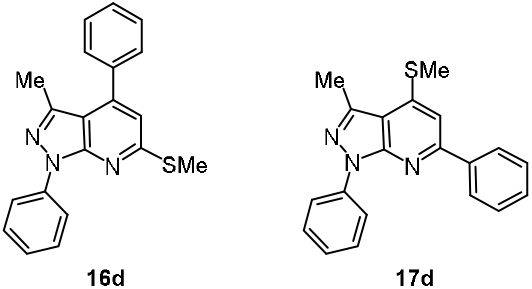

The condensation of 3-methyl-1-phenyl-1*H*-pyrazol-5-amine (86 mg, 0.50 mmol) **7c**, 3,3-bis(methylthio)-1-phenylprop-2-en-1-one (126 mg, 0.55 mmol) **15a**, and TFA (16 mg, 0.015 mmol) in anhydrous toluene (3 mL) reflux for 5 h **16d** and 3-methyl-4-(methylthio)-1,6-diphenyl-1*H*-pyrazolo[3,4-*b*]pyridine **17d** by following the general procedure in combined yield of 88%.

3-Methyl-6-(methylthio)-1,4-diphenyl-1*H*-pyrazolo[3,4-*b*]pyridine **16d**, white solid. Yield: 106 mg (63%), MP: 126 °C. IR (KBr, cm^-1^) 2942, 1704, 1624, 1380, 1265, 1093, 765. ^1^H NMR (400 MHz, CDCl_3_ + CCl_4_ (1:1)) δ 8.37 - 8.24 (m, 2H), 7.51 - 7.40 (m, 7H), 7.24 (d, *J* = 1.7 Hz, 1H), 6.91 (s, 1H), 2.69 (s, 3H), 2.22 (s, 3H) ppm. ^13^C NMR (100 MHz, CDCl_3_ + CCl_4_ (1:1)) δ 160.4 (C), 151.4 (C), 145.4 (C), 142.6 (C), 139.9 (C), 137.5 (C), 129.1 (CH), 128.9 (CH), 128.8 (CH), 128.4 (CH), 125.4 (CH), 120.8 (CH), 116.4 (CH), 111.9 (C), 15.5 (CH3), 13.5 (CH_3_) ppm. HRMS (ESI+) Calcd for C_20_H_18_N_3_S [M+H]^+^ 332.1216 amu, found 332.1256 amu.

3-Methyl-4-(methylthio)-1,6-diphenyl-1*H*-pyrazolo[3,4-*b*]pyridine **17d**, white solid. Yield: 30 mg (25%). MP: 108 °C. IR (KBr, cm^-1^) 2965, 2922, 2853, 1642, 1580, 1265, 1020, 742. ^1^H NMR (400 MHz, CDCl_3_ + CCl_4_ (1:1)) δ 8.36 (d, *J* = 7.7 Hz, 2H), 8.13 - 8.01 (m, 2H), 7.48 (t, *J* = 8.0 Hz, 5H), 7.23 (d, *J* = 12.4 Hz, 3H), 2.79 (s, 3H), 2.66 (s, 3H) ppm. ^13^C NMR (100 MHz, CDCl_3_+CCl_4_ (1:1)) δ 156.8 (C), 151.0 (C), 146.8 (C), 142.7 (C), 140.0 (C), 139.7 (C), 129.5 (CH), 129.0 (CH), 128.9 (CH), 127.8 (CH), 125.4 (CH), 121.1 (CH), 113.7 (CH), 108.0 (C), 15.8 (CH3), 13.9 (CH_3_) ppm. HRMS (ESI+) Calcd for C_20_H_18_N_3_S [M+H]^+^ 332.1216, found 332.1256.

Synthesis of 1-methyl-6-(methylthio)-3,4-diphenyl-1*H*-pyrazolo[3,4-*b*]pyridine **16e** and 1-methyl-4-(methylthio)-3,6-diphenyl-1*H*-pyrazolo[3,4-*b*]pyridine **17e**.

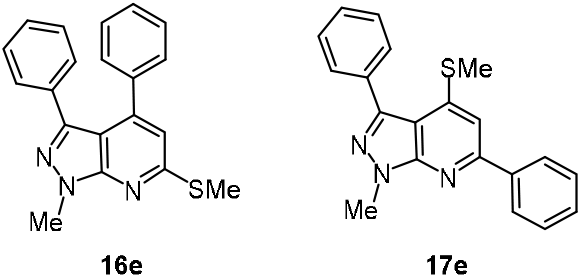

The condensation reaction of 1-methyl-3-phenyl-1*H*-pyrazol-5-amine (173 mg, 1.00 mmol) **7d**, 3,3-bis(methylthio)-1-phenylprop-2-en-1-one (224 mg, 1.00 mmol) **15a**, and TFA (36 mg, 0.03 mmol) in anhydrous in toluene (5 mL) reflux for 7 hr by following general procedure provided **16e** and **17e** in 83% yield. 1-Methyl-6-(methylthio)-3,4-diphenyl-1*H*-pyrazolo[3,4-*b*]pyridine **16e**, white solid. Yield: 238 mg (71%), MP: 120 °C. IR (KBr, cm^-1^) 3062, 2924, 2855, 1561, 1500, 1308, 1064, 756. ^1^H NMR (400 MHz, CDCl_3_ + CCl_4_ (1:1)) δ 7.18 (s, 1H), 7.07 (dd, *J* = 9.3, 7.1 Hz, 5H), 6.99 (dd, *J* = 5.9, 4.6 Hz, 4H), 6.88 (s, 1H), 4.13 (s, 3H), 2.64 (s, 3H) ppm. ^13^C NMR (100 MHz, CDCl_3_ + CCl4 (1:1) δ 160.1 (C), 152.2 (C), 145.2 (C), 144.5 (C), 137.3 (C), 133.6 (C), 129.4 (CH), 129.3 (CH), 128.4 (CH), 128.1 (CH), 127.6 (CH), 127.5 (CH), 116.3 (CH), 108.5 (C), 34.3 (CH3), 13.3 (CH_3_) ppm. HRMS (ESI+) Calcd for C_20_H_18_N_3_S [M + H]^+^ 332.1216, found 332.1256. 1-Methyl-4-(methylthio)-3,6-diphenyl-1*H*-pyrazolo[3,4-*b*]pyridine **17e**, white solid. Yield: 24 mg (12%). MP: 86 °C. IR (KBr, cm^-1^) 2927, 1732, 1556, 1478, 1349, 1223, 766. ^1^H NMR (400 MHz, CDCl_3_ + CCl_4_ (1:1) δ 8.16 - 8.04 (m, 2H), 7.77 - 7.63 (m, 2H), 7.56 - 7.40 (m, 6H), 7.23 (s, 1H), 4.23 (s, 3H), 2.53 (d, *J* = 2.2 Hz, 3H) ppm. ^13^C NMR (100 MHz, CDCl_3_ + CCl_4_ (1:1)) δ 156.4 (C), 151.2 (C), 146.2 (C), 144.2 (C), 139.7 (C), 133.8 (C), 130.3 (CH), 129.5 (CH), 128.9 (CH), 128.6 (CH), 128.0 (CH), 127.7 (CH),110.7 (CH), 108.3 (C), 34.1 (CH3), 14.2 (CH3) ppm. HRMS (ESI+) Calcd for C_20_H_18_N_3_S [M+H]^+^ 332.1216, found 332.1256.

Synthesis of 6-(methylthio)-4-(naphthalen-1-yl)-1,3-diphenyl-1*H*-pyrazolo[3,4-*b*]pyridine **16f**.

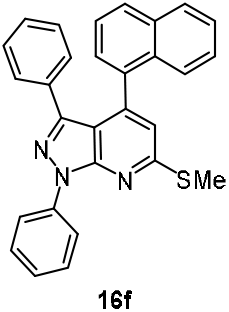

A solution of 1,3-diphenyl-1*H*-pyrazol-5-amine **7e** (118 mg, 0.50 mmol), 3,3-bis(methylthio)-1-(naphthalen-1-yl)prop-2-en-1-one **15a** (118 mg, 0.40 mmol) and TFA (16 mg, 0.015 mmol) in dry toluene (3 mL) reflux by following general procedure provided 170 mg of 6-(methylthio)-4-(naphthalen-1-yl)-1,3-diphenyl-1H-pyrazolo[3,4-b]pyridine **16f** in 77% yield as a white solid. MP: 214 °C. IR (KBr, cm^-1^) 3058, 2924, 1595, 1563, 1499, 1412, 1362, 1302, 1095, 965, 777. ^1^H NMR (400 MHz, CDCl_3_ + CCl_4_ 1:1) δ 8.57 (d, *J* = 8.2 Hz, 2H), 7.83 (dd, *J* = 11.6, 8.2 Hz, 2H), 7.64 (dd, *J* = 13.2, 5.2 Hz, 3H), 7.48 – 7.37 (m, 3H), 7.35 (dd, *J* = 12.3, 3.9 Hz, 3H), 6.97 (dd, *J* = 9.9, 2.8 Hz, 3H), 6.82 (t, *J* = 7.6 Hz, 2H), 2.87 (s, 3H) ppm. ^13^C NMR (100 MHz, CDCl_3_) δ 160.9 (C), 151.5 (C), 146.7 (C), 143.8 (C), 139.9 (C), 135.1 (C),133.3 (C), 132.5 (C), 131.6 (C), 129.0 (C), 128.9 (CH), 128.6 (CH), 128.1 (CH), 127.6 (CH), 127.3 (CH), 127.1 (CH), 126.5 (CH), 126.0 (CH), 125.8 (CH), 125.4 (CH), 124.9 (CH), 121.3 (C), 118.4 (C), 112.2 (C), 13.6 (CH_3_) ppm. HRMS (ESI+) Calcd for C_29_H_21_N_3_S [M+H]^+^ 444.1456, found 444.1560.

Synthesis of 1-ethyl-6-(methylthio)-4-phenyl-1*H*-pyrazolo[3,4-*b*]pyridine **16g**.

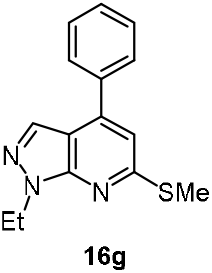

Condensation of 1-ethyl-1*H*-pyrazol-5-amine **7e** (111 mg, 0.10 mmol), 3,3-bis(methylthio)-1-phenylprop-2-en-1-one **15a** (224 mg, 0.10 mmol) and TFA (39 mg, 0.03 mmol) in dry toluene (4 mL) reflux for 6.5 h by following the general procedure provided 1-ethyl-6-(methylthio)-4-phenyl-1*H*-pyrazolo[3,4-*b*]pyridine **16g** as viscous light yellow oil. Yield: 228 mg (85%). IR (KBr, cm^-1^) 2968, 2870, 1660, 1581, 1451, 1372, 1296, 1180, 798. ^1^H NMR (400 MHz, CDCl_3_ + CCl_4_, 1:1) δ 7.98 (s, 1H), 7.77 - 7.56 (m, 2H), 7.53 - 7.34 (m, 3H), 7.01 (s, 1H), 4.55 (q, *J* = 7.3 Hz, 2H), 2.65 (s, 3H), 1.55 (t, *J* = 7.3 Hz, 3H) ppm. ^13^C NMR (100 MHz, CDCl_3_ + CCl_4_, 1:1) δ 159.9 (C), 150.5 (C), 143.1 (C), 137.2 (C), 131.4 (CH), 129.2 (CH), 129.0 (CH), 128.2 (CH), 113.9 (CH), 110.9 (C), 42.0 (CH2), 14.9 (CH3), 13.1 (CH_3_) ppm. HRMS (ESI+) Calcd for C_29_H_21_N_3_S [M +H]^+^ 444.0987, found 444.1076.

Synthesis of ethyl 1-methyl-4-(methylthio)-3,6-diphenyl-1*H*-pyrazolo[3,4-*b*]pyridine-5-carboxylate **16h**.

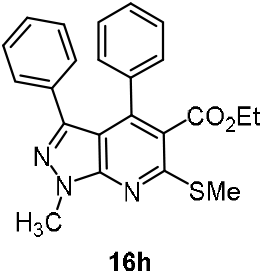

The reaction of 1-methyl-3-phenyl-1H-pyrazol-5-amine **7d** (100 mg, 0.57 mmol) and ethyl 2-benzoyl-3,3-bis(methylthio)acrylate **15c** (170 mg, 0.57 mmol) and a catalytic amount of trifluoroacetic acid (TFA, 19 mg, 30 mol%) in toluene (4 mL) reflux under the blanket of N_2_ atmosphere for 24 h by following the general procedure provided **16h** as white solid. Yield: 190 mg (83%). MP: 140.8 °C. UV (λ_max_, nm) 304 (log ε = 2.491), 217 (log ε = 2.911). IR (KBr, cm^-1^) 3444, 3199, 2959, 2837, 2724, 2578, 1752, 1695, 1601, 1519, 1469, 1397, 1324, 1195, 1127, 945, 840, 735, 676, 508, 405. ^1^H NMR (400 MHz, CDCl_3_) δ 7.22 - 7.18 (m, 1H), 7.11 - 7.00 (m, 5H), 6.98 - 6.93 (m, 4H), 4.19 (s, 3H), 4.02 (q, *J* = 7.2 Hz, 2H), 2.70 (s, 3H), 0.89 (t, *J* = 7.6 Hz, 3H) ppm. ^13^C NMR (100 MHz, CDCl_3_) δ 167.5 (CO), 158.3 (C), 150.9 (C), 145.6 (C), 144.1 (C), 135.7 (C), 132.8 (C), 129.0 (CH), 129.0 (CH), 128.4 (CH), 127.8 (CH), 127.5 (CH), 127.4 (CH), 122.2 (C), 108.7 (C), 61.5 (CH_2_), 33.9 (CH_3_), 14.0 (CH_3_), 13.7 (CH_3_) ppm. HRMS (ESI+): m/z calcd for C_23_H_21_N_3_O_2_S [M+H]^+^ 404.1354, found 404.1439.

Synthesis of ethyl 6-(4-fluorophenyl)-1-methyl-4-(methylthio)-3-phenyl −1*H*-pyrazolo[3,4-*b*]pyridine-5-carboxylate **16i**.

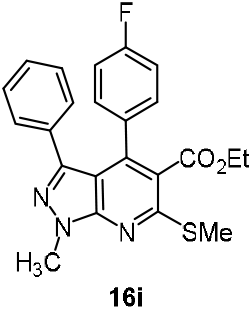

The reaction of 1-methyl-3-phenyl-1*H*-pyrazol-5-amine **7d** (100 mg, 0.57 mmol) and ethyl 2-(4-fluorobenzoyl)-3,3-bis(methylthio)acrylate **15d** (181 mg, 0.57 mmol) and a catalytic amount of trifluoroacetic acid (TFA, 19 mg, 30 mol%) in toluene (4 mL) reflux by following the general procedure given above resulted in **16i**; R_f_ = 0.8. Yield: 195 mg (81%). MP: 137.5 °C. UV (λ_max_, nm) 355.5 (log ε = 1.755), 252 (log ε = 2.425). IR (KBr, cm^-1^) 3444, 3199, 2959, 2837, 2724, 2578, 1752, 1695, 1601, 1519, 1469, 1397, 1324, 1195, 1127, 945, 840, 735, 676, 508, 405. ^1^H NMR (400 MHz, CDCl_3_) δ 7.18 - 7.14 (m, 1H), 7.07 - 7.02 (m, 4H), 6.98 - 6.93 (m, 4H), 4.19 (s, 3H), 4.02 (q, *J* = 7.2 Hz, 2H), 2.70 (s, 3H), 0.89 (t, *J* = 7.2 Hz, 3H) ppm. ^13^C NMR (100 MHz, CDCl_3_) δ 167.5 (CO), 164.4 (C), 161.7 (C), 158.4 (C), 150.7 (C), 145.6 (C), 143.0 (C), 132.6 (C), 131.4 (C), 130.7 (CH), 130.6 (CH), 129.0 (CH), 127.7 (CH), 127.6 (CH), 122.1 (C), 115.0 (CH), 114.7 (CH), 108.8 (C), 61.5 (CH_2_), 33.9 (CH_3_), 14.0 (CH_3_), 13.7 (CH_3_) ppm. HRMS (ESI+): m/z calcd for C_23_H_21_N_3_O_2_S [M+H]^+^ 422.1300, found 422.1317.

Synthesis of ethyl 1-methyl-4-(methylthio)-3-phenyl-6-(p-tolyl)-1*H*-pyrazolo[3,4-*b*]pyridine-5-carboxylate **16j**.

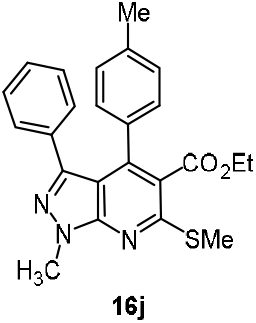

The reaction of 1-methyl-3-phenyl-1*H*-pyrazol-5-amine **7d** (100 mg, 0.57 mmol) and ethyl 2-(4-methylbenzoyl)-3,3-bis(methylthio)acrylate **15f** (176 mg, 0.57 mmol) and a catalytic amount of trifluoroacetic acid (TFA, 19 mg, 30 mol%) in toluene (4 mL) reflux by following the general procedure given above resulted in **16j**, R_f_ = 0.8. Yield: 190 mg (80%). MP: 141.9 °C. UV (λ_max_, nm) 324 (log ε = 2.8107), 256 (log ε = 3.042). IR (KBr, cm^-1^) 3446, 3199, 2987, 2922, 2835, 2724, 2578, 1646, 1519, 1456, 1387, 1246, 1167, 1032, 957, 823, 743, 653, 568, 440. ^1^H NMR (400 MHz, CDCl_3_) δ 7.11 (d, J = 6.8 Hz, 1H), 7.02 - 6.98 (m, 4H), 6.96 - 6.88 (m, 4H), 4.18 (s, 3H), 4.08 (q, J = 7.2 Hz, 2H), 2.70 (s, 3H), 2.27 (s, 3H), 0.95 (t, J = 7.2 Hz, 3H) ppm. ^13^C NMR (100 MHz, CDCl_3_) δ 167.5 (CO), 158.1 (C), 150.9 (C), 145.7 (C), 144.4 (C), 138.3 (C), 132.8 (C), 129.0 (CH), 128.8 (CH), 128.4 (CH), 127.4 (CH), 127.3 (CH), 122.2 (C), 108.9 (C), 61.6 (CH_2_), 33.9 (CH_3_), 14.0 (CH_3_), 13.7 (CH_3_) ppm. HRMS (ESI+): m/z calcd for C_23_H_21_N_3_O_2_S [M+H]^+^ 418.1511, found 418.1602.

Synthesis of ethyl 6-(4-methoxyphenyl)-1-methyl-4-(methylthio)-3-phenyl-1*H*-pyrazolo[3,4-*b*]pyridine-5-carboxylate **16k**.

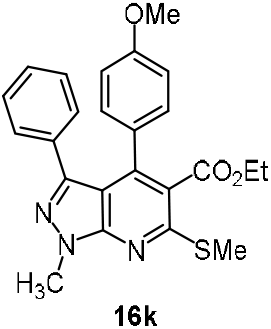

The reaction of 1-methyl-3-phenyl-1*H*-pyrazol-5-amine **7d** (100 mg, 0.57 mmol) and ethyl 2-(4-methoxybenzoyl)-3,3-bis(methylthio)acrylate **15g** (185 mg, 0.57 mmol) and a catalytic amount of trifluoroacetic acid (TFA, 19 mg, 30 mol%) in toluene (4 mL) reflux by following the general procedure given above resulted in **16k**. R_f_ = 0.8. Yield: 210 mg (85%). MP: 133 °C. UV (λ_max_, nm) 324 (log ε = 2.360), 254 (log ε = 2.520). IR (KBr, cm^-1^) 3448, 2986, 2834, 2720, 1689, 1648, 1514, 1462, 1393, 1241, 1393, 1241, 1168, 1024, 936, 823, 746, 450. ^1^H NMR (400 MHz, CDCl_3_) δ 7.15 - 7.11 (m, 1H), 7.05 - 6.97 (m, 6H), 6.63 - 6.60 (m, 2H), 4.17 (s, 3H), 4.10 (q, J = 7.1 Hz, 2H), 3.73 (s, 3H), 2.70 (s, 3H), 0.99 (t, J = 7.1 Hz, 3H) ppm. ^13^C NMR (100 MHz, CDCl_3_) δ 167.9 (C), 159.9 (C), 158.0 (C), 150.0 (C), 145.7 (C), 143.9 (C), 132.9 (C), 130.2 (CH), 129.1 (C), 127.7 (CH), 127.5 (CH), 127.4 (CH), 122.2 (C), 113.3 (C), 108.9 (C), 61.6 (CH_2_), 55.4 (CH_3_), 33.9 (CH_3_), 13.9 (CH_3_), 13.8 (CH_3_) ppm. HRMS (ESI+): m/z calcd for C_24_H_23_N_3_O_3_S [M+H]^+^ 434.1460, found 434.1545.

Synthesis of ethyl 3-(4-chlorophenyl)-1-methyl-4-(methylthio)-6-phenyl-1*H*-pyrazolo[3,4-*b*]pyridine-5-carboxylate **16l**.

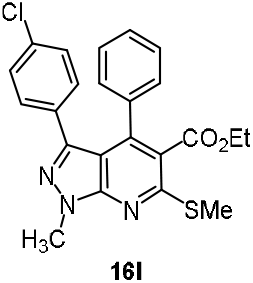

The reaction of 3-(4-chlorophenyl)-1-methyl-1*H*-pyrazol-5-amine **7f** (100 mg, 0.48 mmol) and ethyl 2-benzoyl-3,3-bis(methylthio)acrylate **15c** (142 mg, 0.48 mmol) and a catalytic amount of trifluoroacetic acid (TFA, 18 mg, 30 mol%) in toluene (4 mL) reflux by following the general procedure given above resulted in **16l**. R_f_ = 0.8. Yield: 175 mg (83%). MP: 122.2 °C. UV (λ_max_, nm) 322.5 (log ε = 2.576), 252.5 (log ε = 2.894). IR (KBr, cm^-1^) 3444, 2977, 2926, 2825, 1636, 1516, 1458, 1389, 1236, 1157, 1022, 947, 831, 741, 633, 535, 420. ^1^H NMR (400 MHz, CDCl_3_) δ 7.14 (d, J = 6.2 Hz, 1H), 7.05 (t, J = 7.9 Hz, 2H), 7.04 (dd, J = 7.8 Hz, 2H), 6.96 (dd, J = 6.6 Hz, 2H), 6.87 (td, J = 4.1 Hz, 2H), 4.16 (s, 3H) 4.02 (q, J = 7.1 Hz, 2H), 2.70 (s, 3H), 0.88 (t, J = 7.1 Hz, 3H) ppm. ^13^C NMR (100 MHz, CDCl_3_) δ 167.9 (CO), 158.5 (C), 150.8 (C), 144.3 (C), 144.0 (C), 135.5 (C), 133.6 (C), 131.3 (C), 130.2 (CH), 129.2 (CH), 128.8 (CH), 128.7 (CH), 128.6 (CH), 128.0 (CH), 127.7 (CH), 122.3 (C), 108.7 (C), 61.9 (CH_2_), 33.9 (CH_3_), 14.0 (CH_3_), 13.6 (CH_3_) ppm. HRMS (ESI+): m/z calcd for C_24_H_23_N_3_O_2_S [M+H]^+^ 438.0965, found 438.1036.

Synthesis of ethyl 3-(4-chlorophenyl)-6-(4-fluorophenyl)-1-methyl-4-(methylthio)-1*H*-pyrazolo[3,4-*b*]pyridine-5-carboxylate **16m**.

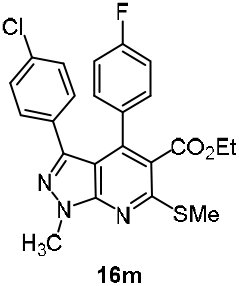

The reaction of 3-(4-chlorophenyl)-1-methyl-1*H*-pyrazol-5-amine **7f** (100 mg, 0.48 mmol) and ethyl 2-(4-fluorobenzoyl)-3,3-bis(methylthio)acrylate **15d** (150 mg, 0.48 mmol) and a catalytic amount of trifluoroacetic acid (TFA, 18 mg, 30 mol%) in toluene (4 mL) reflux by following the general procedure given above resulted in **16m**. R_f_ = 0.8. Yield: 184 mg (84%). MP: 132.5 °C. UV (λ_max_, nm) 328 (log ε = 2.356), 259 (log ε = 1.625). IR (KBr, cm^-1^) 3448, 2987, 2922, 2835, 1646, 1514, 1456, 1386, 1246, 1167, 1032, 957, 833, 743, 653, 565, 440. ^1^H NMR (400 MHz,) δ 7.08 - 7.02 (m, 4H), 6.90 - 6.85 (m, 4H), 4.18 (s, 3H), 4.07 (q, J = 7.1 Hz, 2H), 2.70 (s, 3H), 0.95 (t, J = 7.1 Hz, 3H) ppm. ^13^C NMR (100 MHz, CDCl_3_) δ 167.1 (CO), 158.7 (C), 150.9 (C), 144.2 (C), 142.6 (C), 134.1 (C), 131.4 (C), 130.9 (CH), 130.9 (CH), 130.3 (CH), 127.9 (CH), 122.7 (C), 115.3 (CH), 115.1 (CH), 108.8 (C), 61.6 (CH_2_), 34.0 (CH_3_), 14.0 (CH_3_), 13.9 (CH_3_) ppm. HRMS (ESI+): m/z calcd for C_24_H_23_N_3_O_2_S [M+H]^+^ 456.0871, found 456.0945.

Synthesis of ethyl 3-(4-chlorophenyl)-1-methyl-4-(methylthio)-6-(p-tolyl)-1*H*-pyrazolo[3,4-*b*]pyridine-5-carboxylate **16n**.

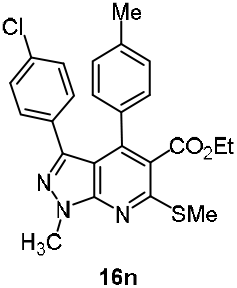

The reaction of 3-(4-chlorophenyl)-1-methyl-1*H*-pyrazol-5-amine **7f** (100 mg, 0.48 mmol) and ethyl 2-(4-methylbenzoyl)-3,3-bis(methylthio)acrylate **15f** (148 mg, 0.48 mmol and a catalytic amount of trifluoroacetic acid (TFA, 17 mg, 30 mol%) in toluene (4 mL) reflux by following the general procedure given above resulted in **16n**. R_f_ = 0.8. Yield: 185 mg (85%). MP: 129.8 °C. UV (λ_max_, nm) 329 (log ε = 2.237), 255.5 (log ε = 2.599). IR (KBr, cm^-1^) 3446, 2987, 2922, 2835, 1646, 1514, 1456, 1386, 1246, 1167, 1032, 957, 833, 743, 653, 565, 440. ^1^H NMR (400 MHz, CDCl_3_) δ 6.97 - 6.88 (m, 8H), 4.17 (s, 3H) 4.07 (q, J = 7.1 Hz, 2H), 2.70 (s, 3H), 2.33 (s, 3H), 0.95 (t, J = 7.1 Hz, 3H) ppm. ^13^C NMR (100 MHz, CDCl_3_) δ 167.4 (CO), 158.4 (C), 150.8 (C), 144.4 (C), 144.0 (C), 138.6 (C), 133.6 (C), 132.6 (C), 131.5 (C), 130.3 (CH), 128.9 (CH), 128.6 (CH), 127.6 (CH), 122.2 (C), 108.9 (C), 61.6 (CH_2_), 33.9 (CH_3_), 14.0 (CH_3_), 13.8 (CH_3_) ppm. HRMS (ESI+): m/z calcd for C_24_H_23_N_3_O_2_S [M+H]^+^ 452.1121, found 452.1228.

Synthesis of ethyl 3-(4-chlorophenyl)-6-(4-methoxyphenyl)-1-methyl-4-(methylthio)-1*H*-pyrazolo[3,4-*b*]pyridine-5-carboxylate **16o**.

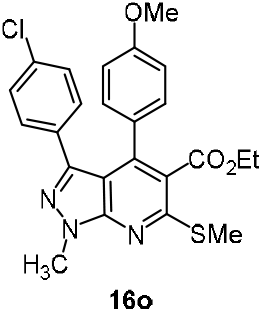

The reaction of 3-(4-chlorophenyl)-1-methyl-1*H*-pyrazol-5-amine **7f** (100 mg, 0.48 mmol) and ethyl 2-(4-methoxybenzoyl)-3,3-bis(methylthio)acrylate **15g** (179 mg, 0.48 mmol and a catalytic amount of trifluoroacetic acid (TFA, 17 mg, 30 mol%) in toluene (4 mL) reflux by following the general procedure given above resulted in **16o**. R_f_ = 0.8. Yield: 180 mg (80%). MP: 130.2 °C. UV (λ_max_, nm) 323 (log ε = 2.763), 257.5 (log ε = 2.963). IR (KBr, cm^-1^) 3446, 2987, 2922, 2835, 1646, 1514, 1456, 1386, 1246, 1167, 1032, 957, 833, 743, 653, 565, 440. ^1^H NMR (400 MHz, CDCl_3_) δ 7.11 (d, J = 6.8 Hz, 1H), 7.02 - 6.88 (m, 8H), 4.18 (s, 3H), 4.08 (q, J = 7.1 Hz, 2H), 2.70 (s, 3H), 2.27 (s, 3H), 0.95 (t, J = 7.1 Hz, 3H) ppm. ^13^C NMR (100 MHz, CDCl_3_) δ δ 167.7 (CO), 160.0 (C), 158.2 (C), 150.7 (C), 144.3 (C), 143.6 (C), 133.4 (C), 131.4 (C), 130.2 (C), 130.1 (CH), 128.2 (C), 127.6 (CH), 127.4 (C), 122.3 (C), 108.9 (C), 61.6 (CH_2_), 55.4 (CH_3_), 33.8 (CH_3_), 13.8 (CH_3_), 13.7 (CH_3_) ppm. HRMS (ESI+): m/z calcd for C_24_H_23_N_3_O_2_S [M+H]^+^ 468.1070, found 468.1102.

Synthesis of ethyl 3-(4-bromophenyl)-1-methyl-4-(methylthio)-6-phenyl-1*H*-pyrazolo[3,4-*b*]pyridine-5-carboxylate **16p**.

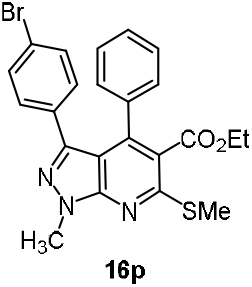

The reaction of 3-(4-bromophenyl)-1-methyl-1*H*-pyrazol-5-amine **7g** (100 mg, 0.39 mmol) and ethyl 2-(4-methylbenzoyl)-3,3-bis(methylthio)acrylate **15c** (115 mg, 0.39 mmol) and a catalytic amount of trifluoroacetic acid (TFA, 13 mg, 30 mol%) in toluene (4 mL) reflux by following the general procedure given above resulted in **16p**. R_f_ = 0.8. Yield: 150 mg (80%). MP: 142.4 °C. UV (λ_max_, nm) 355.5 (log ε = 2.343), 283.5 (log ε = 1.892). IR (KBr, cm^-1^) 3446, 2987, 2922, 2835, 1646, 1514, 1456, 1386, 1246, 1167, 1032, 957, 833, 743, 653, 565, 440. ^1^H NMR (400 MHz, CDCl_3_) δ 7.23 - 7.18 (m, 1H), 7.09 - 7.03 (m, 4H), 7.00 - 6.98 (d, J = 6.0 Hz, 2H), 6.77 - 6.74 (t, J = 4.4 Hz, 2H), 4.14 (s, 3H), 4.08 (q, J = 6.8 Hz, 2H), 2.70 (s, 3H), 0.82 (t, J = 7.2 Hz, 3H) ppm. ^13^C NMR (100 MHz, CDCl_3_) δ 167.6 (CO), 159.9 (C), 158.0 (C), 150.8 (C), 145.7 (C), 143.9 (C), 132.9 (C), 130.2 (CH), 129.1 (CH), 127.7 (CH), 127.5 (CH), 127.4 (C), 122.2 (C), 113.3 (CH), 108.9 (C), 61.7 (CH_2_), 55.4 (C), 33.9 (CH_3_), 13.9 (CH_3_), 13.8 (CH_3_) ppm. HRMS (ESI+): m/z calcd for C_23_H_20_BrN_3_O_2_S [M+H]^+^ 482.0460, found 482.0527.

Synthesis of ethyl 3-(4-bromophenyl)-6-(4-fluorophenyl)-1-methyl-4-(methylthio)-1*H*-pyrazolo[3,4-*b*]pyridine-5-carboxylate **16q**.

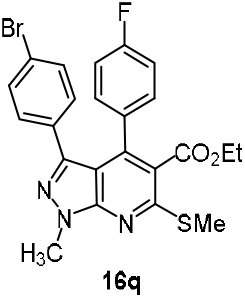

The reaction of 3-(4-bromophenyl)-1-methyl-1*H*-pyrazol-5-amine **7g** (100 mg, 0.39 mmol) and ethyl 2-(4-fluorobenzoyl)-3,3-bis(methylthio)acrylate **15d** (122 mg, 0.39 mmol) and a catalytic amount of trifluoroacetic acid (TFA, 13 mg, 30 mol%) in toluene (4 mL) reflux by following the general procedure given above resulted in **16q**. R_f_ = 0.8. Yield: 158 mg (81%). MP: 147.2 °C. UV (λ_max_, nm) 349.5 (log ε = 2.451), 256 (log ε = 1.904). IR (KBr, cm^-1^) 3446, 2987, 2922, 2835, 1646, 1514, 1456, 1386, 1246, 1167, 1032, 957, 833, 743, 653, 565, 440. ^1^H NMR (400 MHz, CDCl_3_) δ 7.13 (d, J = 8.0 Hz, 2H), 7.01-6.98 (m, 2H), 6.83-6.78 (m, 4H), 4.11 (s, 3H) 4.01 (q, J = 8.0 Hz, 2H), 2.64 (s, 3H), 0.91 (t, J = 8.0 Hz, 3H) ppm. ^13^C NMR (100 MHz, CDCl_3_) δ 167.9 (C), 158.1 (C), 148.2 (C), 142.5 (C), 136.4 (CH), 130.7 (CH), 130.5 (CH), 130.4 (CH), 126.2 (C), 122.0 (C), 120.4 (C), 116.1 (CH), 116.0(CH), 114.9 (CH), 81.0 (CH_2_), 61.7 (CH_2_), 33.9 (CH_3_), 13.9 (CH_3_), 13.6 (CH_3_) ppm. HRMS (ESI+): m/z calcd for C_23_H_19_BrFN_3_O_2_S [M+H]^+^ 500.0365, found 500.1602.

Synthesis of ethyl 3-(4-bromophenyl)-1-methyl-4-(methylthio)-6-(p-tolyl)-1*H*-pyrazolo[3,4-*b*]pyridine-5-carboxylate **16r**.

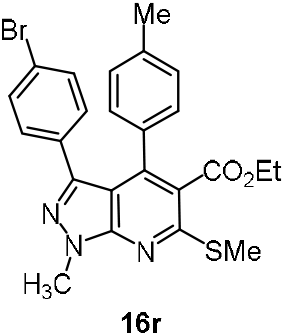

The reaction of 3-(4-bromophenyl)-1-methyl-1*H*-pyrazol-5-amine **7g** (100 mg, 0.39 mmol) and ethyl 2-(4-methylbenzoyl)-3,3-bis(methylthio)acrylate **15f** (120 mg, 0.39 mmol) and a catalytic amount of TFA (13 mg, 30 mol%) in toluene (4 mL) by following the general procedure given above resulted in **16r**. R_f_ = 0.8. Yield: 164 mg (84%). MP: 140.9 °C. UV (λ_max_, nm) 324 (log ε = 2.635), 256 (log ε = 2.850). IR (KBr, cm^-1^) 3446, 2987, 2922, 2835, 1646, 1514, 1456, 1386, 1246, 1167, 1032, 957, 833, 743, 653, 565, 440. ^1^H NMR (400 MHz, CDCl_3_) δ 7.12 (d, J = 8.4 Hz, 2H), 6.94 - 6.93 (m, 4H), 6.82 (d, J = 8.8 Hz, 2H), 4.17 (s, 3H) 4.07 (q, J = 7.2 Hz, 2H), 2.70 (s, 3H), 2.34 (s, 3H), 0.96 (t, J = 7.2 Hz, 3H) ppm. ^13^C NMR (100 MHz, CDCl_3_) δ 167.9 (C), 158.1 (C), 150.8 (C), 145.7 (C), 144.4 (C), 138.3 (C), 132.5 (C), 129.0 (C), 128.8 (CH), 128.4 (CH), 127.4 (CH), 127.3 (CH), 122.2 (CH), 108.9 (C), 61.6 (CH_2_), 33.9 (CH_3_), 14.0 (CH_3_), 13.8 (CH_3_), 13.5 (CH_3_) ppm. HRMS (ESI+): m/z calcd for C_24_H_22_BrN_3_O_2_S [M+H]^+^ 496.0616, found 496.1602.

Synthesis of ethyl 3-(4-bromophenyl)-6-(4-methoxyphenyl)-1-methyl-4-(methylthio)-1*H*-pyrazolo[3,4-*b*]pyridine-5-carboxylate **16s**.

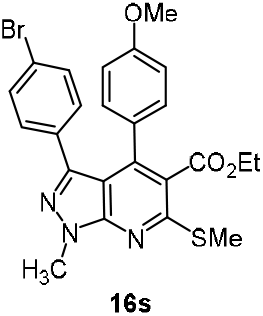

The reaction of 3-(4-bromophenyl)-1-methyl-1*H*-pyrazol-5-amine **7g** (100 mg, 0.39 mmol) and ethyl 2-(4-methoxybenzoyl)-3,3-bis(methylthio)acrylate **15g** (129 mg, 0.39 mmol) in toluene (4 mL) a catalytic amount of (trifluoroacetic acid) TFA (13 mg, 30 mol%) by following the general procedure given above resulted in **16s**. R_f_ = 0.8. Yield: 170 mg (85%). MP: 149.2 °C. UV (λ_max_, nm) 352 (log ε = 2.497), 285 (log ε = 2.816). IR (KBr, cm^-1^) 3446, 2987, 2922, 2835, 1646, 1514, 1456, 1386, 1246, 1167, 1032, 957, 833, 743, 653, 565, 440. ^1^H NMR (400 MHz, CDCl_3_) δ 7.19 (d, J = 8.0 Hz, 2H), 6.99 (d, J = 8.0 Hz, 2H), 6.88 (d, J = 8.0 Hz, 2H), 6.69 (d, J = 8.0 Hz, 2H), 4.19 (s, 3H) 4.13 (q, J = 8.0 Hz, 2H), 3.80 (s, 3H), 2.73 (s, 3H), 1.02 (t, J = 8.0 Hz, 3H) ppm. ^13^C NMR (100 MHz, CDCl_3_) δ 167.7 (CO), 160.0 (C), 158.3 (C), 150.7 (C), 144.3 (C), 143.6 (C), 131.8 (C), 130.5 (CH), 130.5 (CH), 130.1 (CH), 127.4 (C), 122.3(C), 121.6 (CH), 113.4 (CH), 108.9 (C), 61.6 (CH_2_), 55.5 (CH_3_), 33.8 (CH_3_), 13.8 (CH_3_), 13.7 (CH_3_) ppm. HRMS (ESI+): m/z calcd for C_24_H_22_BrN_3_O_3_S [M+H]^+^ 512.0565, found 512.0633.

Synthesis of ethyl 3-(4-methoxyphenyl)-1-methyl-4-(methylthio)-6-phenyl-1*H*-pyrazolo[3,4-*b*]pyridine-5-carboxylate **16t**.

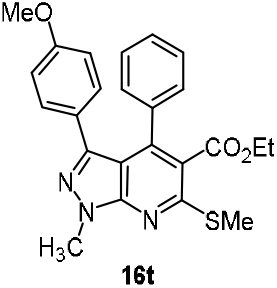

The reaction of 3-(4-methoxyphenyl)-1-methyl-1*H*-pyrazol-5-amine **7h** (100 mg, 0.49 mmol) and ethyl 2-benzoyl-3,3-bis(methylthio)acrylate **15c** (145 mg, 0.49 mmol) and a catalytic amount of trifluoroacetic acid (TFA, 16 mg, 30 mol%) in toluene (4 mL) reflux by following the general procedure given above resulted in **16t**. R_f_ = 0.8. Yield: 183 mg (86%). MP: 152.9 °C. UV (λ_max_, nm) 333 (log ε = 2.434), 254 (log ε = 2.741). IR (KBr, cm^-1^) 2984, 2931, 2837, 1723, 1723, 1566, 1533, 1451, 1235, 1165, 1027, 833, 759, 436, 401. ^1^H NMR (400 MHz, CDCl_3_) δ 7.15 (d, J = 7.6 Hz, 1H), 7.07 (d, J = 8.0 Hz, 2H), 7.00 (d, J = 2.0 Hz, 2H), 6.80 (d, J = 2.0 Hz, 2H), 6.43 (d, J = 2.0 Hz, 2H), 4.1 (3H, s), 3.96 - 3.91 (q, J = 7.2 Hz, 2H), 3.65 (s, 3H), 2.62 (s, 3H), 0.82 (t, J = 7.2 Hz, 3H) ppm. ^13^C NMR (100 MHz, CDCl_3_) δ 167.6 (CO), 159.0 (C), 158.2 (C), 150.8 (C), 145.4 (C), 144.2 (C), 135.7 (C), 130.1 (CH), 129.0 (CH), 128.3 (CH), 127.9 (CH), 125 (C), 122.0 (C), 113.0 (CH), 108.6 (CH), 61.7 (CH_2_), 55.2 (CH_3_), 33.8 (CH_3_), 14.02 (CH_3_), 13.7 (CH_3_) ppm. HRMS (ESI+): m/z calcd for C_24_H_23_N_3_O [M+H]^+^ 434.1460, found 434.1543.

Synthesis of ethyl 6-(4-fluorophenyl)-3-(4-methoxyphenyl)-1-methyl-4-(methylthio)-1*H*-pyrazolo[3,4-*b*]pyridine-5-carboxylate **16u**.

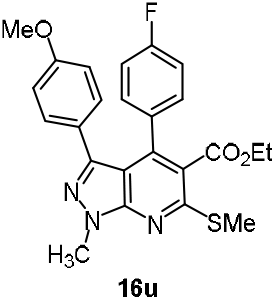

The reaction of 3-(4-methoxyphenyl)-1-methyl-1*H*-pyrazol-5-amine **7h** (60 mg, 0.29 mmol) and ethyl 2-(4-fluorobenzoyl)-3,3-bis(methylthio)acrylate **15d** (91 mg, 0.29 mmol) and a catalytic amount of trifluoroacetic acid (TFA, 10 mg, 30 mol%) in toluene (4 mL) reflux by following the general procedure given above resulted in **16u**. R_f_ = 0.8. Yield: 105 mg (81%). MP: 151.3 °C. UV (λ_max_, nm) 329 (log ε = 2.516), 252 (log ε = 2.652). IR (KBr, cm^-1^) 2987, 2934, 2831, 1721, 1604, 1564, 1506, 1230, 1165, 1015, 835, 402. ^1^H NMR (400 MHz, CDCl_3_) δ 7.25 (d, J = 5.6 Hz, 1H), 7.05 (d, J = 5.2 Hz, 2H), 6.90 - 6.88 (m, 3H), 6.59 (d, J = 8.0 Hz, 2H), 4.16 (s, 3H), 4.08 - 4.04 (q, J = 4.8 Hz, 2H), 3.73 (s, 3H), 2.70 (s, 3H), 0.97 (t, J = 7.2 Hz, 3H) ppm. ^13^C NMR (100 MHz, CDCl_3_) δ 167.3 (CO), 164.4 (C), 161.7 (C), 159.3 (C), 158.3 (C), 150.7 (C), 145.3 (C), 131.6 (C), 131.5 (C), 130.8 (CH), 130.7 (CH), 130.2 (CH), 128.4 (C), 125.2 (C), 122.1 (C), 115.0 (CH), 114.8 (CH), 113.2 (CH), 108.7 (C), 61.7 (CH_2_), 55.3 (CH_3_), 33.8 (CH_3_), 14.0 (CH_3_), 13.7 (CH_3_) ppm. HRMS (ESI+): m/z calcd for C_24_H_22_N_3_O_3_SF [M+H]^+^ 452.1366, found 452.1444.

Synthesis of ethyl 6-(4-chlorophenyl)-3-(4-methoxyphenyl)-1-methyl-4-(methylthio)-1*H*-pyrazolo[3,4-*b*]pyridine-5-carboxylate **16v**.

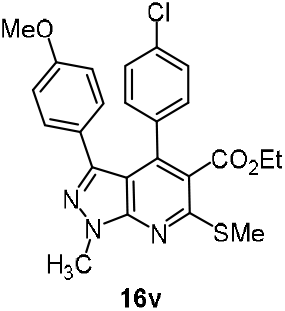

The reaction of 3-(4-methoxyphenyl)-1-methyl-1*H*-pyrazol-5-amine **7h** (100 mg, 0.49 mmol) and ethyl 2-(4-chlorobenzoyl)-3,3-bis(methylthio)acrylate **15e** (162 mg, 0.49 mmol) and a catalytic amount of trifluoroacetic acid (TFA, 17 mg, 30 mol%) in toluene (4 mL) reflux by following the general procedure given above resulted in **16v**. R_f_ = 0.8. Yield: 182 mg (80%). MP: 147.5 °C. UV (λ_max_, nm 332.5 (log ε = 2.521), 253.5 (log ε = 2.955). IR (KBr, cm^-1^) 3738, 2982, 2931, 2837, 1724, 1564, 1237, 1165, 1024, 836, 436, 401. ^1^H NMR (400 MHz, CDCl_3_) δ 7.11 (d, J = 8.4 Hz, 2H), 7.00 (d, J = 8.4 Hz, 2H), 6.84 (d, J = 8.7 Hz, 2H), 6.56 (d, J = 8.8 Hz, 2H), 4.16 (s, 3H), 4.06 (q, J = 7.2 Hz, 2H), 3.75 (s, 3H), 2.69 (s, 3H), 0.99 - 0.96 (t, J = 7.2 Hz, 3H) ppm. ^13^C NMR (100 MHz, CDCl_3_) δ 167.3 (CO), 159.4 (C), 158.4 (C), 150.7 (C), 145.3 (C), 142.8 (C), 134.6 (C), 134.1 (C), 130.3 (CH), 130.2 (CH), 128.0 (CH), 125.1 (C), 121.8 (C), 113.2 (CH), 108.7 (CH), 61.6 (CH_2_), 55.3 (CH_3_), 33.8 (CH_3_), 14.0 (CH_3_), 13.8 (CH_3_) ppm. HRMS (ESI+): m/z calcd for C_24_H_22_N_3_O_3_SCl [M+H]^+^ 468.1070, found 468.1151.

Synthesis of ethyl 3,6-bis(4-methoxyphenyl)-1-methyl-4-(methylthio)-1*H*-pyrazolo[3,4-*b*]pyridine-5-carboxylate **16w**.

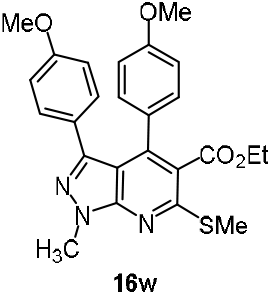

The reaction of 3-(4-methoxyphenyl)-1-methyl-1*H*-pyrazol-5-amine **7h** (80 mg, 0.39 mmol) and ethyl 2-(4-methoxybenzoyl)-3,3-bis(methylthio)acrylate **15g** (127 mg, 0.39 mmol and a catalytic amount of trifluoroacetic acid (TFA, 13 mg, 30 mol%) in toluene (4 mL) reflux by following the general procedure given above resulted in **16w**. R_f_ = 0.8. Yield: 150 mg (83%). MP: 156 °C. UV (λ_max_, nm) 326 (log ε = 2.511), 230 (log ε = 2.772). IR (KBr, cm^-1^) 3620, 2996, 2928, 2843, 1117, 1610, 1567, 1513, 1263, 1171, 1029, 834, 755, 435, 403. ^1^H NMR (400 MHz, CDCl_3_) δ 7.00 (d, J = 8.8 Hz, 2H), 6.91 (d, J = 8.8 Hz, 2H), 6.64 (d, J = 9.2 Hz, 2H), 6.57 (d, J = 8.8 Hz, 2H), 4.16 (s, 3H), 4.02 (q, J = 6.8 Hz 2H,), 3.75 (s, 3H), 3.73 (s, 3H), 2.7 (s, 3H), 1.01 - 0.96 (t, J = 6.8 Hz, 3H) ppm. ^13^C NMR (100 MHz, CDCl_3_) δ 168.0 (CO), 159.9 (C), 159.1 (C), 145.5 (C), 144.0 (C), 131.6 (C), 130.3 (CH), 127.8 (C), 125.5 (C), 114.0 (CH), 113.4 (CH), 113.1 (CH), 108.4 (C), 61.7 (CH_2_), 55.4 (CH_3_), 55.3 (CH_3_), 33.8 (CH_3_), 14.0 (CH_3_), 13.8 (CH_3_) ppm. HRMS (ESI+): m/z calcd for C_24_H_22_N_3_O_3_S [M+H]^+^ 464.1566, found 464.1648.

Synthesis of ethyl 4-(methylthio)-1,3,6-triphenyl-1*H*-pyrazolo[3,4-*b*]pyridine-5-carboxylate **16x**.

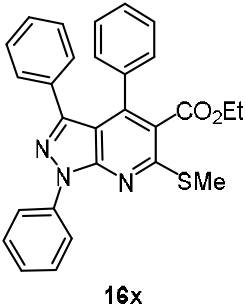

The reaction of 1,3-diphenyl-1*H*-pyrazol-5-amine **7a** (100 mg, 0.42 mmol) and ethyl 2-benzoyl-3,3-bis(methylthio)acrylate **15c** (125 mg, 0.42 mmol) and a catalytic amount of trifluoroacetic acid (TFA, 14 mg, 30 mol%) in toluene (4 mL) reflux by following the general procedure given above resulted in **16x**. R_f_ = 0.8. Yield: 165 mg (80%). MP: 169.6 °C. UV (λ_max_, nm) 329 (log ε = 2.405), 265 (log ε = 2.751). IR (KBr, cm^-1^) 3052, 2981, 2928, 1723, 1593, 1554, 1501, 1455, 1413, 1364, 1290, 1233, 1180, 1127, 1034, 903, 843, 756, 696, 514, 428, 402. ^1^H NMR (400 MHz, CDCl_3_) δ 8.31 (d, J = 7.7 Hz, 2H), 7.46 (t, J = 8.0 Hz, 2H), 7.26 (d, J = 7.4 Hz, 1H), 7.16 - 6.90 (m, 10H), 3.97 (q, J = 7.1 Hz, 2H), 2.66 (s, 3H), 0.81 (t, J = 7.1 Hz, 3H) ppm. ^13^C NMR (100 MHz, CDCl_3_) δ 167.5 (C), 159.0 (C), 150.2 (C), 147.3 (C), 144.3 (C), 139.3 (C), 135.1 (C), 132.4 (C), 129.1 (CH), 129.0 (CH), 128.9 (CH), 128.5 (CH), 127.8 (CH), 127.8 (CH), 127.6 (CH), 126.2 (CH), 122.8 (CH), 121.3 (C), 110.6 (C), 61.7 (CH_2_), 14.4 (CH_3_), 13.6 (CH_3_) ppm. HRMS (ESI+): m/z calcd for C_28_H_23_N_3_O_2_S [M+Na]^+^ 488.1511, found 488.1443.

Synthesis of ethyl 6-(4-fluorophenyl)-4-(methylthio)-1,3-diphenyl-1*H*-pyrazolo[3,4-*b*]pyridine-5-carboxylate **16y**.

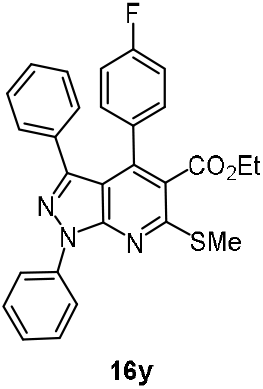

The reaction of 1,3-diphenyl-1*H*-pyrazol-5-amine **7a** (100 mg, 0.42 mmol), ethyl 2-(4-fluorobenzoyl)-3,3-bis(methylthio)acrylate **15d** (143 mg, 0.42 mmol) and a catalytic amount of trifluoroacetic acid (TFA, 14 mg, 30 mol%) in toluene (4 mL) reflux by following the general procedure given above resulted in **16y**. R_f_ = 0.8. Yield: 169 mg (83%). MP: 163.8 °C. UV (λ_max_, nm) 328 (log ε = 2.453), 263 (log ε = 2.816). IR (KBr, cm^-1^) 3061, 2982, 2928, 1723, 1599, 1558, 1504, 1458, 1413, 1366, 1291, 1231, 1177, 1128, 1034, 904, 813, 757, 699, 519, 444, 401. ^1^H NMR (400 MHz, CDCl_3_) δ 8.38 (d, J = 7.8 Hz, 2H), 7.55 (t, J = 7.5 Hz, 2H), 7.34 (d, J = 6.4 Hz, 1H), 7.08 (dt, J = 8.0 Hz, 6H), 6.82 (t, J = 8.8 Hz, 2H), 4.11 (q, J = 7.1 Hz, 2H), 2.73 (s, 3H), 0.99 (t, J = 7.1 Hz, 3H) ppm. ^13^C NMR (100 MHz, CDCl_3_) δ 167.3 (C), 164.2 (C), 161.8 (C), 159.2 (C), 150.2 (C), 147.1 (C), 143.1 (C), 139.3 (C), 132.3 (C), 131.1 (C), 130.8 (CH), 130.7 (CH), 129.2 (CH), 129.1 (CH), 128.0 (CH), 127.7 (CH), 126.2 (CH), 123.0 (C), 121.4 (CH), 115.0 (CH), 114.8 (CH), 110.7 (C), 61.8 (CH_2_), 14.4 (CH_3_), 13.7 (CH_3_) ppm. HRMS (ESI+): m/z calcd for C_28_H_22_N_3_O_2_SF [M+H]^+^ 484.1417, found 484.1528.

Synthesis of ethyl 4-(methylthio)-1,3-diphenyl-6-(p-tolyl)-1*H*-pyrazolo[3,4-*b*]pyridine-5-carboxylate **16z**.

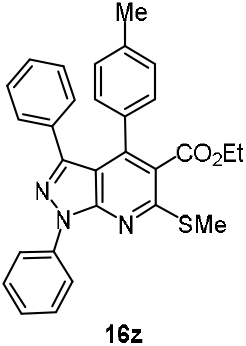

The reaction of 1,3-diphenyl-1*H*-pyrazol-5-amine **7a** (100 mg, 0.42 mmol), ethyl 2-(4-methylbenzoyl)-3,3-bis(methylthio)acrylate **15f** (130 mg, 0.42 mmol and a catalytic amount of trifluoroacetic acid (TFA, 14 mg, 30 mol%) in toluene (4 mL) reflux by following the general procedure given above resulted in **16z**. R_f_ = 0.8. Yield: 163 mg (81%). MP: 171 °C. UV (λ_max_, nm) 329 (log ε = 1.485), 265 (log ε = 1.826). IR (KBr, cm^-1^) 3061, 2982, 2925, 2859, 1724, 1678, 1601, 1558, 1504, 1414, 1367, 1288, 1235, 1179, 1126, 1035, 959, 904, 804, 757, 699, 537, 408. ^1^HNMR (400 MHz, CDCl_3_) δ 8.39 (d, J = 7.7 Hz, 2H), 8.00 (d, J = 7.9 Hz, 1H), 7.53 (d, J = 8.0 Hz, 2H), 7.34 (t, J = 7.2 Hz, 1H), 7.15 (t, J = 6.8 Hz, 1H), 7.05 (t, J = 6.0 Hz, 4H), 6.97 (d, J = 8.0 Hz, 2H), 6.90 (d, J = 8.4 Hz, 2H), 4.09 (q, J = 6.8 Hz, 2H), 2.73 (s, 3H), 2.29 (s, 3H), 0.98 (t, J = 6.8 Hz, 3H) ppm. ^13^C NMR (100 MHz, CDCl_3_) δ 167.6 (C), 158.9 (C), 150.3 (C), 147.4 (C), 144.6 (C), 139.4 (C), 138.5 (C), 132.5 (C), 132.2 (C), 130.3 (CH), 129.3 (CH), 129.2 (CH), 128.8 (CH), 128.5 (CH), 127.6 (CH), 127.5 (CH), 126.1 (CH), 123.0 (C), 121.4 (CH), 110.8 (C), 61.7 (CH_2_), 21.3 (CH_3_), 14.4 (CH_3_), 13.7 (CH_3_) ppm. HRMS (ESI+): m/z calcd for C_29_H_25_N_3_O_2_S [M+H]^+^ 480.1667, found 480.1779.

Synthesis of ethyl 6-(4-methoxyphenyl)-4-(methylthio)-1,3-diphenyl-1*H*-pyrazolo[3,4-*b*]pyridine-5-carboxylate **16aa**.

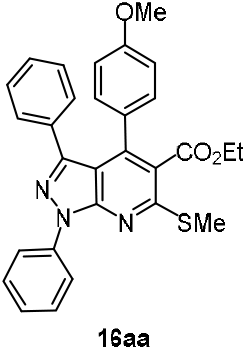

The reaction of 1,3-diphenyl-1*H*-pyrazol-5-amine **7a** (100 mg, 0.42 mmol), ethyl 2-(4-methoxybenzoyl)-3,3-bis(methylthio)acrylate **15g** (137 mg, 0.42 mmol) and a catalytic amount of trifluoroacetic acid (TFA, 14 mg, 30 mol%) in toluene (4 mL) reflux by following the general procedure given above resulted in **16aa**. R_f_ = 0.8. Yield: 177 mg (85%). MP: 176.1 °C. UV (λ_max_, nm) 330 (log ε = 2.521), 269 (log ε = 1.534). IR (KBr, cm^-1^) 3062, 2927, 2847, 1722, 1674, 1602, 1560, 1458, 1364, 1246, 1177, 1125, 1033, 905, 836, 762, 699. ^1^H NMR (400 MHz, CDCl_3_) δ 8.38 (d, J = 8.4 Hz, 2H), 7.53 (t, J = 7.6 Hz, 2H), 7.32 (q, J = 7.2 Hz, 1H), 7.17 (q, J = 6.9 Hz, 1H), 7.06 (t, J = 5.2 Hz, 4H), 7.00 (d, J = 8.0 Hz, 2H), 6.62 (d, J = 8.0 Hz, 2H), 4.11 (q, J = 7.1 Hz, 2H), 3.75 (s, 3H), 2.73 (s, 3H), 1.01 (t, J = 7.1 Hz, 3H) ppm. ^13^C NMR (100 MHz, CDCl_3_) δ 167.5 (CO), 160.1 (C), 158.9 (C), 150.4 (C), 147.2 (C), 144.0 (C), 139.5 (C), 132.7 (C), 130.4 (CH), 129.3 (CH), 129.0 (CH), 127.7 (CH), 127.6 (C), 127.5 (CH), 126.1 (CH), 123.2 (CH), 121.3 (CH), 113.4 (CH), 110.9 (C), 61.6 (CH_2_), 55.4 (CH_3_), 14.4 (CH_3_), 13.9 (CH_3_) ppm. HRMS (ESI+): m/z calcd for C_29_H_25_N_3_O_3_S [M+H]^+^ 496.1600, found 496.1683.

Synthesis of ethyl 3-(4-chlorophenyl)-1,6-dimethyl-4-(methylthio)-1*H*-pyrazolo[3,4-*b*]pyridine-5-carboxylate **17f**.

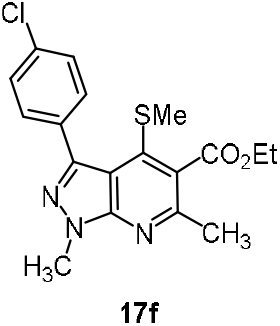

The reaction of 3-(4-chlorophenyl)-1-methyl-1*H*-pyrazol-5-amine **7f** (100 mg, 0.48 mmol) and ethyl 2-(bis(methylthio)methylene)-3-oxobutanoate **15h** (112 mg, 0.48 mmol) and a catalytic amount of trifluoroacetic acid (TFA, 16 mg, 30 mol%) in toluene (4 mL) reflux by following the general procedure given above resulted in **17f**. R_f_ = 0.8. Yield: 180 mg (83%). MP: 142.7 °C. UV (λ_max_, nm) 324 (log ε = 2.356), 229.5 (log ε = 1.704). IR (KBr, cm^-1^) 2926, 2857, 1729, 1569, 1503, 1450, 1243, 1130, 1091, 1014, 965, 828. ^1^H NMR (400 MHz, CDCl_3_) δ 7.78 (d, J = 2.0 Hz, 2H), 7.41 (d, J = 2.0 Hz, 2H), 4.45 (q, J = 7.2 Hz, 2H), 4.15 (s, 3H), 2.63 (s, 3H), 1.87 (s, 3H), 1.43 (t, J = 7.2 Hz, 3H) ppm. ^13^C NMR (100 MHz, CDCl_3_) δ 168.4 (CO), 154.8 (C), 150.7 (C), 143.3 (C), 140.2 (C), 134.8 (C), 131.9 (C), 131.0 (CH), 128.7 (C), 128.3 (CH), 111.5 (C), 61.8 (CH_2_), 34.2 (CH_3_), 23.4 (CH_3_), 19.6 (CH_3_), 14.3 (CH_3_) ppm. HRMS (ESI+): m/z calcd for C_18_H_18_ClN_3_O_2_S [M+H]^+^ 376.0808, found 376.0914.

Synthesis of ethyl 3-(4-methoxyphenyl)-1,6-dimethyl-4-(methylthio)-1*H*-pyrazolo[3,4-*b*]pyridine-5-carboxylate **17g**.

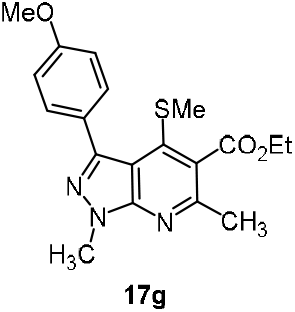

The reaction of 3-(4-methoxyphenyl)-1-methyl-1*H*-pyrazol-5-amine **7h** (200 mg, 0.98 mmol) and ethyl 2-(bis(methylthio)methylene)-3-oxobutanoate **15h** (220 mg, 0.98 mmol and a catalytic amount of trifluoroacetic acid (TFA, 32 mg, 30 mol%) in toluene (10 mL) reflux by following the general procedure given above resulted in **17g**. R_f_ = 0.8. Yield: 290 mg (80%). MP: 148.6 °C. UV (λ_max_, nm) 326 (log ε = 2.260), 297 (log ε = 1.523). IR (KBr, cm^-1^) 2928, 2851, 1726, 1609, 1571, 1526, 1453, 1297, 1244, 1176, 1130, 1027, 899, 835, 736, 610, 436, 401. ^1^H NMR (400 MHz, CDCl_3_) δ 7.72 (d, J = 4 Hz, 2H), 6.94 (d, J = 8.0 Hz, 2H), 4.42 (q, J = 7.1 Hz, 2H), 4.13 (s, 3H), 3.84 (s, 3H), 2.62 (s, 3H), 1.85 (s, 3H), 1.41 (t, J = 12 Hz, 3H) ppm. ^13^C NMR (100 MHz, CDCl_3_) δ 168.5 (CO), 160.0 (C), 154.4 (C), 150.7 (C), 144.4 (C), 140.6 (C), 130.9 (CH), 128.2 (C), 125.87 (C), 113.4 (CH), 111.6 (C), 61.7 (CH_2_), 55.2 (OCH_3_), 34.1 (CH_3_), 23.3 (CH_3_), 19.5 (CH_3_), 14.3 (CH_3_) ppm. HRMS (ESI+): m/z calcd for C_19_H_21_N_3_O_3_S [M+H]^+^ 372.1304, found 372.1408.

### 4.3. Reductive elimination of SMe group: General procedure

Synthesis of ethyl 1-methyl-3,6-diphenyl-1*H*-pyrazolo[3,4-*b*]pyridine-5-carboxylate **25a**.

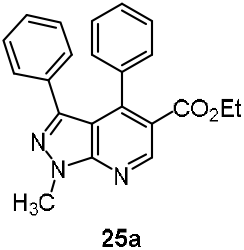

To the solution of ethyl 1-methyl-4-(methylthio)-3,6-diphenyl-1*H*-pyrazolo[3,4-b]pyridine-5-carboxylate **16h** (50 mg, 0.13 mmol) in ethyl acetate (12 mL) Raney Ni (about 10 mg) was added with the help of ethanol (3 mL). The air was replaced with hydrogen using hydrogen balloon and vacuum pump. Then the flask was placed in a pre-heated oil-bath of 50 °C and stirred for 24 h by which time TLC indicated absence of starting material. The cooled reaction mixture was filtered through a bed of Celite. Removal of solvent under reduced pressure resulted in **25a** as analytically pure white solid. R_f_ = 0.9 (hexane and EtOAc, 9.9:0.1). Yield: 37 mg (80%). MP: 152 °C. UV (λ_max_, nm) 350 (log ε = 1.722), 314 (log ε = 2.014). IR (KBr, cm^-1^) 3434, 3189, 2759, 2724, 2578, 1699, 1621, 1419, 1397, 1324, 1185, 1137, 947, 848, 715, 667, 511, 406. ^1^H NMR (400 MHz, CDCl_3_) δ 9.09 (s, 1H), 7.12 - 6.98 (m, 10H), 4.28 (s, 3H), 4.13 (q, J = 8.0 Hz, 2H), 0.98 (t, J = 8.0 Hz, 3H) ppm. ^13^C NMR (100 MHz, CDCl_3_) δ 167.5 (CO), 165.2 (C), 150.7 (C), 149.5 (C), 143.5 (C), 135.7 (C), 132.8 (C), 128.9 (CH), 128.6 (CH), 128.0 (CH), 127.4 (CH), 127.4 (CH), 120.1 (C), 116.1 (C), 112.5 (CH), 108.2 (C), 61.0 (CH_2_), 34.1 (CH_3_), 13.6 (CH_3_) ppm. HRMS (ESI+): m/z calcd for C_22_H_19_N_3_O_2_ [M+H]^+^ 358.1477, found 358.1556.

Synthesis of ethyl 6-(4-fluorophenyl)-1-methyl-3-phenyl-1*H*-pyrazolo[3,4-*b*]pyridine-5-carboxylate **25b**.

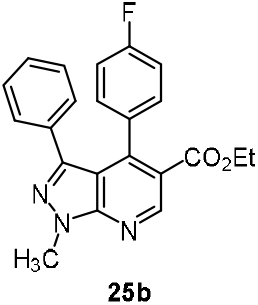

The reductive elimination of SMe group in ethyl 6-(4-fluorophenyl)-1-methyl-4-(methylthio)-3-phenyl −1H-pyrazolo[3,4-b]pyridine-5-carboxylate **16i** (50 mg, 0.12 mmol) by following the general procedure described above provided 40 mg (88%) of **25b** as a white solid. R_f_ = 0.9. MP: 142.4 °C. UV (λ_max_, nm) 356 (log ε = 1.942), 314 (log ε = 2.204). IR (KBr, cm^-1^) 3339, 3089, 2658, 2558, 1690, 1621, 1449, 1369, 1214, 1185, 1137, 957, 627, 521, 401. ^1^H NMR (400 MHz, CDCl_3_) δ 9.11 (s, 1H), 7.21 - 7.04 (m, 1H), 7.03 - 7.01 (t, J = 4.0 Hz, 5H), 6.99 - 6.80 (m, 3H), 4.28 (s, 3H), 4.17 (q, J = 8.0 Hz, 2H), 1.08 (t, J = 8.0 Hz, 3H) ppm. ^13^C NMR (100 MHz, CDCl_3_) δ 166.8 (CO), 161.5 (C), 151.6 (C), 146.5 (C), 146.1 (C), 132.4 (C), 131.7 (C), 130.5 (CH), 130.4 (CH), 129.0 (CH), 127.6 (CH), 127.5 (C), 119.8 (CH), 114.5 (CH), 114.3 (CH), 112.9 (C), 61.1 (CH_2_), 34.3 (CH_3_), 13.7 (CH_3_) ppm. HRMS (ESI+): m/z calcd for C_22_H_18_FN_3_O_2_ [M+H]^+^ 376.1835, found 376.1459.

Synthesis of ethyl 3-(4-bromophenyl)-1-methyl-6-phenyl-1*H*-pyrazolo[3,4-*b*]pyridine-5-carboxylate **25c**.

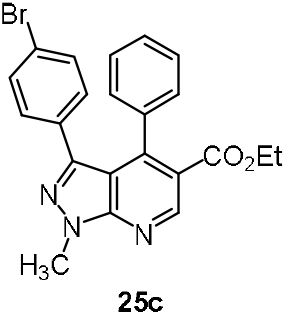

The reductive elimination of SMe group in ethyl 3-(4-bromophenyl)-1-methyl-4-(methylthio)-6-phenyl-1*H*-pyrazolo[3,4-*b*]pyridine-5-carboxylate **16p** (50 mg, 0.11 mmol) by following the general procedure described above provided 40 mg (82%) of **25c** as a white solid. R_f_ = 0.9. Yield: MP: 151 °C. UV (λ_max_, nm) 355 (log ε = 1.827), 314 (log ε = 2.215). IR (KBr, cm^-1^) 3339, 3089, 2658, 2558, 1690, 1621, 1449, 1369, 1214, 1185, 1137, 957, 627, 521, 401. ^1^H NMR (400 MHz, CDCl_3_) δ 9.09 (s, 1H), 7.18 - 6.98 (m, 9H), 4.27 (s, 3H), 4.11 (q, J = 8.0 Hz, 2H), 0.98 (t, J = 8.0 Hz, 3H) ppm. ^13^C NMR (100 MHz, CDCl_3_) δ 167.1 (CO), 151.6 (C), 150.7 (C), 147.5 (C), 146.2 (C), 130.5 (C), 130.4 (CH), 128.9 (CH), 128.7 (CH), 128.6 (CH), 127.4 (CH), 120.0 (C), 112.5 (C), 61.1 (CH_2_), 34.2 (CH_3_), 13.6 (CH_3_) ppm. HRMS (ESI+): m/z calcd for C_22_H_18_N_3_BrO_2_ [M+H]^+^ 436.0582, found 436.0654.

### 4.4. Hydrolysis of the ester group: General procedure

Synthesis of 1-methyl-4-(methylthio)-3,6-diphenyl-1*H*-pyrazolo[3,4-*b*]pyridine-5-carboxylic acid **26a**.

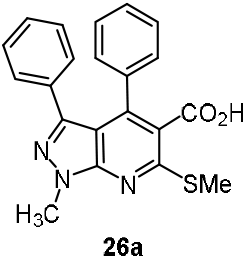

To the solution of ethyl 1-methyl-4-(methylthio)-3,6-diphenyl-1*H*-pyrazolo[3,4-*b*]pyridine-5-carboxylate **16h** (20 mg, 0.04 mmol) in 1,4-dioxane (8 mL), LiOH (4 mg, 0.15 mmol) in water (2 mL) was added and the reaction mixture was stirred for 12 h by which time TLC indicated absence of the ester. Aqueous solution of oxalic acid (0.1 N) was added to the cooled reaction mixture to adjust the pH to 7, then diluted with EtOAc (25 mL). The organic phase was washed with water (2 x 10 mL) and brine (1 x 10 mL) and dried over anhydrous sodium sulphate. Clear organic phase was subjected to evaporation under reduced pressure in a rotator evaporator. Resulting white solid was recrystallized with ethyl acetate in hexanes (98:2) and dichloromethane (DCM). Yield: 15 mg (83%). MP: 153 °C. UV (λ_max_, nm) 329 (log ε = 2.425), 251 (log ε = 2.773). IR (KBr, cm^-1^) 3457, 2928, 1717, 1564, 1493, 1449, 1357, 1262, 1233, 1174, 1079, 1017, 924, 898, 756, 703, 669, 615, 514. ^1^H NMR (400 MHz, CDCl_3_ + DMSO d_6_) δ 7.19 (s, 4H), 7.04 (d, J = 4.0 Hz, 3H), 6.94 - 6.88 (m, 3H), 4.13 (s, 3H), 2.67 (s, 3H) ppm. ^13^C NMR (100 MHz, CDCl_3_) δ 169.3 (CO_2_H), 159.7 (C), 145.4 (C), 138.2 (C), 131.8 (CH), 128.6 (CH), 127.6 (CH), 127.3 (C), 127.1 (CH), 126.1 (CH), 123.2 (CH), 33.6 (CH_3_), 13.8 (CH_3_) ppm. HRMS (ESI+): m/z calcd for C_29_H_25_N_3_O_2_ [M+H]^+^ 376.1114, found 376.1122.

Synthesis of 6-(4-fluorophenyl)-1-methyl-4-(methylthio)-3-phenyl-1*H*-pyrazolo[3,4-*b*]pyridine-5-carboxylic acid **26b**.

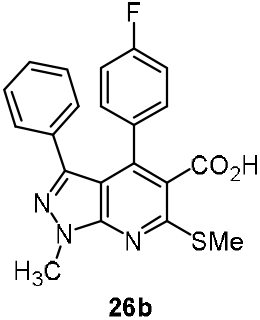

The hydrolysis of the ester group in ethyl 6-(4-fluorophenyl)-1-methyl-4-(methylthio)-3-phenyl-1*H*-pyrazolo[3,4-*b*]pyridine-5-carboxylate **16i** (10 mg, 0.02 mmol) with excess LiOH in 4:1 dioxane and water by following the general procedure described above provided 8.5 mg (94%) of **26b** as a white solid. R_f_ = 0.2. MP: 154 °C. UV (λ_max_, nm) 342 (log ε = 2.505), 258 (log ε = 2.843). IR (KBr, cm^-1^) 3457, 2928, 1717, 1564, 1493, 1448, 1357, 1261, 1233, 1173, 1074, 1014, 923, 891, 755, 701, 662, 614, 510. ^1^H NMR (400 MHz, DMSO d_6_) δ 7.82 (t, J = 4.0 Hz, 3H), 7.42 (t, J = 4.0 Hz, 3H), 7.19 – 7.05 (m, 3H), 4.21 (s, 3H), 2.66 (s, 3H) ppm. ^13^C NMR (100 MHz, DMSO d_6_) δ 169.3 (CO_2_H), 157.5 (C), 145.2 (C), 129.1 (C), 129.0 (C), 128.8 (C), 128.4 (C), 127.9 (C), 127.5 (C), 127.3 (C), 34.1 (C), 13.4 (C) ppm. HRMS (ESI+): m/z calcd for C_21_H_16_FN_3_O_2_S [M+H]^+^394.0947, found 394.1033.

Synthesis of 1-methyl-4-(methylthio)-3-phenyl-6-(p-tolyl)-1H-pyrazolo[3,4-b]pyridine-5-carboxylic acid **26c**.

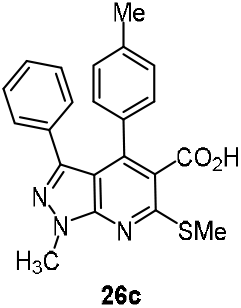

The hydrolysis of ester group in ethyl 1-methyl-4-(methylthio)-3-phenyl-6-(p-tolyl)-1H-pyrazolo[3,4-b]pyridine-5-carboxylate **16j** (20 mg, 0.04 mmol) by following the general procedure described above provided 16 mg (87%) of **26c** as a white solid. R_f_ = 0.2. MP: 157 °C. UV (λ_max_, nm) 356 (log ε = 2.551), 269 (log ε = 2.820). IR (KBr, cm^-1^) 3467, 2948, 1715, 1564, 1497, 1442, 1351, 1262, 1239, 1178, 1074, 1012, 929, 899, 751, 705, 667, 619, 517. ^1^H NMR (400 MHz, DMSO d_6_) δ 7.09-7.04 (m, 1H), 6.96-6.81 (m, 8H), 4.11 (s, 3H), 2.64 (s, 3H), 2.21 (s, 3H) ppm. ^13^C NMR (100 MHz, DMSO d_6_) δ 169.2 (CO2H), 150.0 (C), 156.6 (C), 147.1 (C), 145.6 (C), 145.4 (C), 144.3 (C), 138.2 (C), 132.4 (C), 128.9 (CH), 128.6 (CH), 128.3 (C), 127.3 (CH), 127.2 (CH), 114.5 (C), 108.8 (C), 33.7 (CH_3_), 21.9 (CH_3_), 13.5 (CH_3_) ppm. HRMS (ESI+): m/z calcd for C_22_H_19_N_3_O_2_S [M+H]^+^ 390.1271, found 390.1281.

Synthesis of 6-(4-methoxyphenyl)-1-methyl-4-(methylthio)-3-phenyl-1H-pyrazolo[3,4-b]pyridine-5-carboxylic acid **26d**.

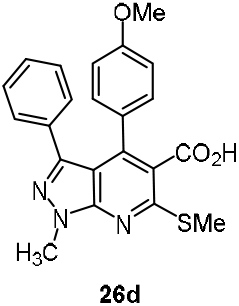

The hydrolysis of ester group in ethyl 6-(4-methoxyphenyl)-1-methyl-4-(methylthio)-3-phenyl-1H-pyrazolo[3,4-b]pyridine-5-carboxylate **16k** (10 mg, 0.02 mmol) with excess LiOH in 4:1 dioxane and water by following the general procedure described above provided 8 mg (86%) of **26d** as a white solid. R_f_ = 0.2. MP: 155 °C. UV (λ_max_, nm) 339 (log ε = 2.695), 272 (log ε = 2.920). IR (KBr, cm^-1^) 3469, 2924, 1719, 1559, 1497, 1444, 1358, 1268, 1237, 1170, 1076, 1014, 925, 899, 754, 709, 669, 619, 519. ^1^H NMR (400 MHz, DMSO d_6_) δ 7.08-6.98 (m, 1H), 6.97-6.91 (m, 6H), 6.56 (d, J = 8.0 Hz, 2H), 4.12 (s, 3H), 3.68 (s, 3H), 2.66 (s, 3H) ppm. ^13^C NMR (100 MHz, DMSO d_6_) ^13^C NMR (100 MHz, DMSO d_6_) δ 168.5 (CO_2_H), 161.7 (C), 159.8 (C), 158.2 (C), 156.8 (C), 154.0 (C), 150.2 (C), 149.8 (C), 144.7 (C), 142.8 (C), 139.2 (C), 132.5 (C), 130.4 (C), 130.4 (CH), 128.6 (CH), 127.6 (C), 127.4 (CH), 127.3 (CH), 126.2 (CH), 116.9 (C), 113.6 (C), 111.2 (C), 108.5 (C), 55.5 (CH_3_), 33.7 (CH_3_), 13.5 (CH_3_) ppm. HRMS (ESI+): m/z calcd for C_22_H_19_N_3_O_3_S [M+H]^+^406.1147, found 406.1219.

### 4.5. Suzuki coupling of the aryl bromide with phenyl boronic acids

Synthesis of ethyl 3-([1,1’-biphenyl]-4-yl)-1-methyl-4-(methylthio)-6-phenyl-1H-pyrazolo[3,4-b]pyridine-5-carboxylate **27**.

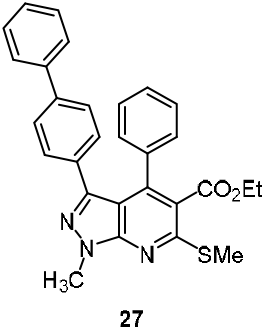

To a solution of ethyl 3-(4-bromophenyl)-1,4-dimethyl-6-phenyl-1H-pyrazolo[3,4-b]pyridine-5-carboxylate **16p** (20 mg, 0.041 mmol), PhB(OH)_2_ (phenylboronic acid) (10 mg, 0.082 mmol), K_3_PO_4_ (26 mg, 0.12 mmol), PPh_3_ (triphenylphosphine) (32 mg, 0.12 mmol) a catalytic amount of Pd(OAc)_2_ (palladium(II) acetate) (0.9 mg, 0.004 mmol) were taken and applied high vacuum for 10 minutes. To this (1,2-Dichloroethane) DCE (2 mL) was added under N_2_ atm, then the reaction mixture was refluxed for 10 h. The reaction was progressively monitored by TLC using mixture of hexane and ethyl acetate (9.0:1.0). Upon completion of the reaction as indicated by TLC for absence of **16p**, the reaction mixture was diluted with 20 mL DCM. The organic phase was washed with water (2 x 10 mL) and brine (1 x 10 mL) and dried over anhydrous sodium sulphate. Clear organic phase was subjected to evaporation under reduced pressure in a rotator evaporator (30 rpm) at rt. Resulting viscous liquid was subjected to column chromatography with increasing percentage of ethyl acetate in hexanes (0.1% to 2%). Evaporation of pooled fractions having **27** resulted in an analytically pure white solid. R_f_ = 0.9. Yield: 15mg (78%). MP: 209.4 °C. UV (λ_max_, nm) 318 (log ε = 2.541), 253 (log ε = 2.888). IR (KBr, cm^-1^) 3444, 3195, 2954, 2839, 2726, 2575, 1755, 1697, 1609, 1515, 1465, 1399, 1325, 1186, 1129, 948, 847, 735, 686, 520, 409. ^1^H NMR (400 MHz, CDCl_3_) δ 7.37 - 6.86 (m, 14H), 4.33 (s, 3H), 4.15 (q, J = 8.0 Hz, 2H), 3.46 (s, 3H), 1.03 (t, J = 8.0 Hz, 3H) ppm. ^13^C NMR (100 MHz, CDCl_3_) δ 167.5 (C), 159.0 (C), 150.2 (C), 147.3 (C), 144.3 (C), 133.3 (CH), 129.1 (CH), 128.7 (CH), 128.6 (CH), 127.9 (CH), 115.1 (C), 108.5 (C), 61.2 (CH_2_), 33.8 (CH_3_), 13.9 (CH_3_) ppm. HRMS (ESI+): m/z calcd for C_29_H_25_N_3_O_2_ [M+H]^+^ 480.1667, found 480.1741.

### 4.6. Procedure for in-vitro MTB MABA assay

The inoculum was prepared from fresh LJ medium re-suspended in 7H9-S medium (7H9 broth, 0.1% casitone, 0.5% glycerol, supplemented oleic acid, albumin, dextrose, and catalase [OADC]), adjusted to a McFarland tube No. 1, and diluted 1:20; 100 µl was used as inoculum. Each drug stock solution was thawed and diluted in 7H9-S at four-fold the final highest concentration tested. Serial two-fold dilutions of each drug were prepared directly in a sterile 96-well microtiter plate using 100 µl 7H9-S. A growth control containing no antibiotic and a sterile control were also prepared on each plate. Sterile water was added to all perimeter wells to avoid evaporation during the incubation. The plate was covered, sealed in plastic bags, and incubated at 37 °C in normal atmosphere. After 7 days incubation, 30 µl of a lamar blue solution was added to each well, and the plate was re-incubated overnight. A change in color from blue (oxidized state) to pink (reduced) indicated the growth of bacteria, and the MIC was defined as the lowest concentration of drug that prevented this change in color.

### 4.7. Molecular Docking studies

See the supplementary information for the procedures and other details.

## Contributors

HSPR supervised the findings of this work and conceived the studies. He was involved in planning and supervised the work. RG and LNA were involved the synthesis. JM performed the in silico studies. HSPR aided in interpreting the results obtained from synthesis, in silico and in vitro studies. HSPR, RG, LNA and JM prepared the first draft of the manuscript. HSPR and JM revised the manuscript into its final version, to which all the authors gave approval.

## Declaration of competing interest

None

## Acknowledgment

We thank Professor D. Sriram. Ph.D. Senior Professor, Department of Pharmacy, Birla Institute of Technology & Science-Pilani, Hyderabad Campus, Jawahar Nagar, Hyderabad-500 078. for conducting anti-tuberculosis studies. We thank DST-FIST, UGC SAP DRS-2 and CIF, PU for facilities and spectral recordings. Dr. RG thanks CSIR for SRF and UGC-PU for JRF. Dr. JM thanks Sharda University, Greater Noida for support.

## Notes

### Competing Interest Statement

The authors have declared no competing interest.

## References

1 (a) Matthew, W. E., Snyder, S. A., Stockwel, B. R. Privileged scaffolds for library design and drug discovery. Curr. Opin. Chem. Biol. 2010, 14, 347–361. (b) Newman, D. J., Cragg, G. M., Snader, K. M. The influence of natural products upon drug discovery. Natural Product Reports. 2000, 17, 215–234.

2 (a) Edon,V., Smith, D. T., Njardarson, J. T. Analysis of the structural diversity, substitution patterns, and frequency of nitrogen heterocycles among US FDA approved pharmaceuticals: Mini perspective. J. Med. Chem. 2014, 57, 10257–10274. (b) Likhosherstov, A. M., Filippova, O. V., Peresada, V. P., Kryzhanovskii, S. A., Vititnova, M. B., Kaverina, N. V., Reznikov, K. M. Azacycloalkanes. XXXIV. Synthesis and w antiarrhythmic activity of 2-(2′-*R*-2′-hydroxyethyl)-1, 2, 3, 4-tetra-hydro-pyrrolo-[1,2-a]pyrazines. Pharm. Chem. J. 2003, 37, 6–9.

3 a) Kumar, V., Kaur, K., Gupta, G. K., Sharma, A. K. Pyrazole containing natural products: synthetic preview and biological significance. Eur. J. Med. Chem. 2013, 69, 735–753. (b) Katritzky, A. R., Rees, C. W. Comprehensive Heterocyclic Chemistry. Pergamon Press, 1984.

4 Reviews: (a) Hardy, C. R. The Chemistry of Pyrazolopyridines in Advances in Heterocyclic Chemistry. 1984, 36, 343–409, Academic Press. (b) Castillo, J.C. and Portilla, J., Recent advances in the synthesis of new pyrazole derivatives. Targets Heterocycl. Syst. 2018, 22, 194–223.

5 Review: K Dodiya, D., R Trivedi, A., B Kataria, V. and H Shah, V. Advances in the Synthesis of Pyrazolo[3,4-b] pyridines. Current Organic Chemistry. 2012, 16, 400–417.

6 Review: Patel, J. B., Malick, J. B., Salama, A. I. and Goldberg, M. E. Pharmacology of Pyrazolopyridines. Pharmacology Biochemistry and Behavior. 1985, 23, 675–680.

7 Review: Davies, L. P., Brown, D. J., Chow, S. C. and Johnston, G. A. Pyrazolo[3,4-*d*]pyrimidines, A New Class of Adenosine Antagonists. Neuroscience letters, 1983, 41, 189–193.

8 Review: Zheleznova, N. N., Sedelnikova, A. and Weiss, D. S. Function and Modulation of δ-Containing GABAA receptors. Psychoneuroendocrinology. 2009, 34, S67–S73.

9 Höhn, H., Polacek, I. and Schulze, E., 1973. Potential antidiabetic agents. Pyrazolo[3,4-b]pyridines. J. med. chem. 16, 1340–1346.

10 https://www.nature.com/articles/d41573-021-00067-x

11 Review: (a) Pitre, T., Su, J., Cui, S., Scanlan, R., Chiang, C., Husnudinov, R., Khalid, M.F., Khan, N., Leung, G., Mikhail, D. and Saadat, P., 2022. Medications for the treatment of pulmonary arterial hypertension: a systematic review and network meta-analysis. European Respiratory Review, 31(165): 220036; [DOI: 10.1183/16000617.0036-2022]. (b) Review: Evgenov, O.V., Pacher, P., Schmidt, P.M., Haskó, G., Schmidt, H.H. and Stasch, J.P., 2006. NO-independent stimulators and activators of soluble guanylate cyclase: discovery and therapeutic potential. Nature reviews Drug discovery, 5(9), pp.755–768.

12 (a) Satya, P., Gupta, M., Gupta, R., Loupy, A. Microwave assisted solvent-free synthesis of pyrazolo[3,4-*b*]quinolines and pyrazolo[3,4-*c*]pyrazoles using p-TsOH. Tetrahedron Lett. 2001, 42, 3827–3829 (b) Hao, Y., Xu, X. P., Chen, T., Zhao, L. L., Ji, S. J. Multicomponent approaches to 8-carboxylnaphthyl-functionalized pyrazolo[3,4-b]pyridine derivatives. Org. Biomol. Chem. 2012, 10, 724–729.

13 Shi, D. Q., Yao, H., Shi, J. W. Three Component, One Pot Synthesis of pyrazolo[3,4-*b*]pyridine derivatives in aqueous media. Synth. Commun. 2008, 38, 1662–1669. (b) Barreiro,E. J., Camara, Celso, A., Verli, H., Brazil-Ma, L., Castro,N. G., Cintra,W. M., Aracava, Y., Rodrigues, C. R., Fraga, C. A. M., Design, Synthesis, and pharmacological profile of novel fused pyrazolo[4,3-*d*]pyridine and pyrazolo[3,4-*b*][1,8]naphthyridine isosteres: a new class of potent and selective acetylcholinesterase inhibitors. J. Med. Chem. 2003, 46, 1144–1152.

14 Ahmad, S., Seyyedhamzeh, M., Maleki, A., Behnam, M., Rezazadeh, F. Synthesis of fully substituted pyrazolo [3,4-*b*]pyridine-5-carboxamide derivatives via a one-pot four-component reaction. Tetrahedron Lett. 2009, 50, 2911–2913.

15 Kalsi, J. S., Rees, R. W., Hobbs, A. J., Royle, M., Kell, P. D., Ralph, D. J., Moncada, S., Cellek,S. BAY41-2272, a novel nitric oxide independent soluble guanylate cyclase activator, relaxes human and rabbit corpus cavernosum in vitro. The Journal of urology. 2003, 169, 761–766.

16 (a) Fabrizio, M., Schenone, S., Bondavalli, F., Brullo, C., Bruno, O., Ranise, A., Mosti, L. Synthesis and 3D QSAR of new pyrazolo[3,4-*b*]pyridines: potent and selective inhibitors of A1 adenosine receptors. J. Med. Chem. 2005, 48, 7172–7185. (b) Silvia, S., Bruno, O., Fossa,P., Ranise, A., Menozzi, G., Mosti, L., Bondavalli, F., Martini, CandTrincavelli, L. Synthesis and biological data of 4-amino-1-(2-chloro-2-phenylethyl)-1H-pyrazolo[3,4-*b*] pyridine-5-carboxylic acid ethyl esters, a new series of A 1-adenosine receptor (A1 AR) ligands. Bioorg. Med. Chem. Lett. 2001, 11, 2529–2531.

17 Nicole, H. J., Angell, T., Ballantine, S. P., Cook, C. M., Cooper, A. W., Dawson, J., Delves, C. J. Pyrazolopyridines as a novel structural class of potent and selective PDE4 inhibitors. Bioorg. Med. Chem. Lett. 2008, 18, 4237–4241.

18 (a) Jason, W., Bordas, V., Gaiba, A., Garton, N. S., Naylor, A., Rawlings, A. D., Slingsby, B. P., Smith, D. G., Takle, A. K., Ward, R. W. 6-Aryl-pyrazolo[3,4-*b*] pyridines: potent inhibitors of glycogen synthase kinase-3 (GSK-3). Bioorg. Med. Chem. Lett. 2003, 13, 3055–3057. (b) Mourad, C., Samadi, A., Soriano, E., Lozach, O., Meijer, L., Marco-Contelles. J. Synthesis and biological evaluation of 3,6-diamino-1*H*-pyrazolo[3,4-*b*]pyridine derivatives as protein kinase inhibitors. Bioorg. Med. Chem. Lett. 2009, 16, 4566–4569.

19 Laszlo, R., Blum, E., Padova, F. E., Buhl, T., Feifel, R., Gram, H., Hiestand, P., Manning, U., Neumann, U., Rucklin, G. Pyrazoloheteroaryls Novel p38α MAP kinase inhibiting scaffolds with oral activity. 2006, 16, 262–266.

20 Recent examples: (a) Maqbool, M., Rajvansh, R., Srividya, K. and Hoda, N., Deciphering the Robustness of Pyrazolo-Pyridine Carboxylate Core Structure-Based Compounds for Inhibiting Α-Synuclein In Transgenic C. *elegans* model of Synucleinopathy. Bioorganic & Medicinal Chemistry. 2020, 28, 17–115640. (b) Li, C., Zhang, F. and Shen, Z., An Efficient Domino Strategy for Synthesis of Novel Spirocycloalkane Fused Pyrazolo[3,4-*b*]pyridine Derivatives. Tetrahedron, 2020, 76, 131727, 1–8. (c) F. Zhang, C. Li, C. Qi. A One-Pot Three-Component Strategy for Highly Diastereoselective Synthesis of Spirocycloalkane Fused Pyrazolo[3,4-*b*]pyridine Derivatives Using Recyclable Solid Acid as a Catalyst. Org. Chem. Front. 2020, 7, 2456–2466. (d) Ji, Y., Li, L., Zhu, G., Zhou, Y., Lu, X., He, W., Gao, L. and Rong, L. Efficient Reactions for the Synthesis of Pyrazolo[3,4-*b*]pyridine and pyrano[2, 3-c]pyrazole Derivatives from N-Methyl-1-(methylthio)-2-nitroethen-1-amine. J. Heterocycl. Chem. 2020, 1–16. (e) Nafie, M., A, A. M., M, AK. Discovery of Novel Pyrazolo[3,4-*b*]pyridine Scaffold-Based Derivatives as Potential PIM-1 Kinase Inhibitors in Breast Cancer MCF-7 cells. Bioorg. Med. Chem. 2020, 28, 115828. (f) Jia, Q., Zhuo, C., X, L., Q, L, G, H., and Q, Li. Discovery of Novel Pyrazolo[3,4-*b*]pyridine Derivatives with Dual Activities of Vascular Remodelling Inhibition and Vasodilatation for the Treatment of Pulmonary Arterial Hypertension. J. Med. Chem. 2020, 63, 11215–11234.

21 Review: Kumar, S.V., Muthusubramanian, S. and Perumal, S. Recent Progress in the Synthesis of Pyrazolopyridines and Their Derivatives. Organic Preparations and Procedures International. 2019, 51, 1–89.

22 Bare, T. M., McLaren, C. D., Campbell, J. B., Firor, J. W., Resch, J. F., Walters, C. P., Salama, A. I., Meiners, B. A., Patel, J. B. Synthesis and structure-activity relationships of a series of anxioselectivepyrazolopyridine ester and amide anxiolytic agents. J. Med. Chem. 1989, 32, 2561–2573.

23 Wang, H. Y., Shi, D. Q. Three component one pot synthesis of pyrazolo[3,4-*b*]quinolin-5 (6*H*)-one derivatives in aqueous media. J. Heterocyclic Chem. 2012, 49, 212–216.

24 Rao, H. S. P., Adigopula, L. N. and Ramadas, K. One-pot synthesis of densely substituted pyrazolo[3,4-*b*]-4,7-dihydropyridines. ACS Combinatorial Science. 2017, 19, 279–285.

25 (a) Review M. A. Metwally, E. Abdel-latif Versatile α-oxoketene dithioacetals and analogues in heterocycle synthesis. J. Sulfur Chemistry. 2004, 25, 359–379. (b) Review: Yokoyama, M., Togo, H., Kondo, S. Synthesis of Heterocycles from Ketene Dithioacetals. Sulfur Reports. 1990, 10, 23–47. (c) Rao, H. S. P., Sivakumar, S. Condensation of α-Aroylketene Dithioacetals 2-Hydroxyarylaldehydes Results in Facile Synthesis of a Combinatorial Library of 3-Aroylcoumarins. J. Org. Chem. 2006, 71, 8715–8723. (d) Rao, H. S. P., Sivakumar, S. Aroylketene dithioacetal chemistry: Facile synthesis of 4-aroyl-3-methylsulfanyl-2-tosylpyrroles from aroylketene dithioacetals and TosMIC. Beilstein J. Org. Chem. 2007, 3, 31–35.

26 (a) Patrick, M., Thuillier, A. Sulfur reagents in organic synthesis. Elsevier, 2013. (b) Chryssostomos, C., Dieter, K. A., Sulfur-Centered Reactive Intermediates in Chemistry and Biology. Springer Science & Business Media, 2013. 197. (c) Review: Dieter, R. K. Tetrahedron. 1986, 42, 3029. (d) Review: Junjappa, H., Ila, H., Ashokan, C. V. Tetrahedron. 1990, 46, 5423–5454. (e) Liu Qun,YangZhiyun. Progress on the Chemistry of alpha-Oxo Ketene Dithioacetals J. Chin. J. Org. Chem. 1992, 12, 225–232.

27 (a) Crofton, J., Horne, N. and Miller, F., 2009. Crofton’s Clinical Tuberculosis. Design. http://www.tbrieder.org/publications/books_english/crofton_clinical.pdf. (b) https://www.nobelprize.org/prizes/medicine/1905/koch/biographical/

28 Koch, A. and Mizrahi, V., 2018. Mycobacterium tuberculosis. Trends in Microbiology, 26, 555–556. doi: 10.1016/j.tim.2018.02.012.

29 Marrakchi, H., Lanéelle, M.A. and Daffé, M., 2014. Mycolic acids: Structures, biosynthesis, and beyond. Chemistry & biology, 21(1), pp.67–85.

30 (a) Devi, P. B., Jogula, S., Reddy, A. P., Saxena, S., Sridevi, J. P., Sriram, D. and Yogeeswari, P., 2015. Design of novel *Mycobacterium tuberculosis* pantothenate synthetase inhibitors: Virtual screening, synthesis and in vitro biological activities. Molecular Informatics, 34, 147–159. (b) Yang, Y., Gao, P., Liu, Y., Ji, X., Gan, M., Guan, Y., Hao, X., Li, Z. and Xiao, C., 2011. A discovery of novel *Mycobacterium tuberculosis* pantothenate synthetase inhibitors based on the molecular mechanism of actinomycin D inhibition. Bioorganic & Medicinal Chemistry Letters, 21, 3943–3946.

31 Singh, S. P., Naithani, R., Aggarwal, R., Prakash, O. Synthesis of Some Novel Fluorinated Pyrazolo[3,4-*b*] Pyridines. Synth. Commun. 2004, 34, 4359–4367.

32 Williams, R. pKa Data Compiled by R. Williams. Available online: https://organicchemistrydata.org/hansreich/resources/pka/pka_data/pka-compilation-williams.pdf (accessed on 15th August 2021).

33 Kim, D. K., Gam, J., Kim, Y. W., Lim, J., Kim, H. T. and Kim, K. H. Synthesis and anti-HIV-1 activity of a series of 1-alkoxy-5-alkyl-6-(arylthio) uracils. J. Med. Chem. 1997, 40, 2363–2373.

34 (a) Review: Rentner, J., Kljajic, M., Offner, L., & Breinbauer, R. Recent Advances and Applications of Reductive Desulfurization in Organic Synthesis. Tetrahedron. 2014, 70, 8983–9027. (b) Griffin, P. J., Fava, M. A., Whittaker, S. J. T., Kolonko, K. J., & Catino, A. J. Synthesis of Tetraarylmethanes via a Friedel-Crafts Cyclization/DesulfurizationSstrategy. Tetrahedron Letters. 2018, 59, 3999–4002.

35 An example for use of LiOH for ester hydrolysis: Hu, L., Ren, Q., Deng, L., Zhou, Z., Cai, Z., Wang, B. and Li, Z., E. J. Med. Chem. 2020, 113106.

36 See supplementary information.

37 Suresh, A., Srinivasarao, S., Khetmalis, Y. M., Nizalapur, S., Sankaranarayanan, M. and Sekhar, K. V. G. C. Inhibitors of pantothenate synthetase of Mycobacterium tuberculosis–a medicinal chemist perspective. RSC Advances, 2020 10, 37098–37115.

38 Activity studies against MTB were carried out by Dr. D. Sriram, BITS Pilani.

39 a) J. A. Riddick, W. B. Bunger. Organic Solvents: Physical Properties and Methods of Purification; Techniques of Chemistry, Vol II, 1970, Wiley-Interscience, New York. b) J. F. Coetzee. Purification of Solvents, Pergamon Press, Oxford, 1982.

