## Supplementary Data for "Design, Synthesis, Molecular Docking and Biological Activity of Pyrazolo[3,4-*b*]pyridines as Promising Lead Candidates Against *Mycobacterium tuberculosis*"

SUPPLIMENTORY INFORMATION

Contents:

| **Entry** | **Title** | **Pages** |
| --- | --- | --- |
| 1. | Materials and Methods for Molecular Docking | 1-2 |
| 2. | Computed ADMET properties of **16kA** and **16kB** | 4 |
| 3. | Computed Lipinski properties of **16kA** and **16kB** | 5 |
| 4. | Computed ADMET properties of rest of the pyrazolopyridines | 6-11 |
| 5. | Computed Lipinski properties of rest of the pyrazolopyridines | 12-17 |
| 6. | Spectra (^1^H, ^13^C, DEPT-135, 2D (selected) NMR) and HRMS output of the pyrazolopyridines | 18-131 |

**Materials and Methods**

**Molecular Docking**

Molecular docking is a computational approach to understand the interaction by the residues at the active site of drug target proteins. Intermolecular interactions, binding orientation, mode of binding, estimated binding free energy (*ΔG*) of ligand towards drug target proteins are expected to emerge from this study. For molecular docking studies we employed the Pantothenate Synthetase from *M. tuberculosis*. Initially, the X-ray crystal structure of drug target protein was retrieved from Research Collaboratory Structural Bioinformatics Protein Data Bank (RCSB-PDB) ([www.rcsb.org](http://www.rcsb.org/)).^[1]^ Molecular docking studies were performed using Auto Dock or Molecular Graphics Tools.^[2]^ The protein preparation step comprised of adding polar hydrogens, merging the non-polar hydrogens, adding the Kollmann charges and finally recording it into the PDBQT (XYZ coordinates + Partial charges + Atom type) format. The energy minimized three-dimensional structure of the ligand molecules was subsequently subjected to Auto Dock Tools for ligand preparation which included adding the polar hydrogens, merging the non-polar hydrogens, adding Gasteiger charges and finally saved in to the PDBQT format. Once protein and ligand preparation steps were completed, the receptor grid box was created on the binding site of drug target protein. The receptor grid map was generated based on reported known-drug binding sites. Site specific or direct molecular dockings were performed between our ligands and target protein using Auto Dock. For the molecular docking calculations, the Lamarckian Genetic Algorithm (LGA) was employed with the input parameters that included docking runs: 100; population size: 150; maximum number of energy evaluations: 2,500,000; maximum number of generations: 27,000; mutation rate: 0.02 and cross-over rate: 0.8. Best docking complexes were chosen based on lowest free energy of binding, largest cluster (higher number of docking solutions in the cluster), key intermolecular interactions, etc. Finally, the molecular docking results of drug target protein with the ligand molecules were analyzed by programs like PyMOL^[3]^, MGL Tools and LigPlot.^[4]^

**Table 1. ADMET Properties of both the isomers (A and B) of the best pyrazolo[3,4-*b*]pyridine 16k**

| **S. No.** | **Compound**  **Structure** | **Compound No.** | **BBB** | **Consensus Log P** | **TPSA** | **Aqueous**  **Solubility**  **(AS)** | **CYP450 2D6**  **inhibitor** |
| --- | --- | --- | --- | --- | --- | --- | --- |
| 1 | 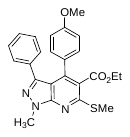 | **16kA**  C(6) SMe | No | 4.57 | 91.54 | Poorly Soluble | No |
| 2 | 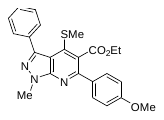 | **16kB**  C(4) SMe | No | 4.57 | 91.54 | Poorly Soluble | No |

**Table 2. Lipinski properties of both the isomers (A and B) of the best pyrazolo[3,4-*b*]pyridine 16k**

| **S. No.** | **Compound structure** | **Compound No.** | **Mol. Wt** | **H-bond donor** | **H-bond acceptor** | **Log P** | **Molecular Refractivity** |
| --- | --- | --- | --- | --- | --- | --- | --- |
| **1** | 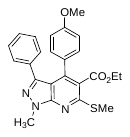 | **16kA**  (C(6) SMe | 433 | 0 | 5 | 5.019489 | 123.009483 |
| **2** | 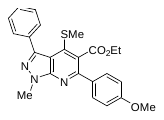 | **16kB**  (C(4) SMe) | 433 | 0 | 5 | 4.829479 | 122.348480 |

**Table 3**. ADMET properties of pyrazolo[3,4-*b*]pyridines

| **^S.No^** | **^Structure of Compound^** | **^BBB^** | **^Cytochrome P450 (Drug Metabolism Enzyme) Inhibition^** | | | | **^HIA^** | **^PGP inhibition^** | **^Pure_water_solubility^** |
| --- | --- | --- | --- | --- | --- | --- | --- | --- | --- |
|  |  |  | **^CYP_2C19^** | **^CYP_2C9^** | **^CYP_2D6^** | **^CYP_3A4^** |  |  |  |
| ^1^ | 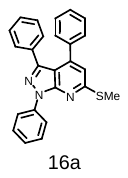 | ^2.16397^ | ^Non^ | ^Non^ | ^Non^ | ^Inhibitor^ | ^97.911067^ | ^Inhibitor^ | ^0.236853^ |
| ^2^ | 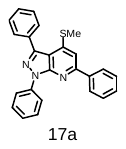 | ^2.94128^ | ^Non^ | ^Non^ | ^Non^ | ^Non^ | ^97.911067^ | ^Inhibitor^ | ^0.0123188^ |
| ^3^ | 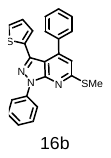 | ^5.42609^ | ^Non^ | ^Non^ | ^Non^ | ^Non^ | ^97.624570^ | ^Inhibitor^ | ^0.438614^ |
| ^4^ | 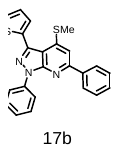 | ^2.94128^ | ^Non^ | ^Non^ | ^Non^ | ^Non^ | ^97.911067^ | ^Inhibitor^ | ^0.0123188^ |
| ^5^ | 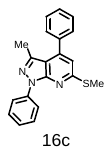 | ^3.0813^ | ^Non^ | ^Non^ | ^Non^ | ^Non^ | ^97.466166^ | ^Non^ | ^25.3508^ |
| ^6^ | 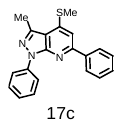 | ^3.25607^ | ^Non^ | ^Non^ | ^Non^ | ^Non^ | ^97.466166^ | ^Non^ | ^1.3185^ |
| ^7^ | 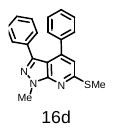 | ^1.94269^ | ^Non^ | ^Non^ | ^Non^ | ^Non^ | ^97.456377^ | ^Non^ | ^3.73165^ |
| ^8^ | 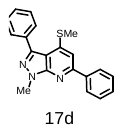 | ^3.12505^ | ^Non^ | ^Non^ | ^Non^ | ^Non^ | ^97.456377^ | ^Non^ | ^0.194085^ |
| ^9^ | 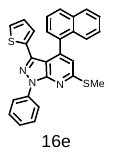 | ^6.11053^ | ^Non^ | ^Non^ | ^Non^ | ^Inhibitor^ | ^98.067708^ | ^Inhibitor^ | ^0.00906665^ |
| ^10^ |  | ^4.18803^ | ^Non^ | ^Non^ | ^Non^ | ^Inhibitor^ | ^98.067708^ | ^Inhibitor^ | ^0.00035576^ |
| ^11^ | 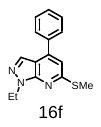 | ^2.30468^ | ^Non^ | ^Non^ | ^Non^ | ^Non^ | ^97.570153^ | ^Non^ | ^367.407^ |
| ^12^ | 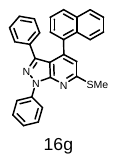 | ^2.40497^ | ^Non^ | ^Non^ | ^Non^ | ^Inhibitor^ | ^98.244428^ | ^Inhibitor^ | ^0.00490429^ |
| ^13^ | 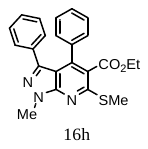 | ^3.55776^ | ^Inhibitor^ | ^Inhibitor^ | ^Non^ | ^Inhibitor^ | ^97.612600^ | ^Inhibitor^ | ^0.0290892^ |
| ^14^ | 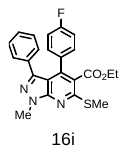 | ^3.10773^ | ^Inhibitor^ | ^Inhibitor^ | ^Non^ | ^Inhibitor^ | ^97.610917^ | ^Inhibitor^ | ^0.0137374^ |
| ^15^ | 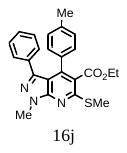 | ^4.29031^ | ^Inhibitor^ | ^Inhibitor^ | ^Non^ | ^Inhibitor^ | ^97.599082^ | ^Inhibitor^ | ^0.00851103^ |
| ^16^ | 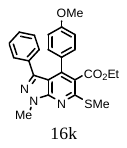 | ^2.01976^ | ^Inhibitor^ | ^Inhibitor^ | ^Non^ | ^Inhibitor^ | ^97.851379^ | ^Inhibitor^ | ^0.0193657^ |
| ^17^ | 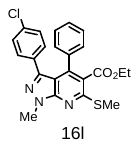 | ^4.64924^ | ^Inhibitor^ | ^Inhibitor^ | ^Non^ | ^Inhibitor^ | ^97.668685^ | ^Inhibitor^ | ^0.00406675^ |
| ^18^ | 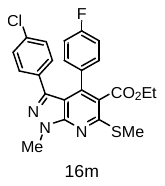 | ^3.99897^ | ^Inhibitor^ | ^Inhibitor^ | ^Non^ | ^Inhibitor^ | ^97.671534^ | ^Inhibitor^ | ^0.00191409^ |
| ^19^ | 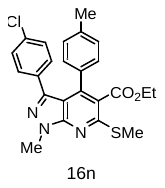 | ^5.06274^ | ^Inhibitor^ | ^Inhibitor^ | ^Non^ | ^Inhibitor^ | ^97.708048^ | ^Inhibitor^ | ^0.00118672^ |
| ^20^ | 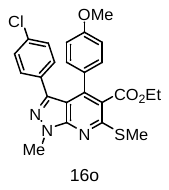 | ^2.93259^ | ^Inhibitor^ | ^Inhibitor^ | ^Non^ | ^Inhibitor^ | ^97.671148^ | ^Inhibitor^ | ^0.00269263^ |
| ^21^ | 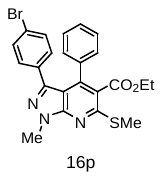 | ^4.75941^ | ^Inhibitor^ | ^Inhibitor^ | ^Non^ | ^Inhibitor^ | ^97.814741^ | ^Inhibitor^ | ^0.00256237^ |
| ^22^ | 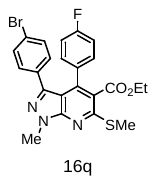 | ^4.03872^ | ^Inhibitor^ | ^Inhibitor^ | ^Non^ | ^Inhibitor^ | ^97.818587^ | ^Inhibitor^ | ^0.00120164^ |
| ^23^ | 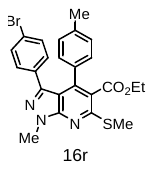 | ^5.10829^ | ^Inhibitor^ | ^Inhibitor^ | ^Non^ | ^Inhibitor^ | ^97.864766^ | ^Inhibitor^ | ^0.00074559^ |
| ^24^ | 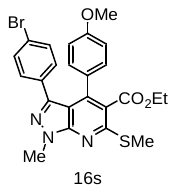 | ^3.00803^ | ^Inhibitor^ | ^Inhibitor^ | ^Non^ | ^Inhibitor^ | ^97.746264^ | ^Inhibitor^ | ^0.00168654^ |
| ^25^ | 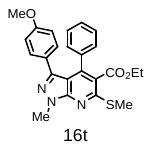 | ^2.5765^ | ^Inhibitor^ | ^Inhibitor^ | ^Non^ | ^Inhibitor^ | ^97.851379^ | ^Inhibitor^ | ^0.0193657^ |
| ^26^ | 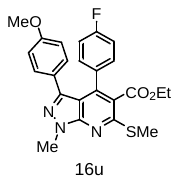 | ^2.53512^ | ^Inhibitor^ | ^Inhibitor^ | ^Non^ | ^Inhibitor^ | ^97.844521^ | ^Inhibitor^ | ^0.00911846^ |
| ^27^ | 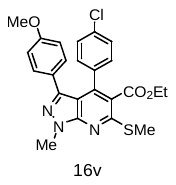 | ^3.4911^ | ^Inhibitor^ | ^Inhibitor^ | ^Non^ | ^Inhibitor^ | ^97.671148^ | ^Inhibitor^ | ^0.00330033^ |
| ^28^ | 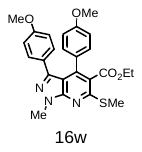 | ^1.63672^ | ^Inhibitor^ | ^Inhibitor^ | ^Non^ | ^Inhibitor^ | ^98.294713^ | ^Inhibitor^ | ^0.0128306^ |
| ^29^ | 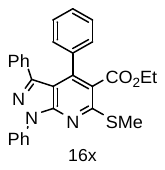 | ^4.37933^ | ^Inhibitor^ | ^Inhibitor^ | ^Non^ | ^Inhibitor^ | ^97.848547^ | ^Inhibitor^ | ^0.000307046^ |
| ^30^ | 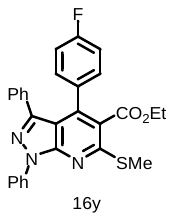 | ^2.36837^ | ^Inhibitor^ | ^Inhibitor^ | ^Non^ | ^Inhibitor^ | ^97.852462^ | ^Inhibitor^ | ^0.000144178^ |
| ^31^ | 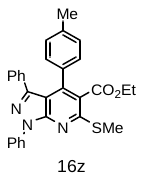 | ^3.71121^ | ^Inhibitor^ | ^Inhibitor^ | ^Non^ | ^Inhibitor^ | ^97.899173^ | ^Inhibitor^ | ^8.94345e-005^ |
| ^32^ | 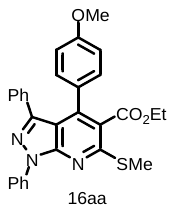 | ^2.79767^ | ^Inhibitor^ | ^Inhibitor^ | ^Non^ | ^Inhibitor^ | ^97.772349^ | ^Inhibitor^ | ^0.000202524^ |
| ^33^ | 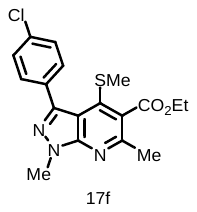 | ^2.08054^ | ^Inhibitor^ | ^Inhibitor^ | ^Non^ | ^Inhibitor^ | ^97.869957^ | ^Inhibitor^ | ^0.342381^ |
| ^34^ | 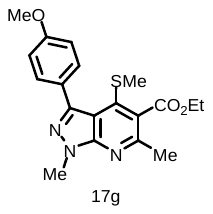 | ^0.608664^ | ^Inhibitor^ | ^Inhibitor^ | ^Non^ | ^Inhibitor^ | ^99.191916^ | ^Inhibitor^ | ^1.62766^ |
| ^35^ | 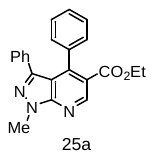 | ^3.6795^ | ^Inhibitor^ | ^Inhibitor^ | ^Non^ | ^Inhibitor^ | ^97.401359^ | ^Inhibitor^ | ^0.138991^ |
| ^36^ | 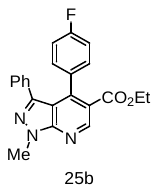 | ^2.30708^ | ^Inhibitor^ | ^Inhibitor^ | ^Non^ | ^Inhibitor^ | ^97.403484^ | ^Inhibitor^ | ^0.0660003^ |
| ^37^ | 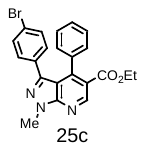 | ^3.31754^ | ^Inhibitor^ | ^Inhibitor^ | ^Non^ | ^Inhibitor^ | ^97.784027^ | ^Inhibitor^ | ^0.0403824^ |
| ^38^ | 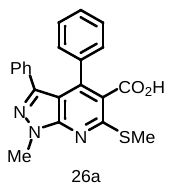 | ^2.29966^ | ^Non^ | ^Inhibitor^ | ^Non^ | ^Non^ | ^98.705409^ | ^Inhibitor^ | ^0.0999299^ |
| ^39^ | 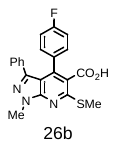 | ^2.14113^ | ^Non^ | ^Inhibitor^ | ^Non^ | ^Non^ | ^98.690963^ | ^Inhibitor^ | ^0.0473427^ |
| ^40^ | 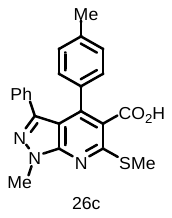 | ^3.09296^ | ^Non^ | ^Inhibitor^ | ^Non^ | ^Non^ | ^98.521832^ | ^Inhibitor^ | ^0.0293113^ |
| ^41^ | 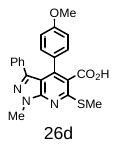 | ^1.05522^ | ^Non^ | ^Inhibitor^ | ^Non^ | ^Inhibitor^ | ^99.336844^ | ^Inhibitor^ | ^0.0668712^ |
| ^42^ | 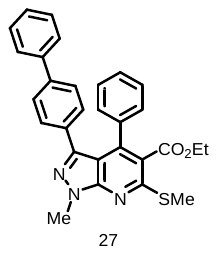 | ^0.652353^ | ^Inhibitor^ | ^Inhibitor^ | ^Non^ | ^Inhibitor^ | ^97.899173^ | ^Inhibitor^ | ^0.000363316^ |

**Table 4**. Lipinski properties of pyrazolo[3,4-*b*]pyridine derivatives

| S.No. | **Structure of the compound** | **Mol Wt (Da)** | **H-bond donar** | **H-bond acceptor** | **LogP** | **Molecular Refractivity** |
| --- | --- | --- | --- | --- | --- | --- |
| 1 | 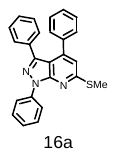 | 393 | 0 | 2 | 6.47 | 121.01 |
| 2 | 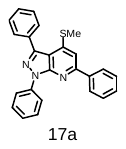 | 393 | 0 | 2 | 6.282 | 120.35 |
| 3 | 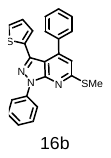 | 399 | 0 | 2 | 6.34 | 118.23 |
| 4 | 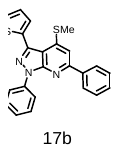 | 399 | 0 | 2 | 6.15 | 117.57 |
| 5 | 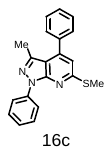 | 331 | 0 | 2 | 5.30 | 100.98 |
| 6 | 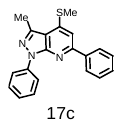 | 331 | 0 | 2 | 5.11 | 100.32 |
| 7 | 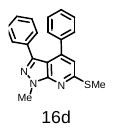 | 331 | 0 | 2 | 4.83 | 100.50 |
| 8 | 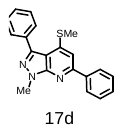 | 331 | 0 | 2 | 4.64 | 99.83 |
| 9 | 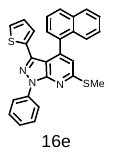 | 449 | 0 | 2 | 7.50 | 135.74 |
| 10 | 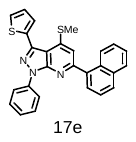 | 449 | 0 | 2 | 7.31 | 135.08 |
| 11 | 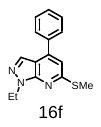 | 269 | 0 | 2 | 3.84 | 80.41 |
| 12 | 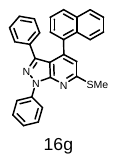 | 443 | 0 | 2 | 7.63 | 138.52 |
| 13 | 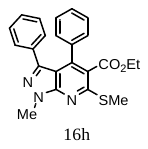 | 403 | 0 | 4 | 5.01 | 116.45 |
| 14 | 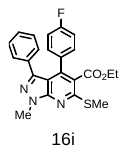 | 421 | 0 | 4 | 5.14 | 116.41 |
| 15 | 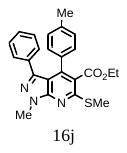 | 417 | 0 | 4 | 5.31 | 121.19 |
| 16 | 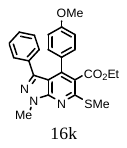 | 433 | 0 | 5 | 5.01 | 123.00 |
| 17 | 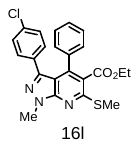 | 437 | 0 | 4 | 5.08 | 119.19 |
| 18 | 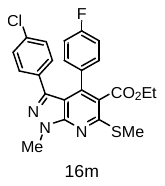 | 455 | 0 | 4 | 5.22 | 119.15 |
| 19 | 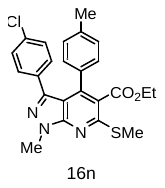 | 451.5 | 0 | 4 | 5.39 | 123.93 |
| 20 | 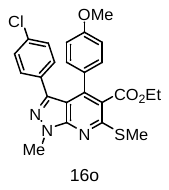 | 467.5 | 0 | 5 | 5.09 | 125.74 |
| 21 | 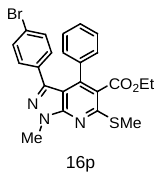 | 481 | 0 | 4 | 5.96 | 124.81 |
| 22 | 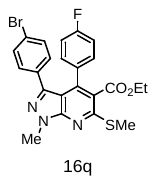 | 499 | 0 | 4 | 6.10 | 124.77 |
| 23 | 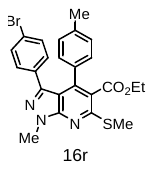 | 495 | 0 | 4 | 6.27 | 129.55 |
| 24 | 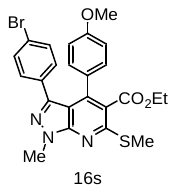 | 511 | 0 | 5 | 5.97 | 131.37 |
| 25 | 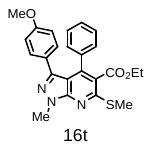 | 433 | 0 | 5 | 5.20 | 123.67 |
| 26 | 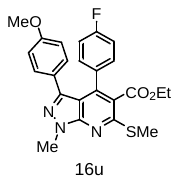 | 451 | 0 | 5 | 5.15 | 122.96 |
| 27 | 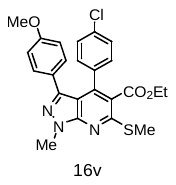 | 467.5 | 0 | 5 | 4.90 | 125.08 |
| 28 | 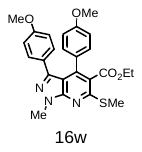 | 463 | 0 | 6 | 5.02 | 129.56 |
| 29 | 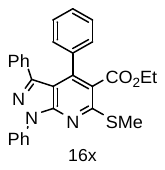 | 465 | 0 | 4 | 6.65 | 136.97 |
| 30 | 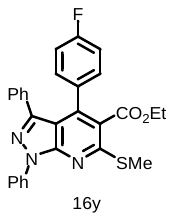 | 483 | 0 | 4 | 6.79 | 136.93 |
| 31 | 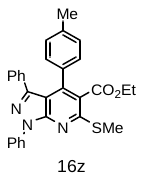 | 479 | 0 | 4 | 6.96 | 141.71 |
| 32 | 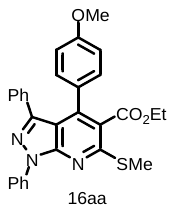 | 495 | 0 | 5 | 6.66 | 143.52 |
| 33 | 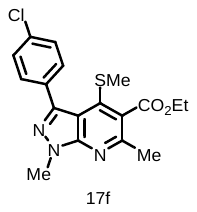 | 375.5 | 0 | 4 | 3.72 | 98.49 |
| 34 | 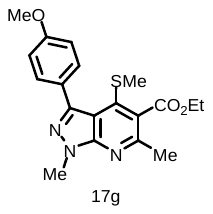 | 371 | 0 | 5 | 3.66 | 102.31 |
| 35 | 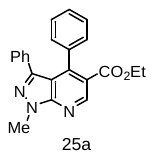 | 357 | 0 | 4 | 4.28 | 104.67 |
| 36 | 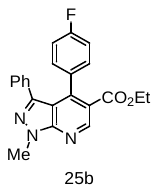 | 375 | 0 | 4 | 4.42 | 104.63 |
| 37 | 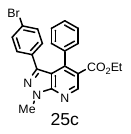 | 435 | 0 | 4 | 5.24 | 113.03 |
| 38 | 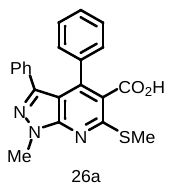 | 375 | 1 | 4 | 4.53 | 107.46 |
| 39 | 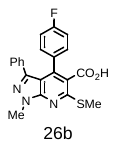 | 393 | 1 | 4 | 4.67 | 107.41 |
| 40 | 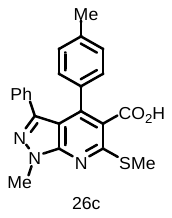 | 389 | 1 | 4 | 4.84 | 112.19 |
| 41 | 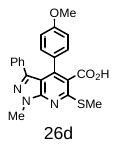 | 405 | 1 | 5 | 4.54 | 114.01 |
| 42 | 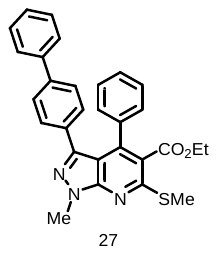 | 479 | 0 | 4 | 6.67 | 141.89 |

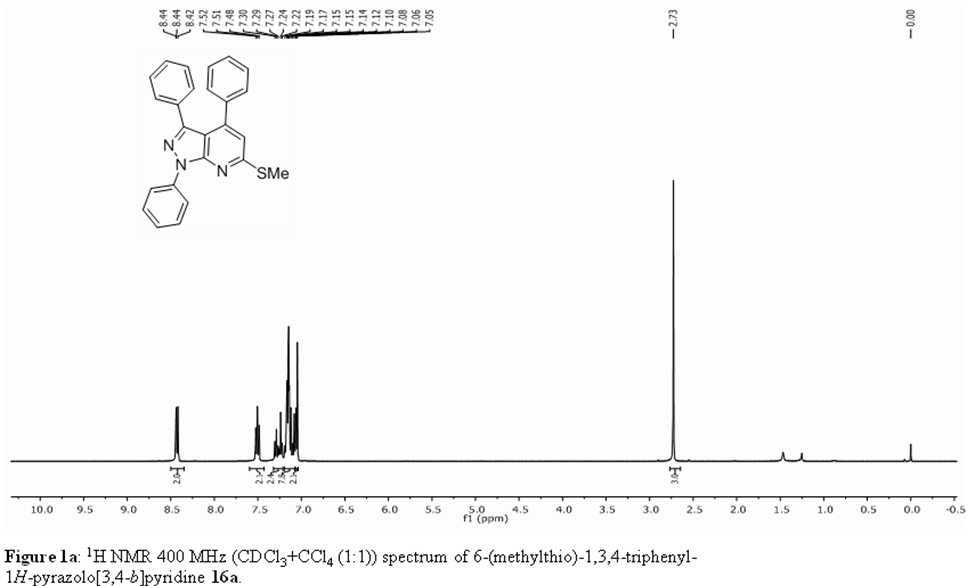

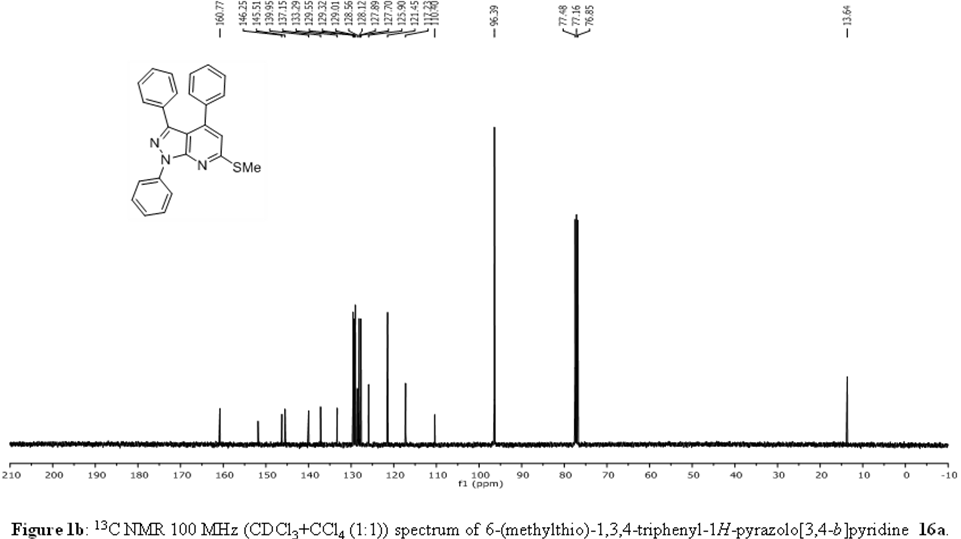

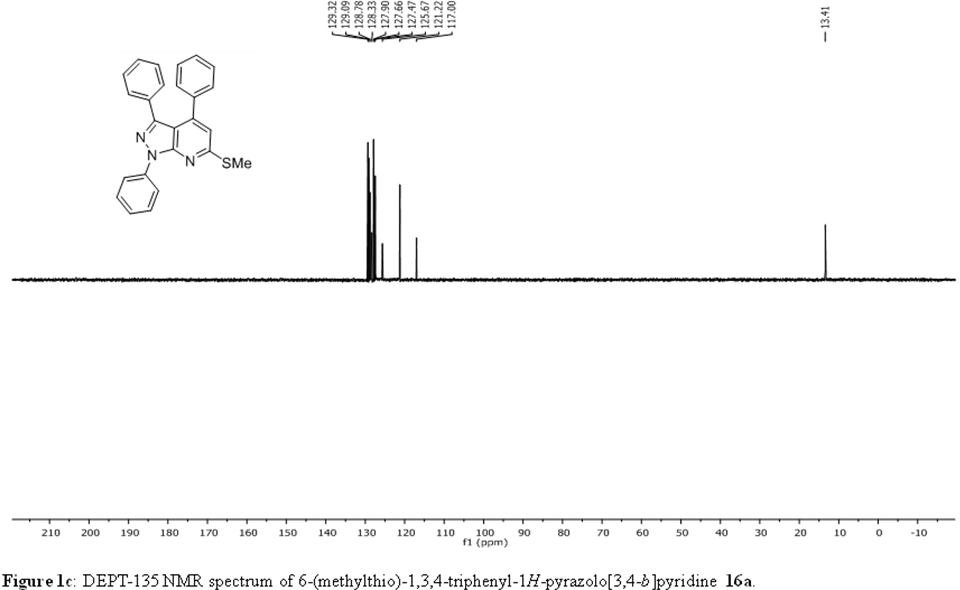

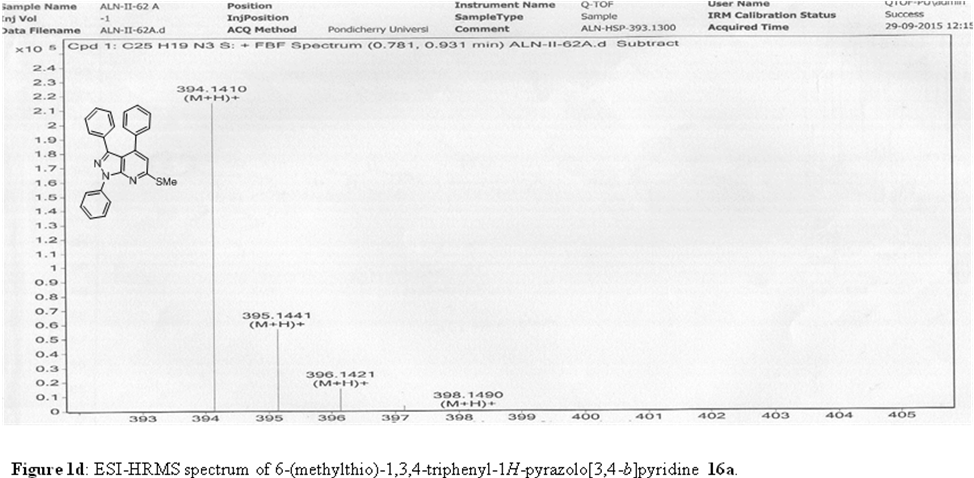

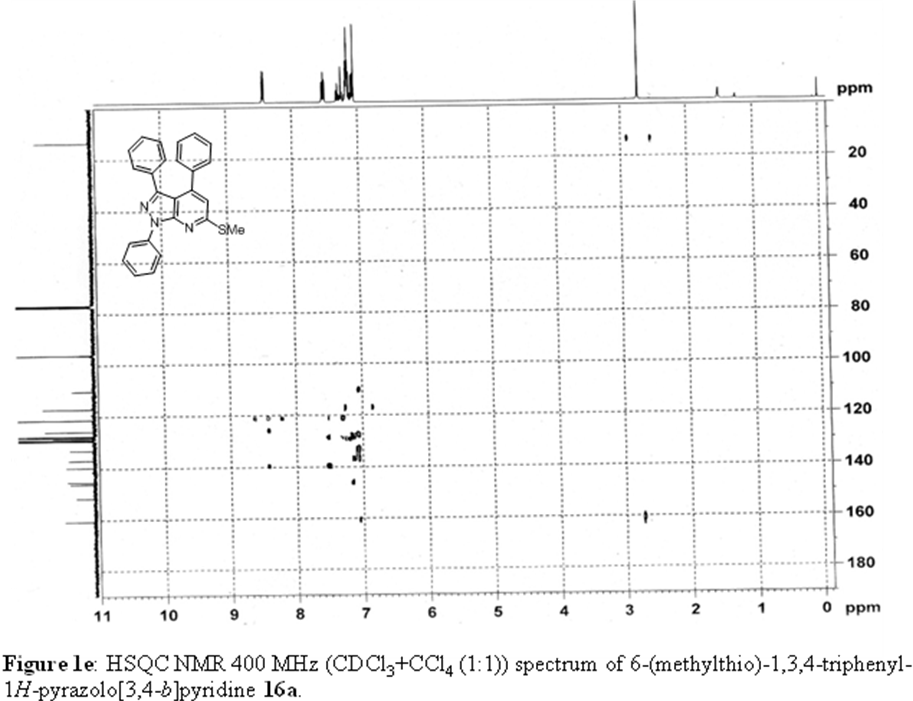

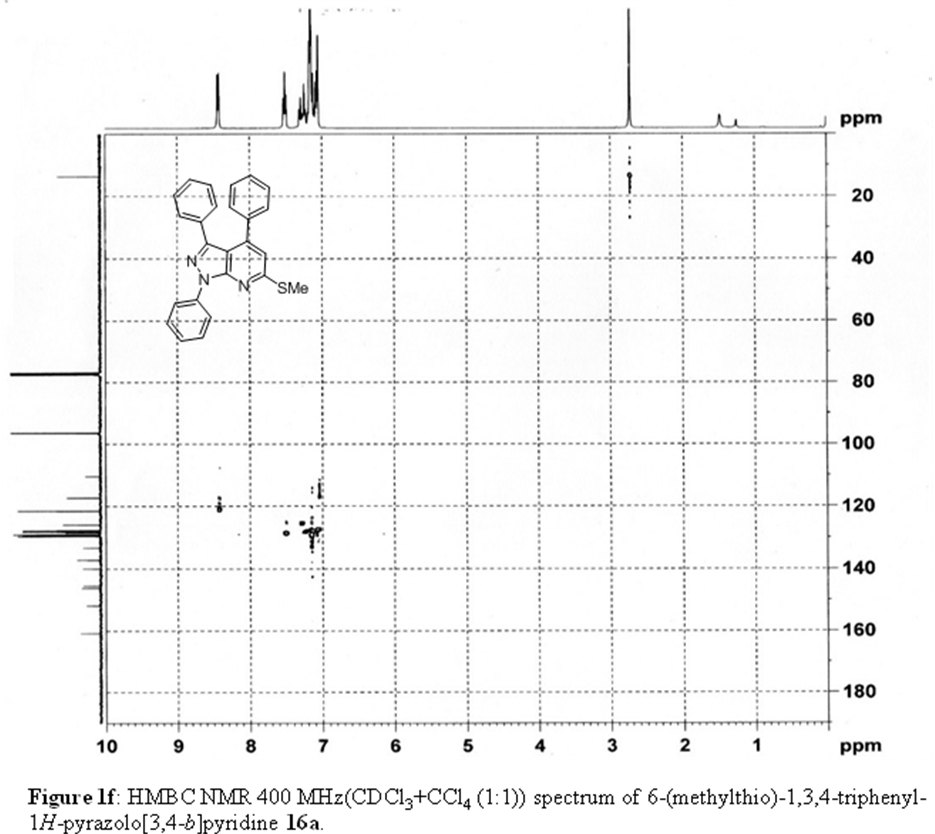

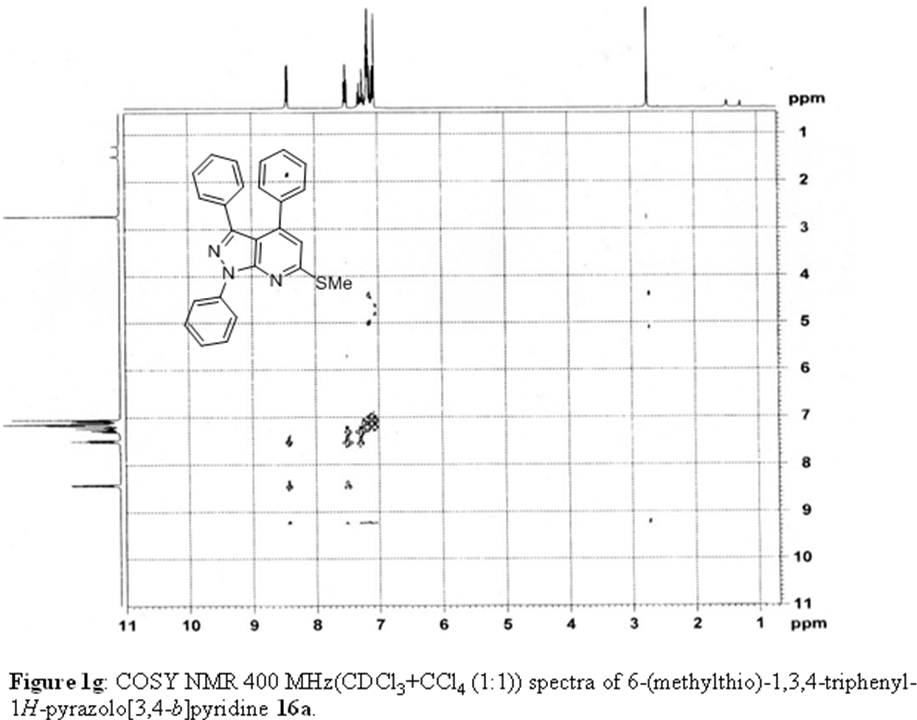

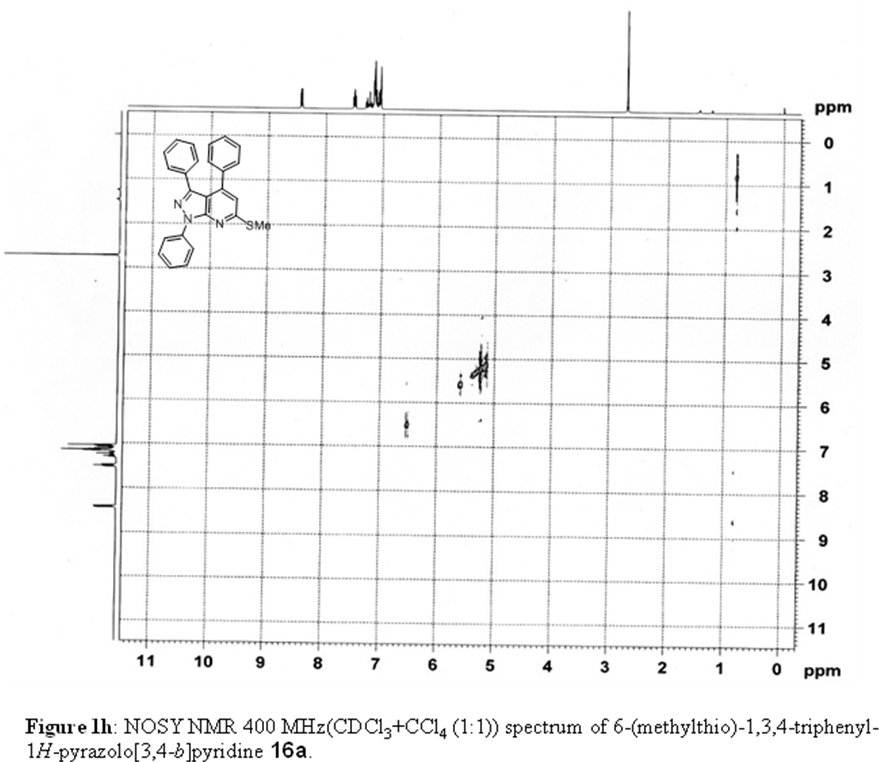

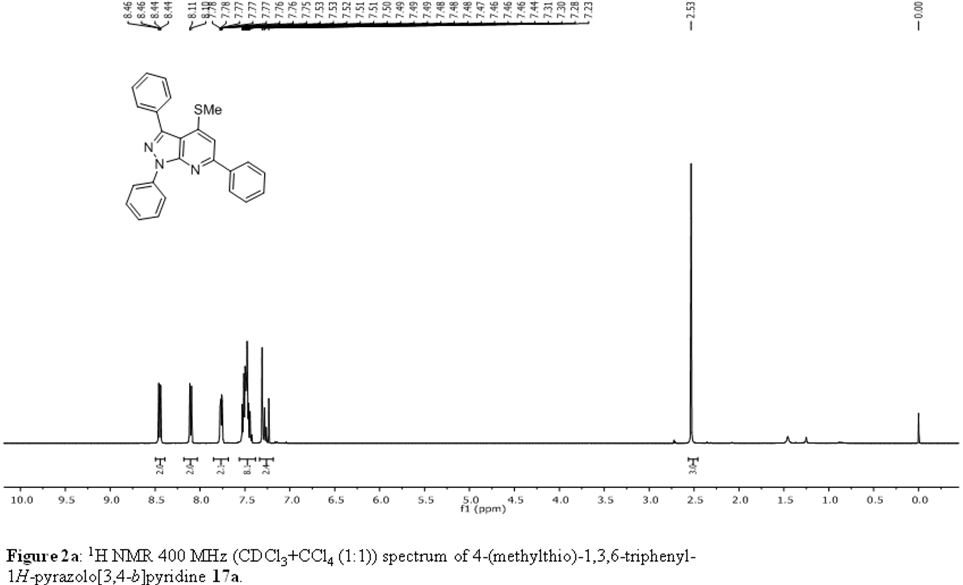

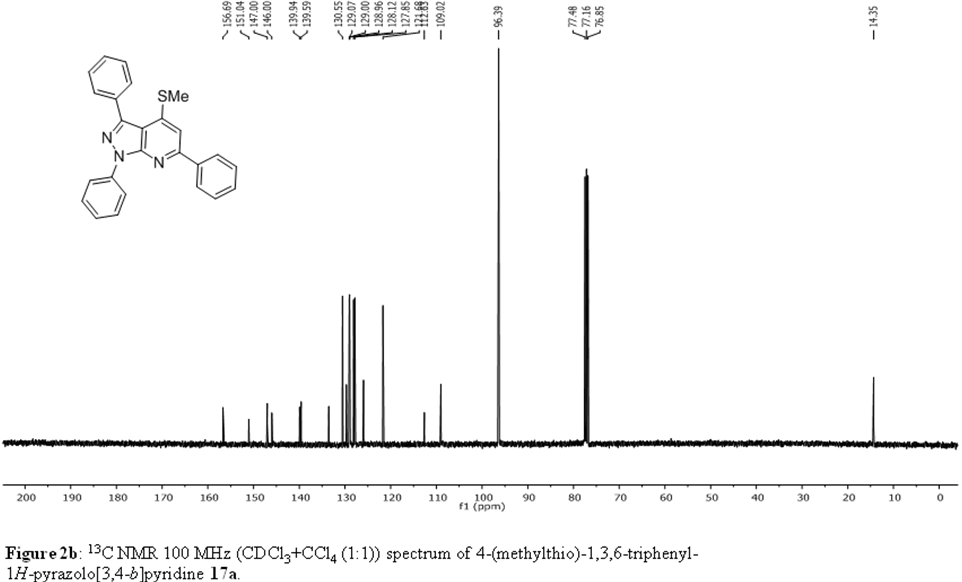

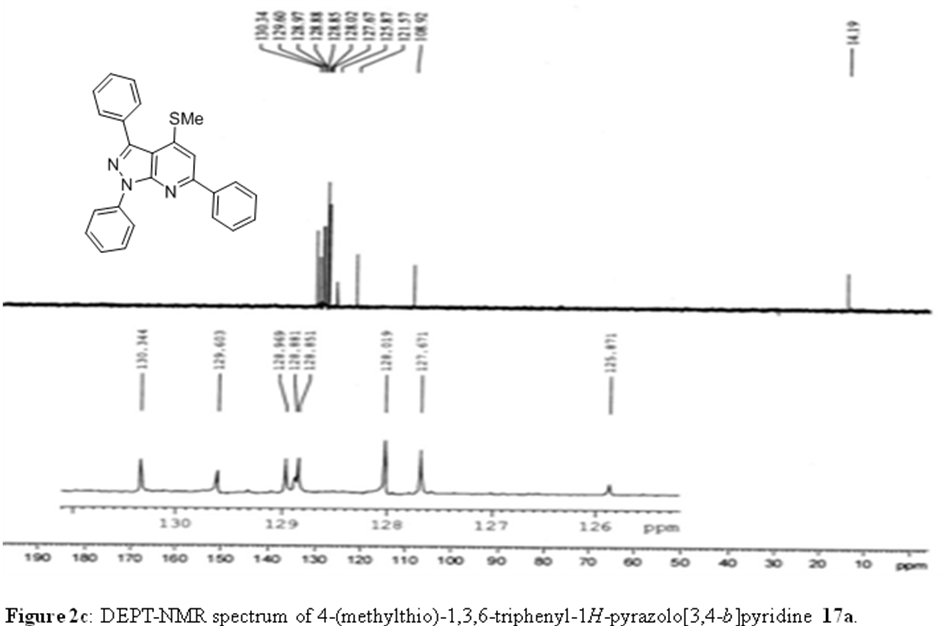

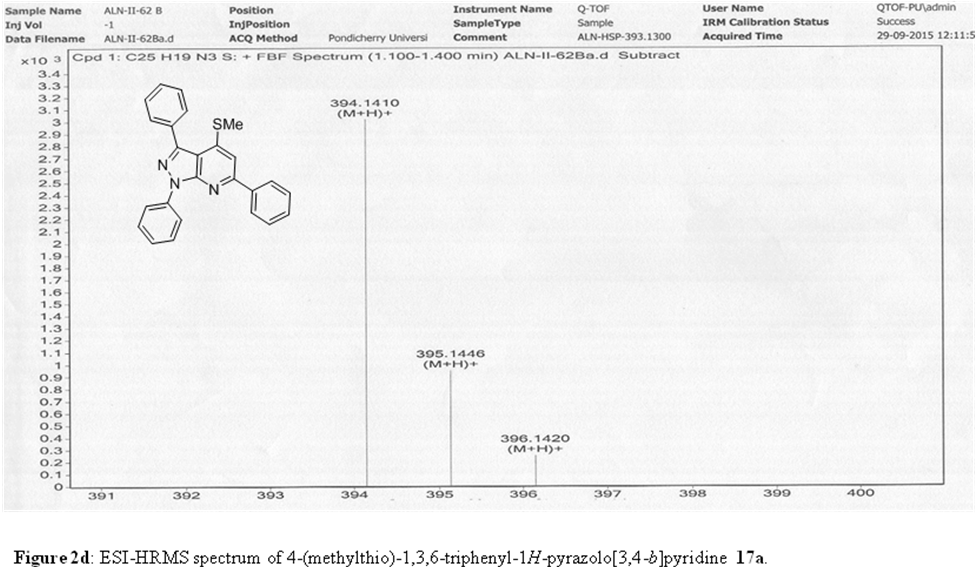

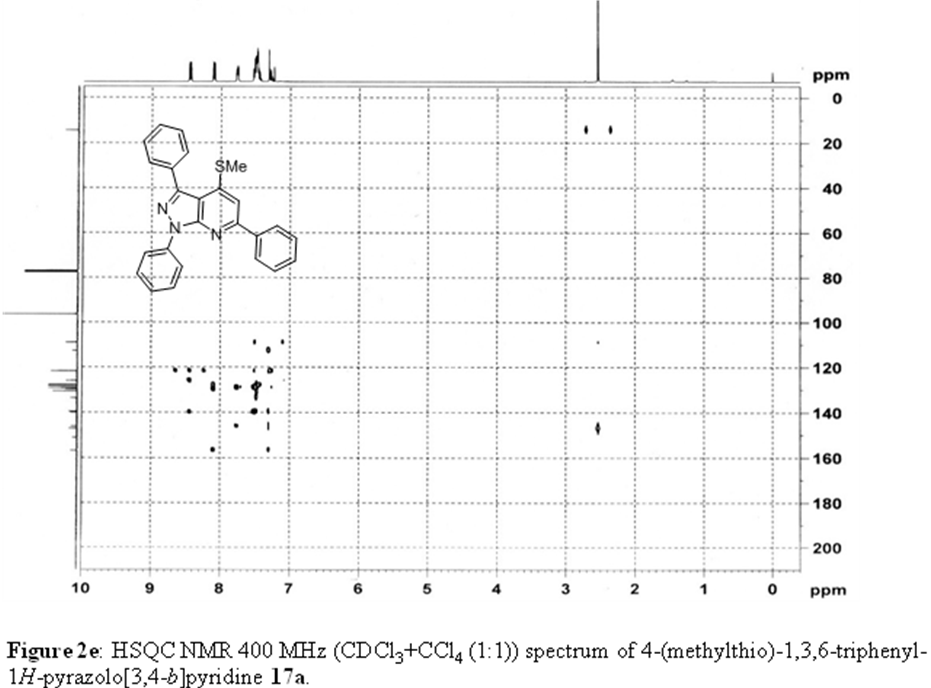

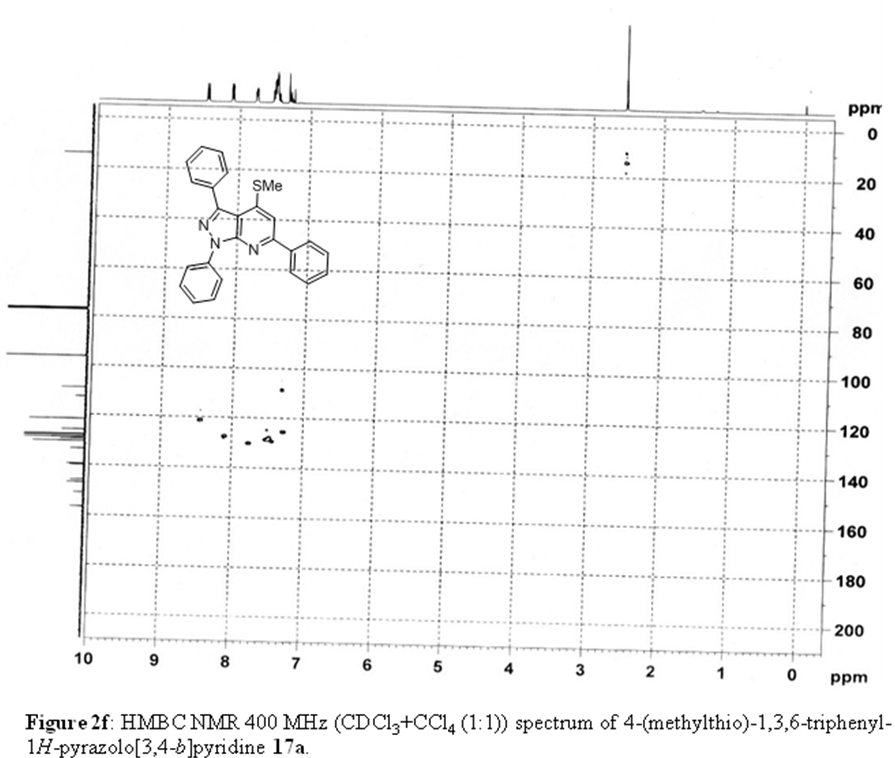

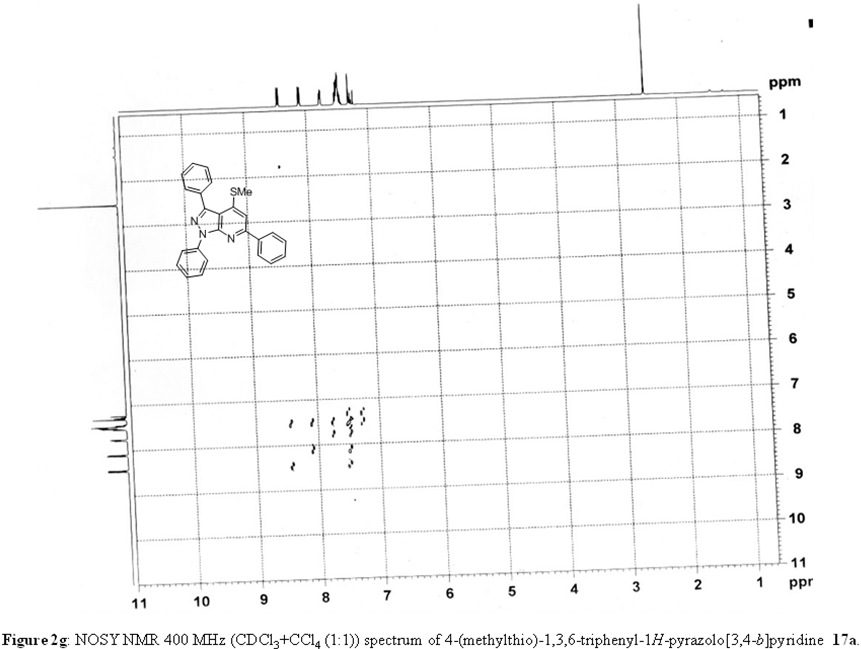

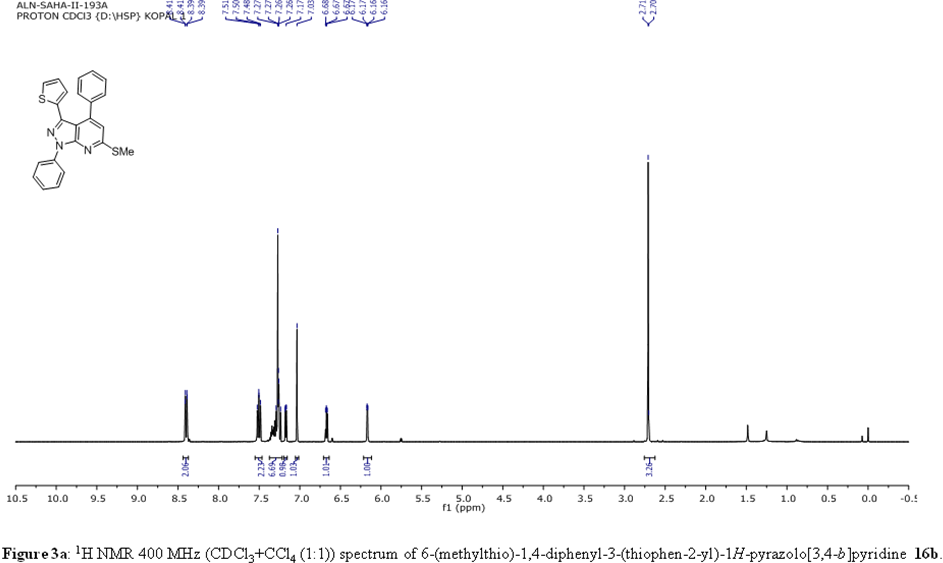

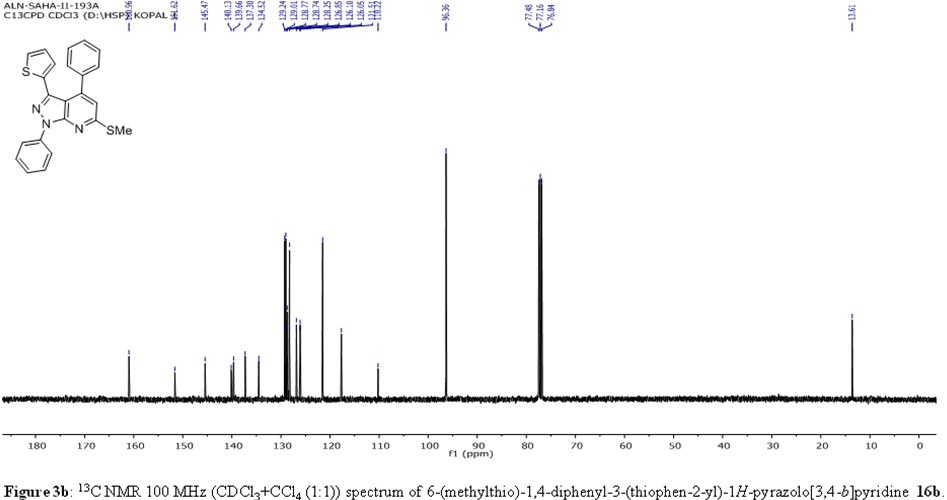

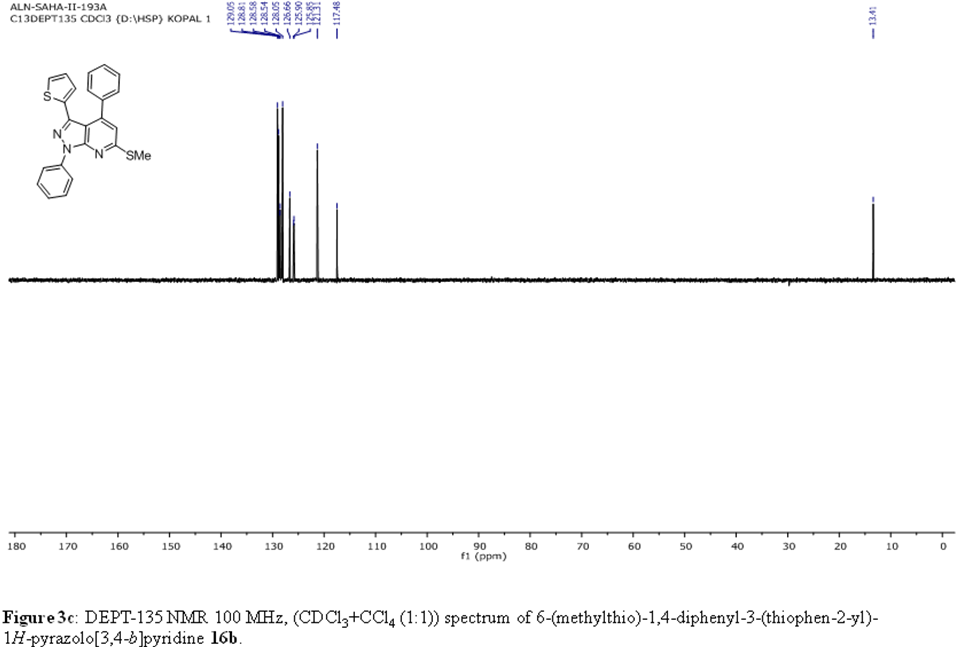

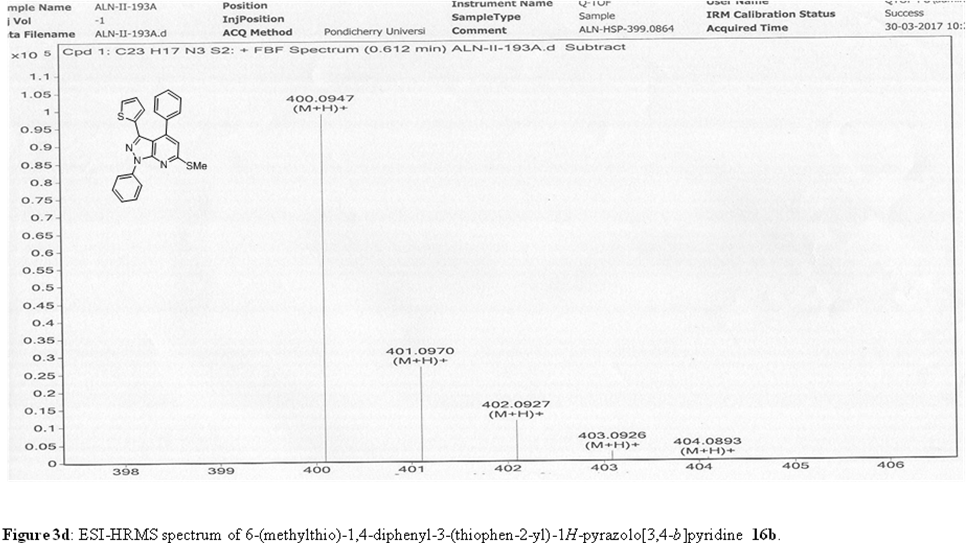

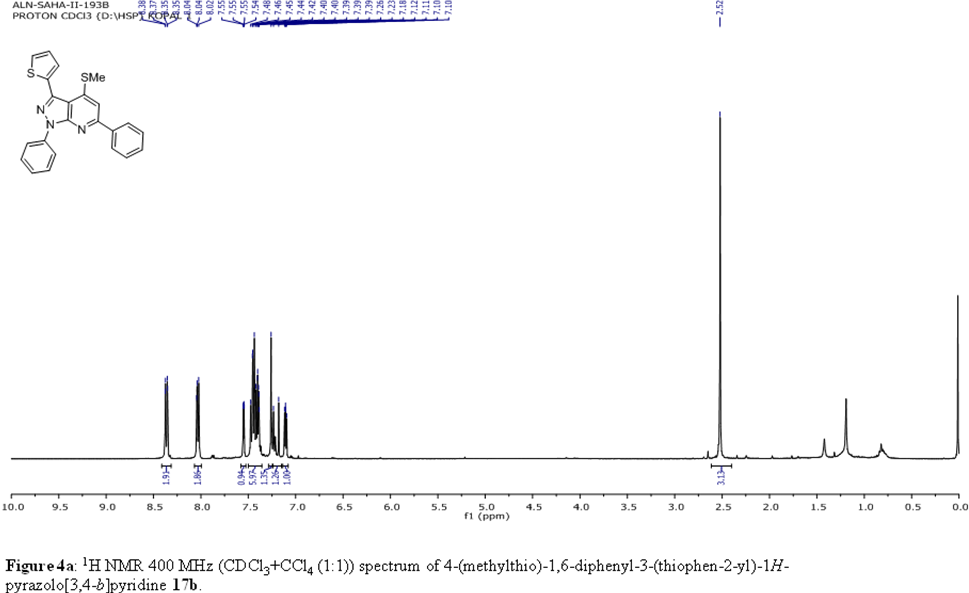

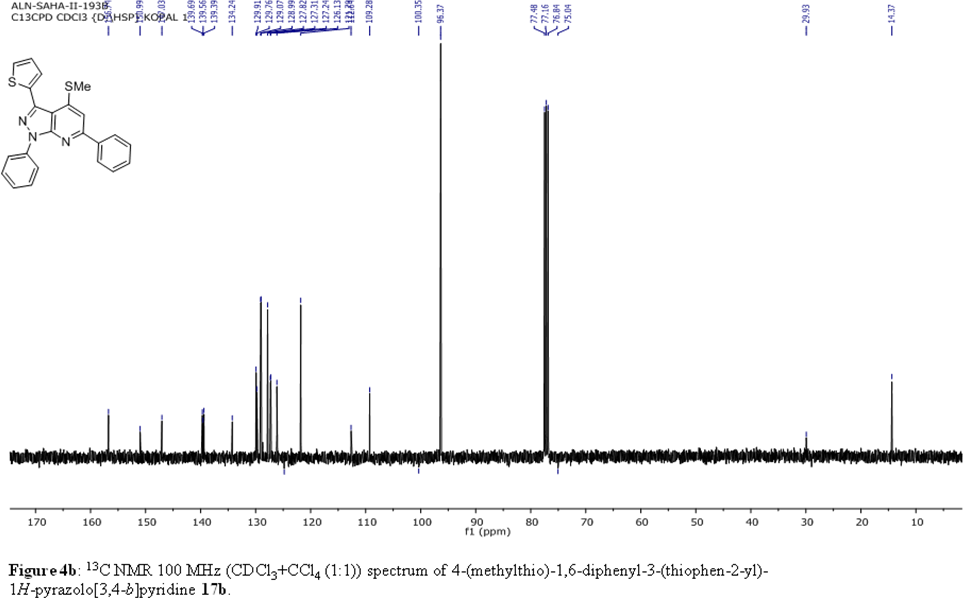

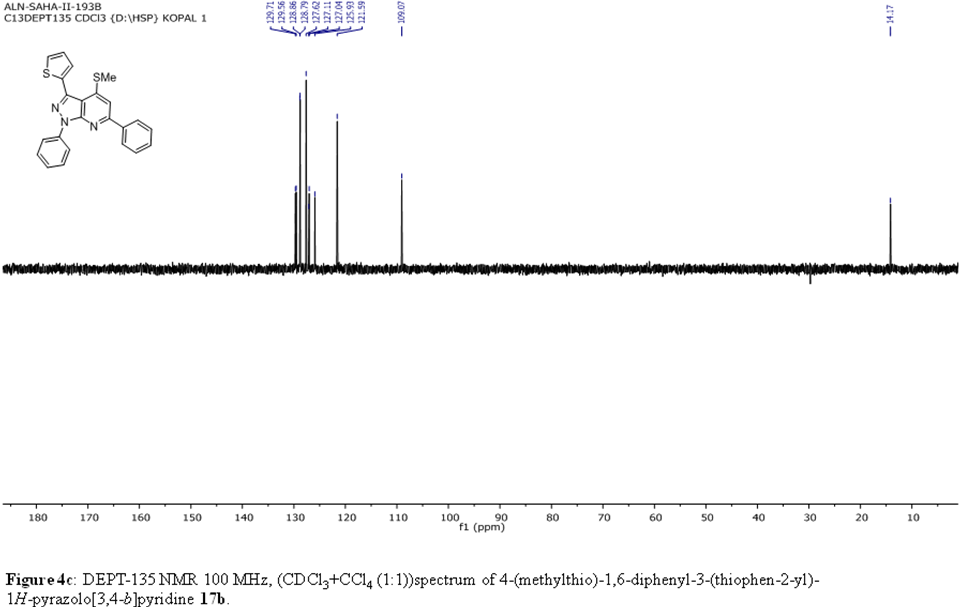

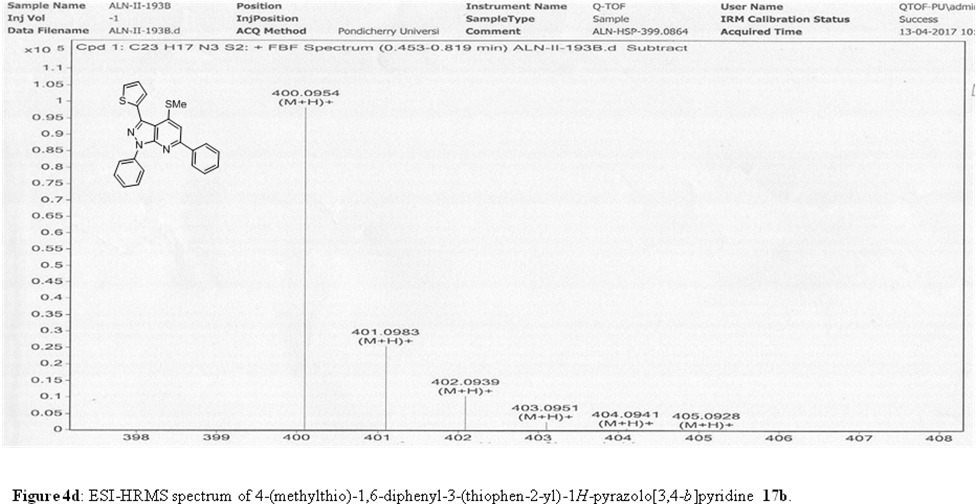

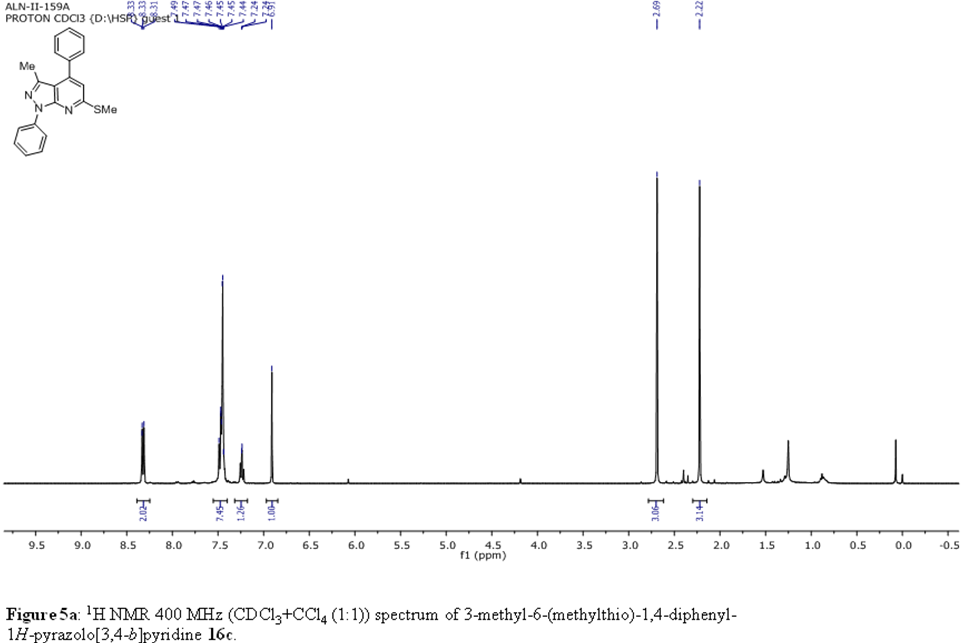

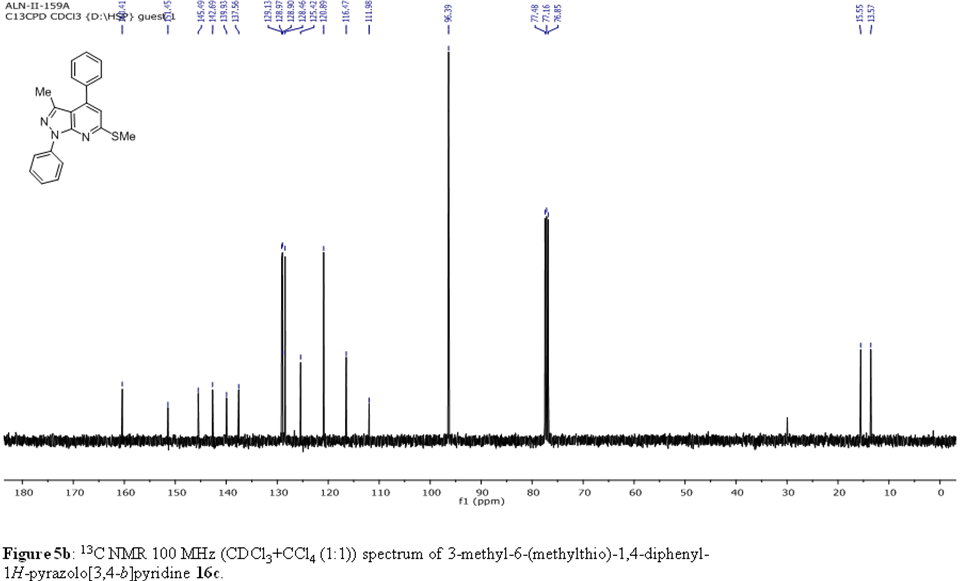

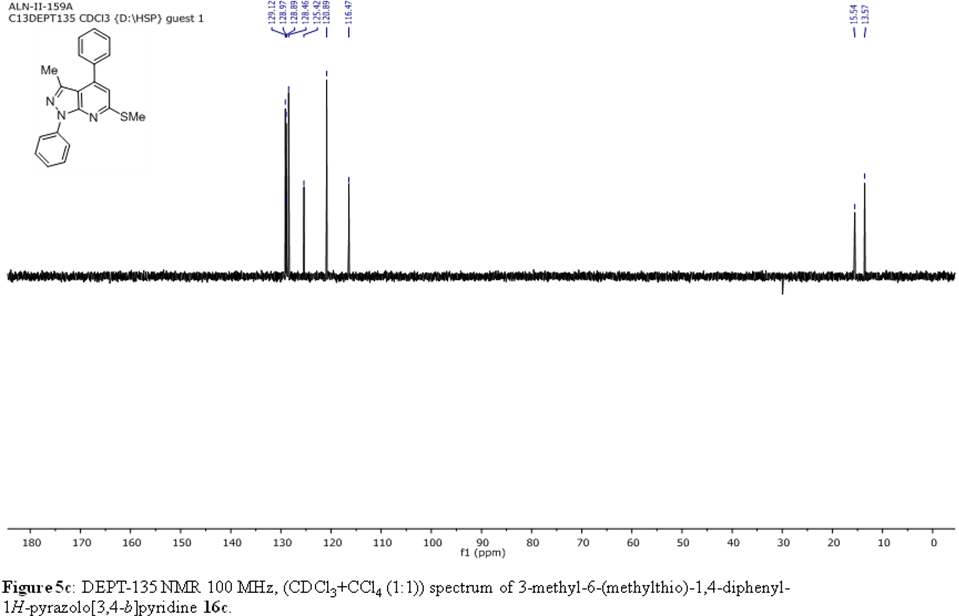

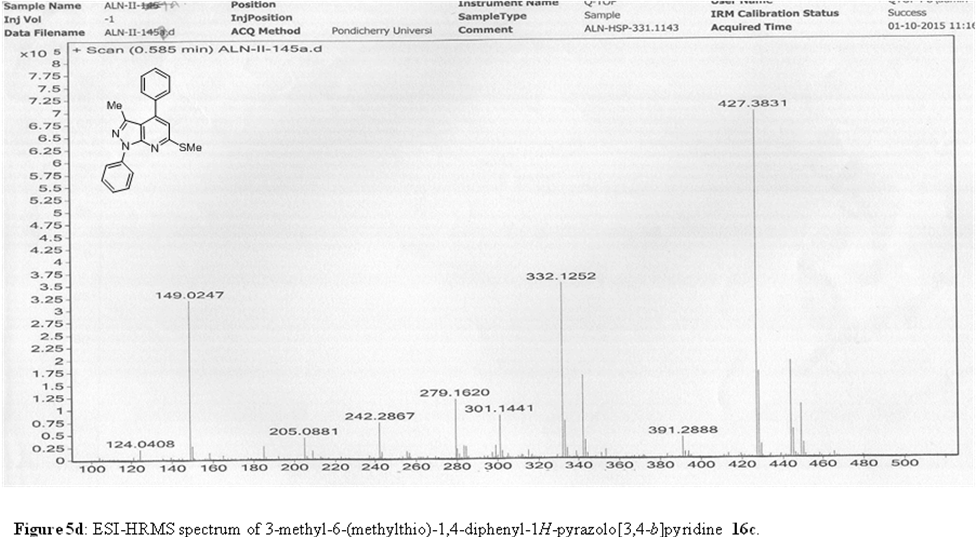

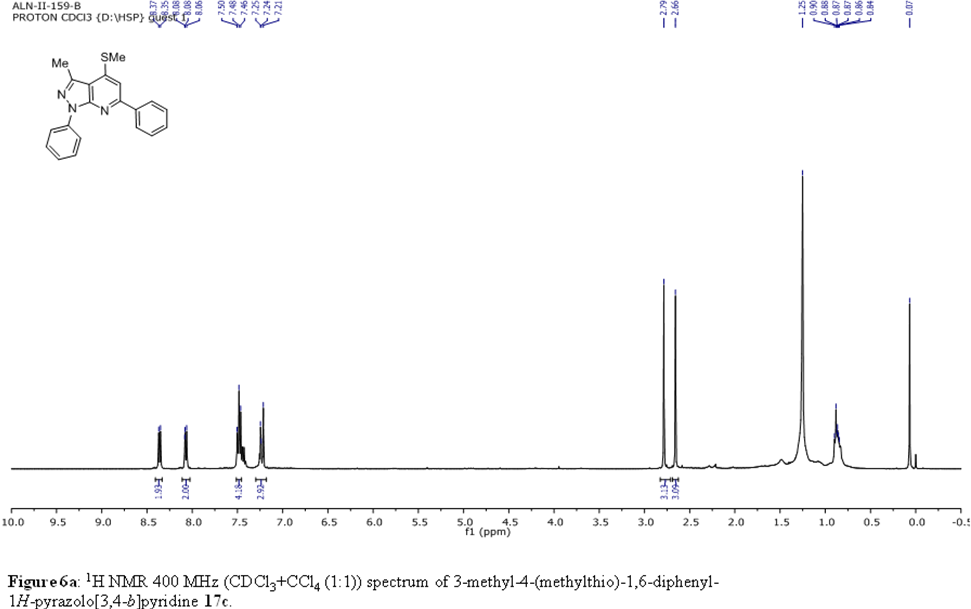

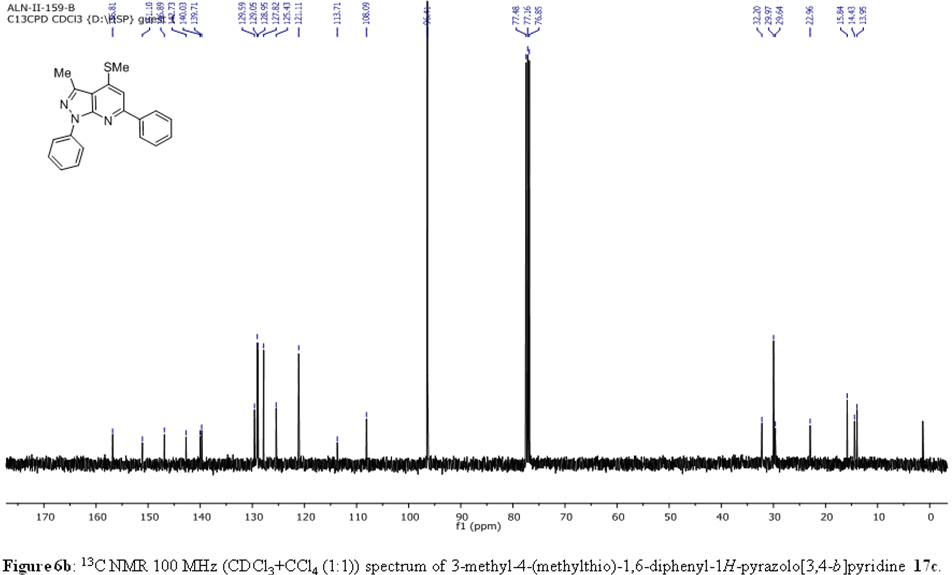

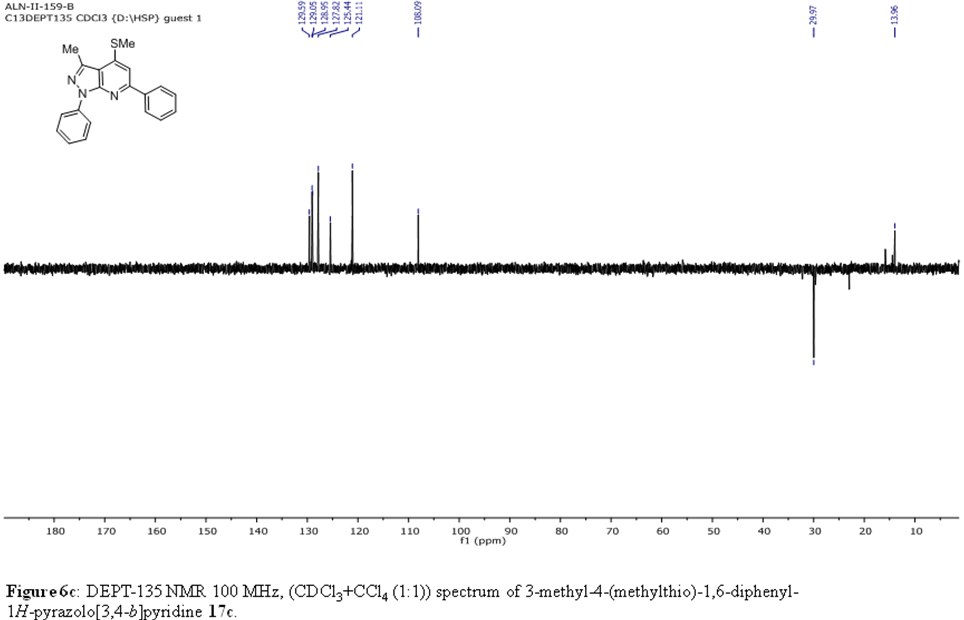

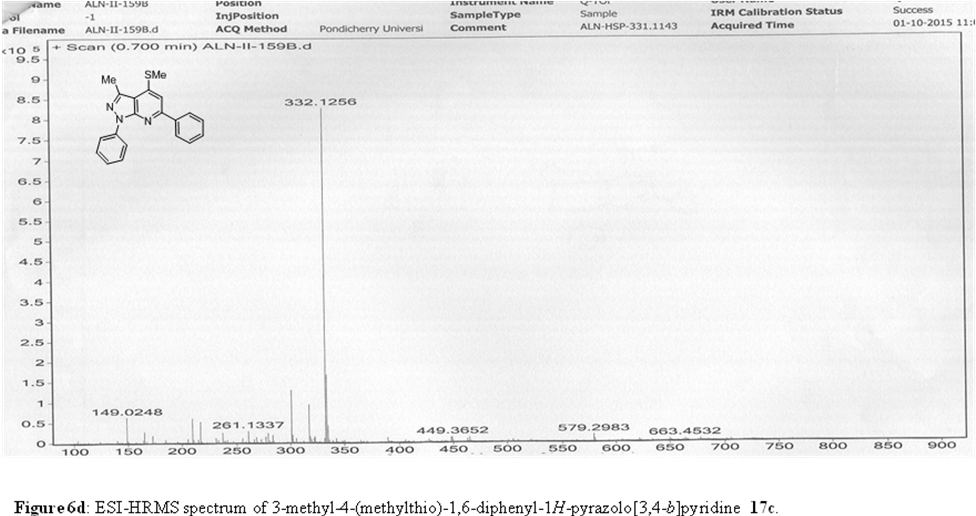

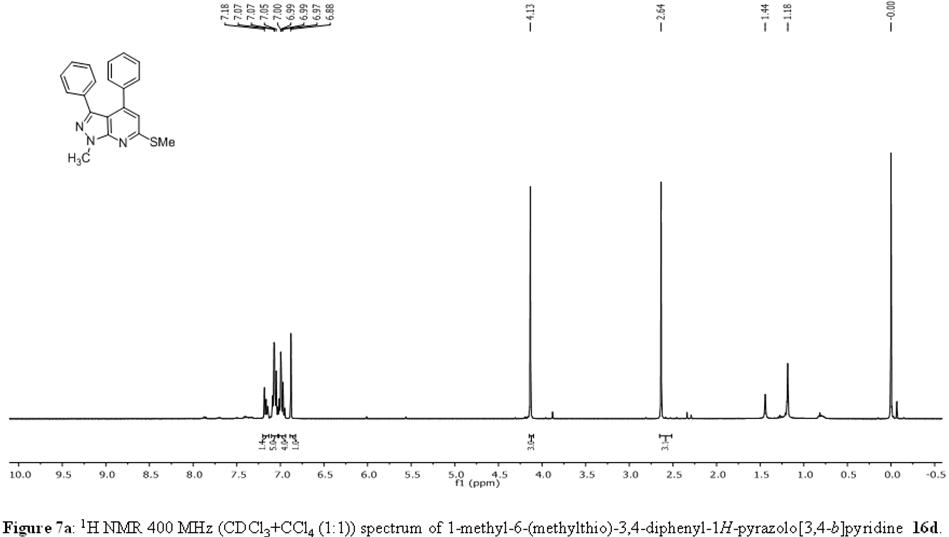

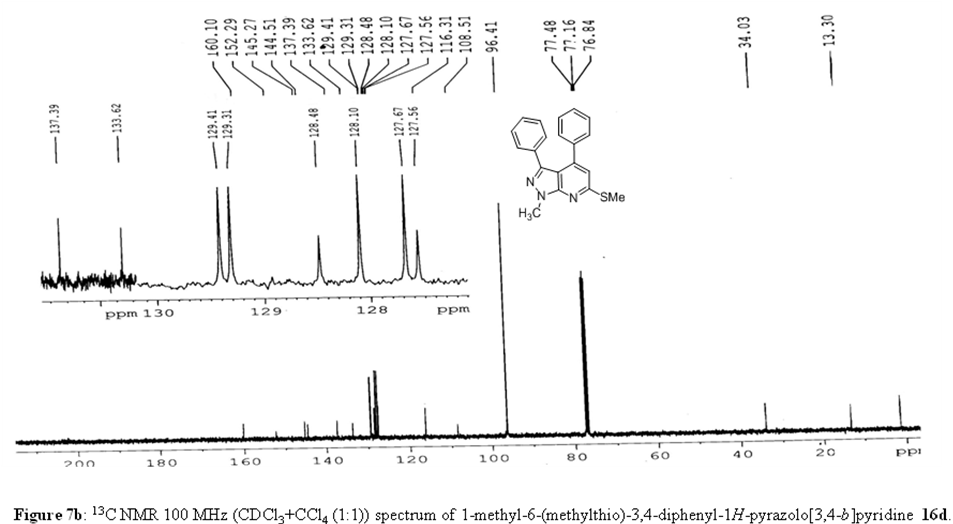

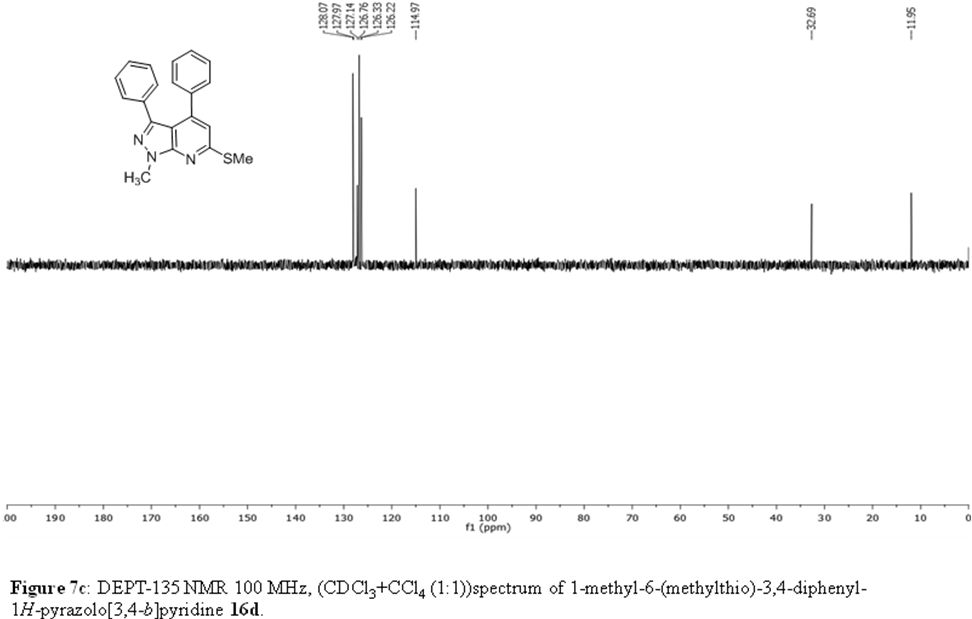

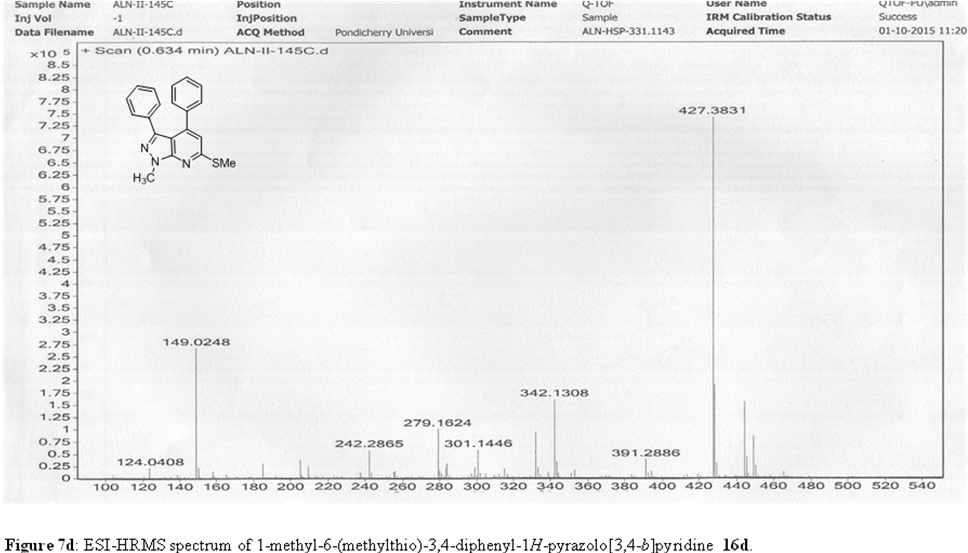

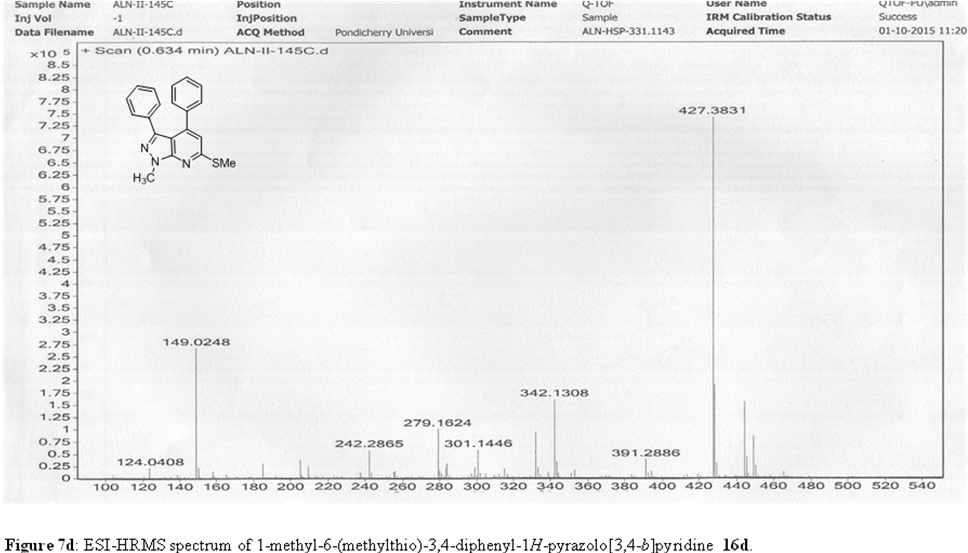

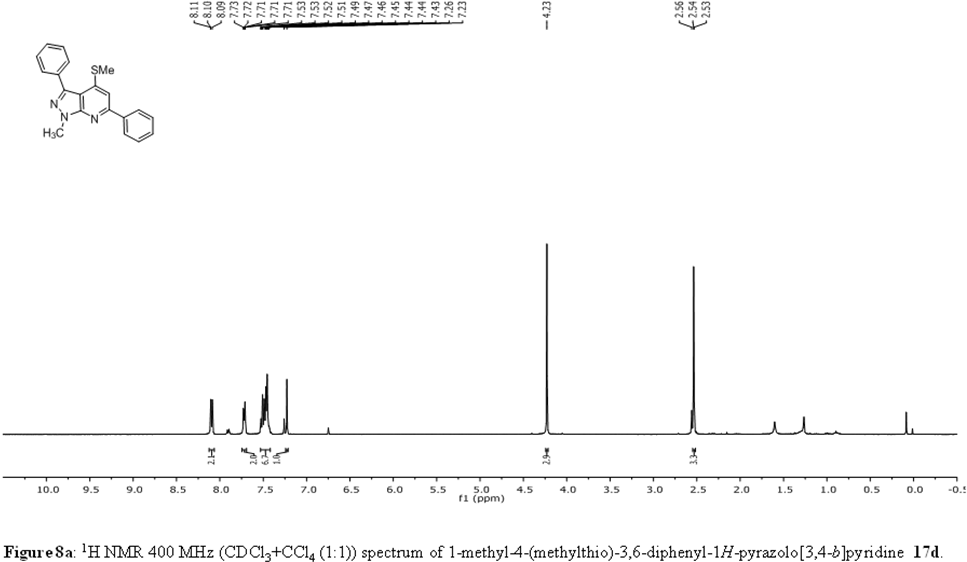

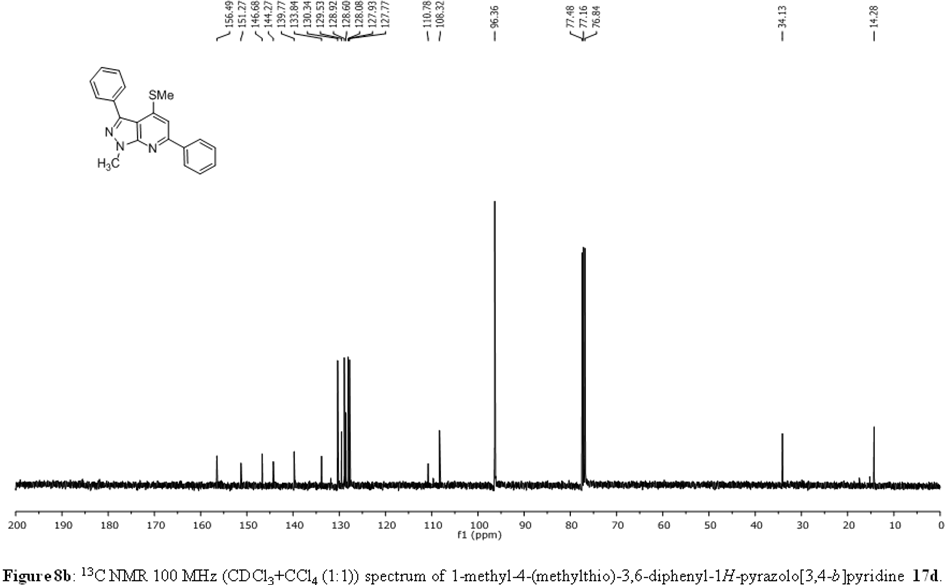

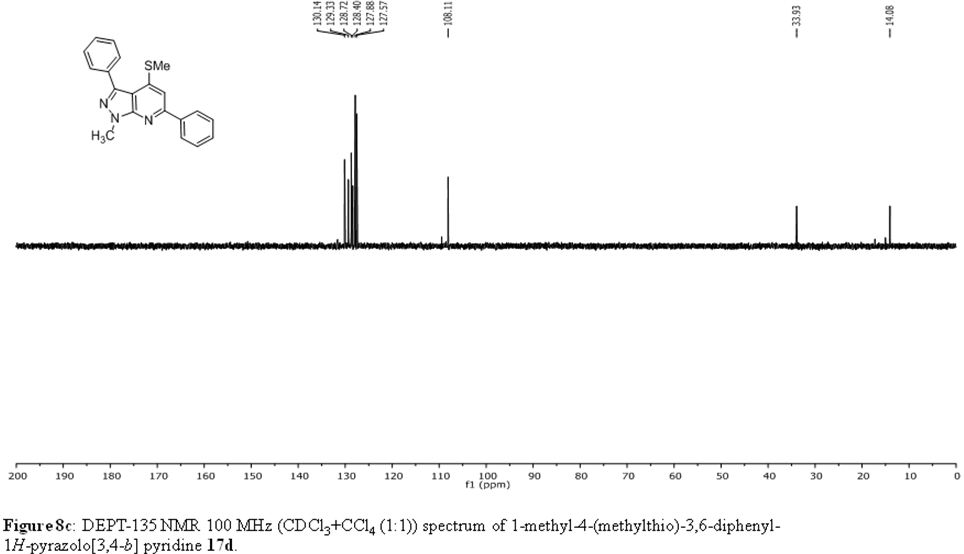

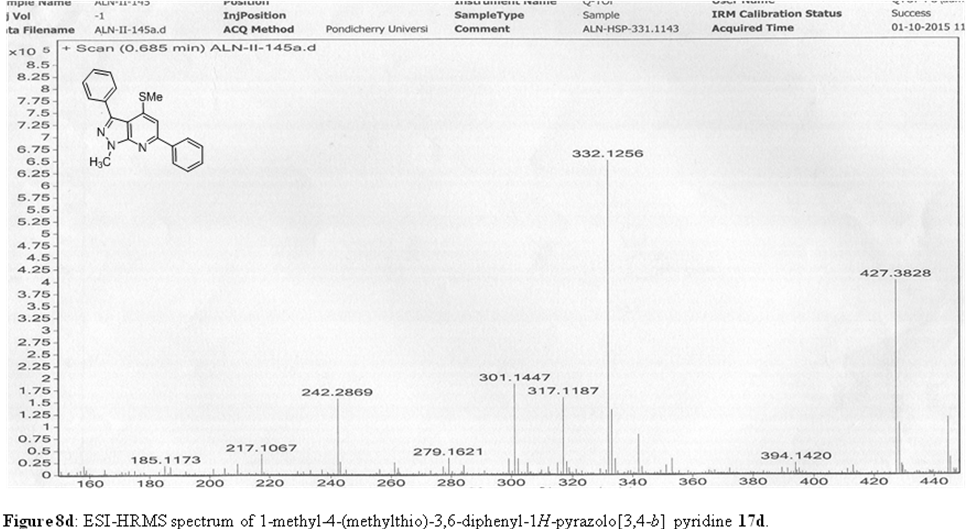

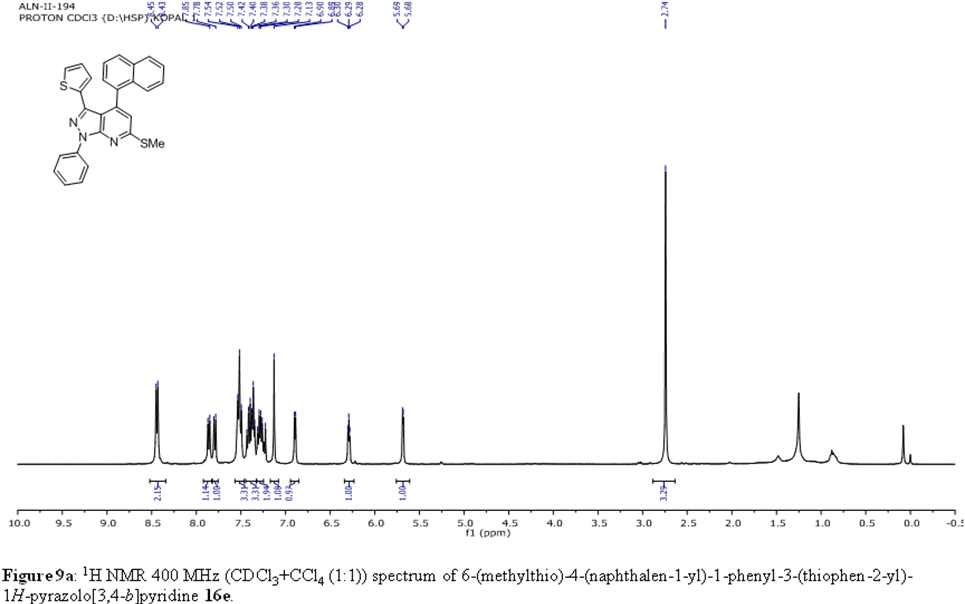

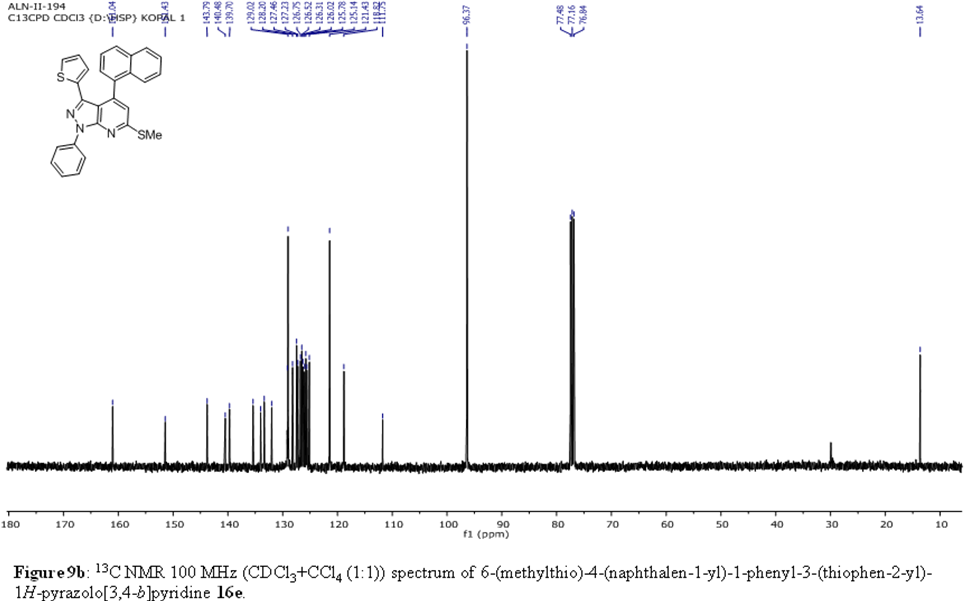

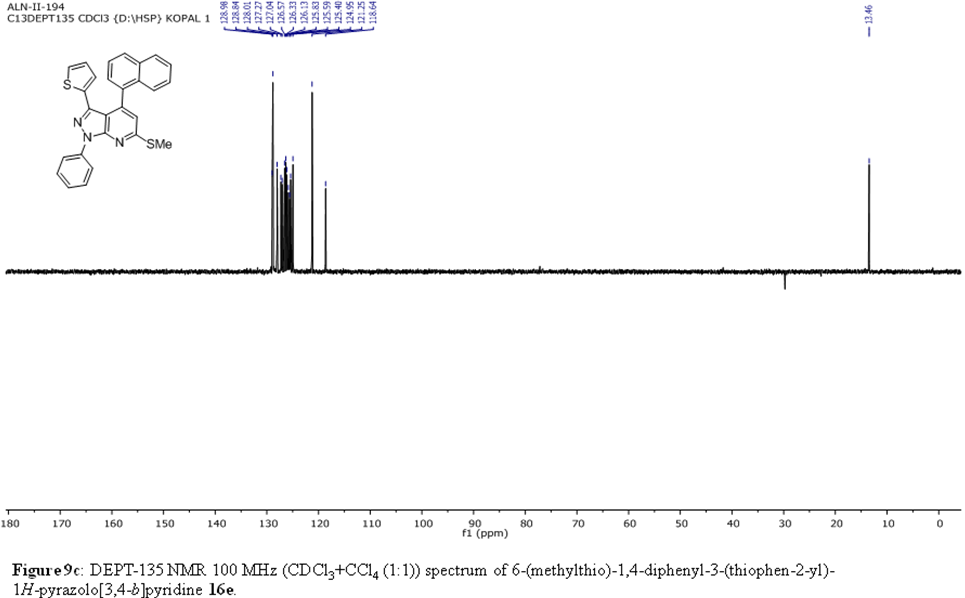

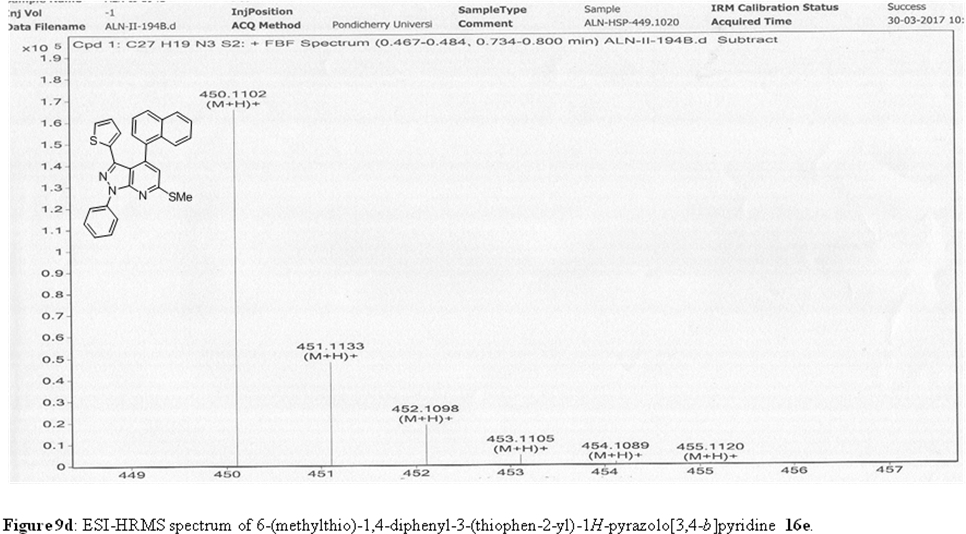

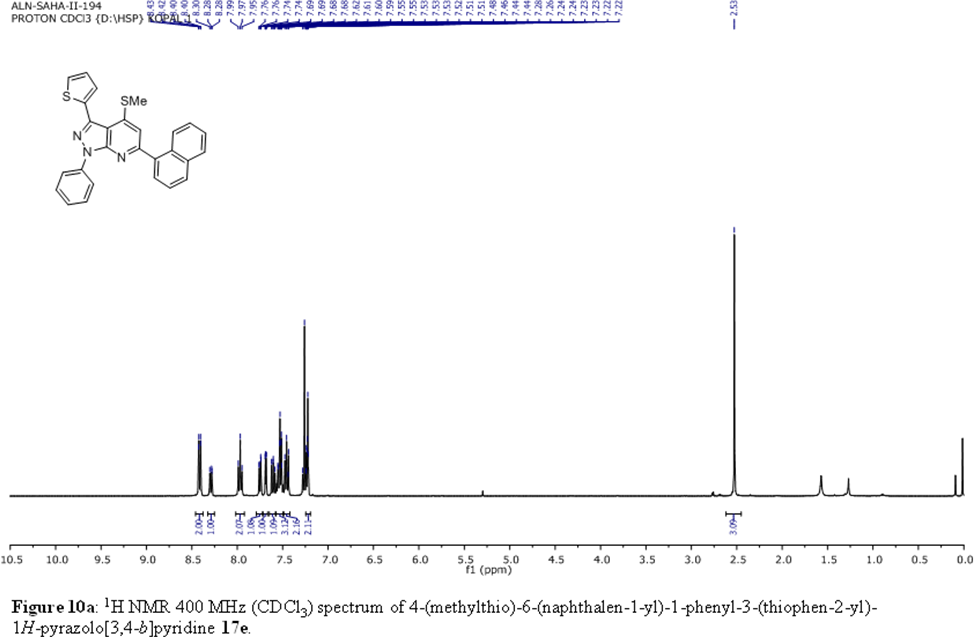

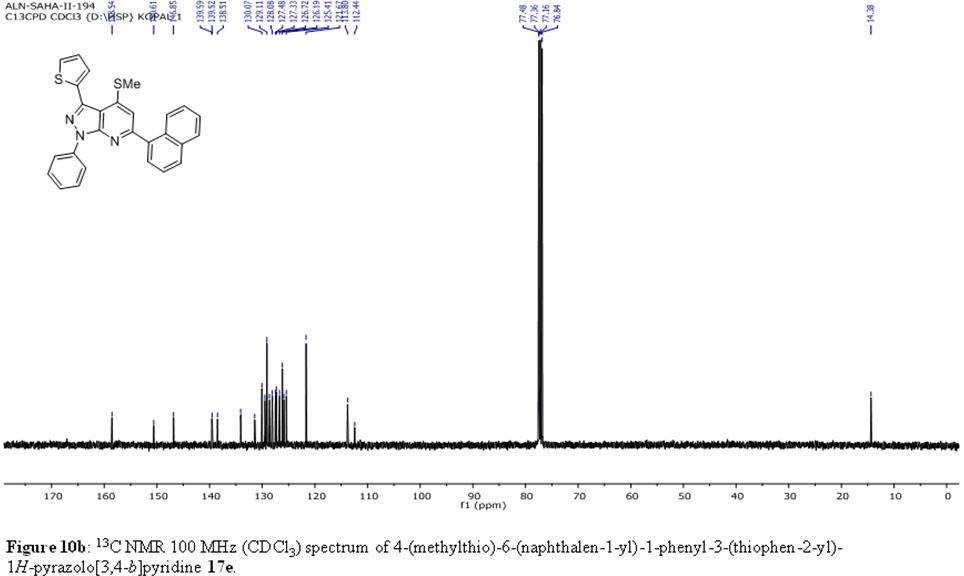

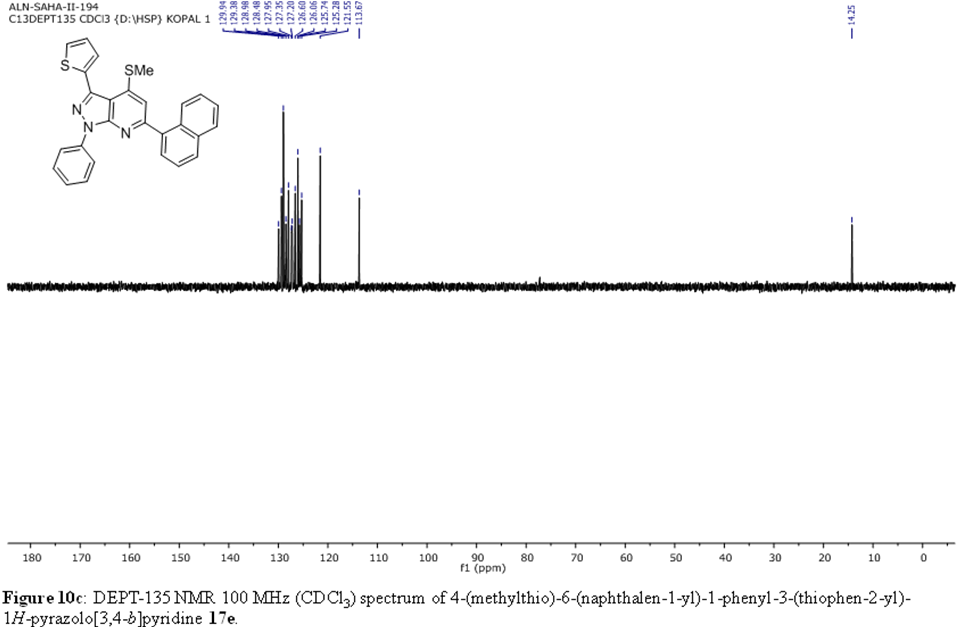

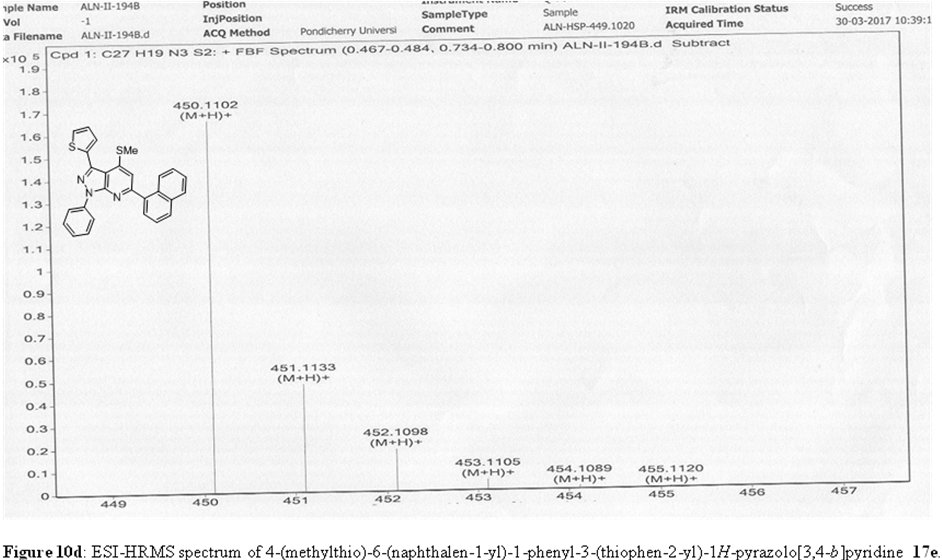

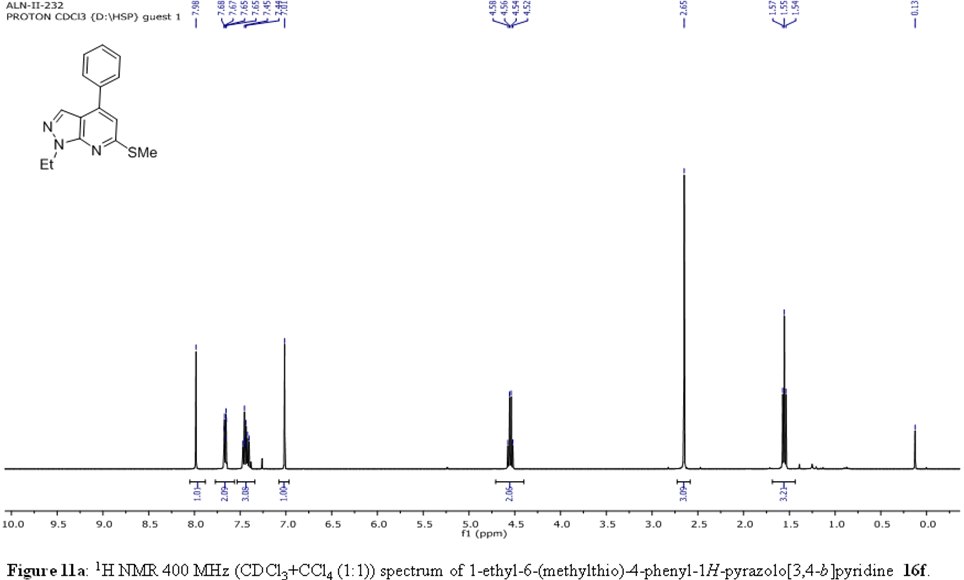

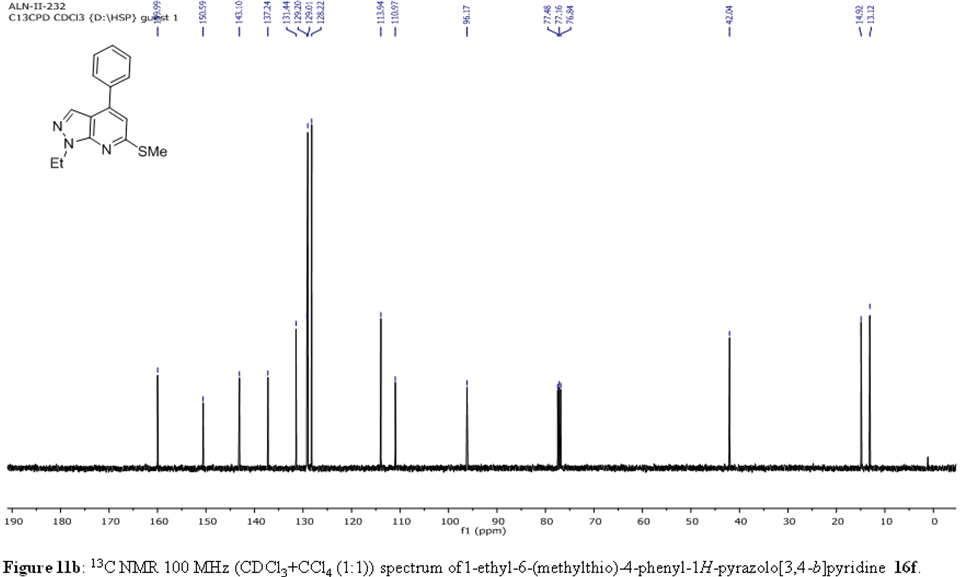

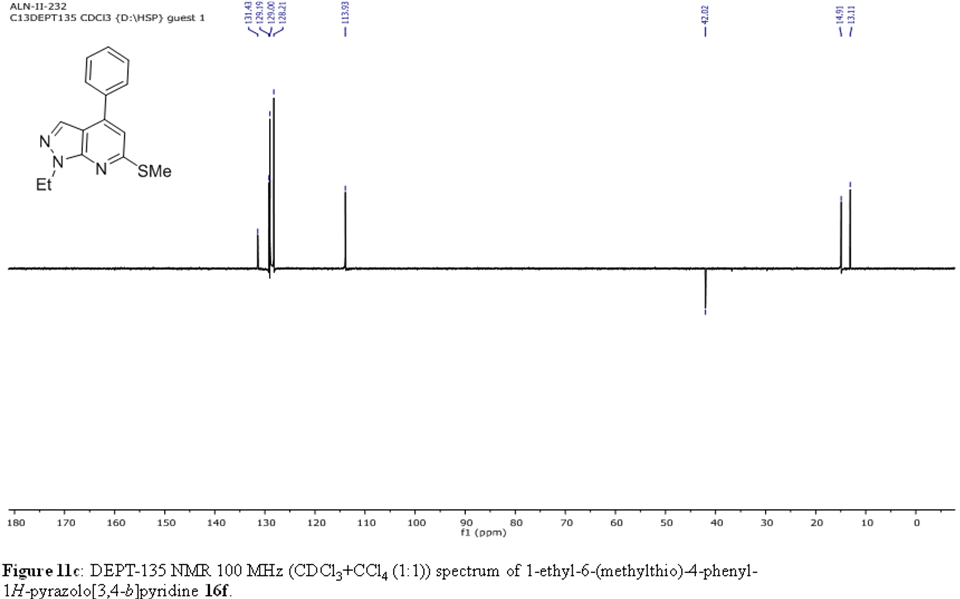

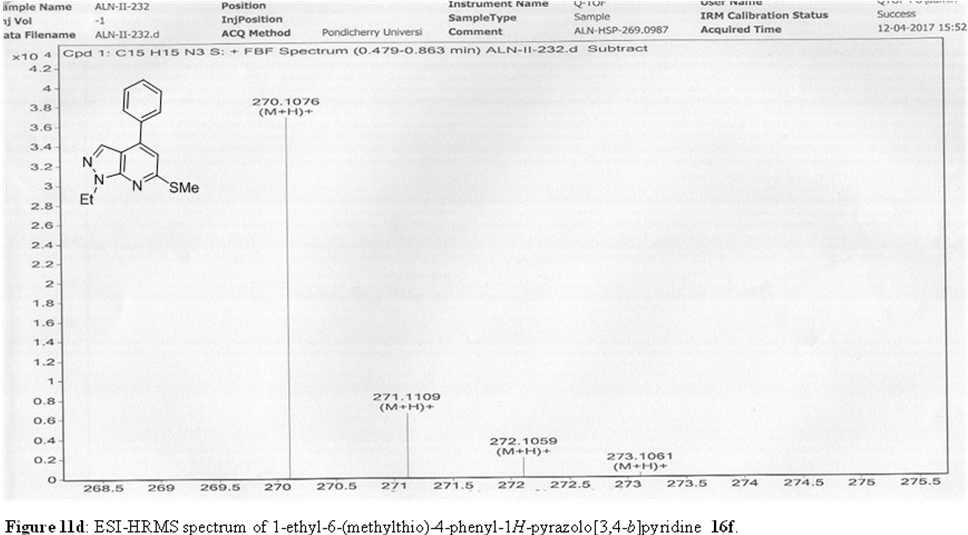

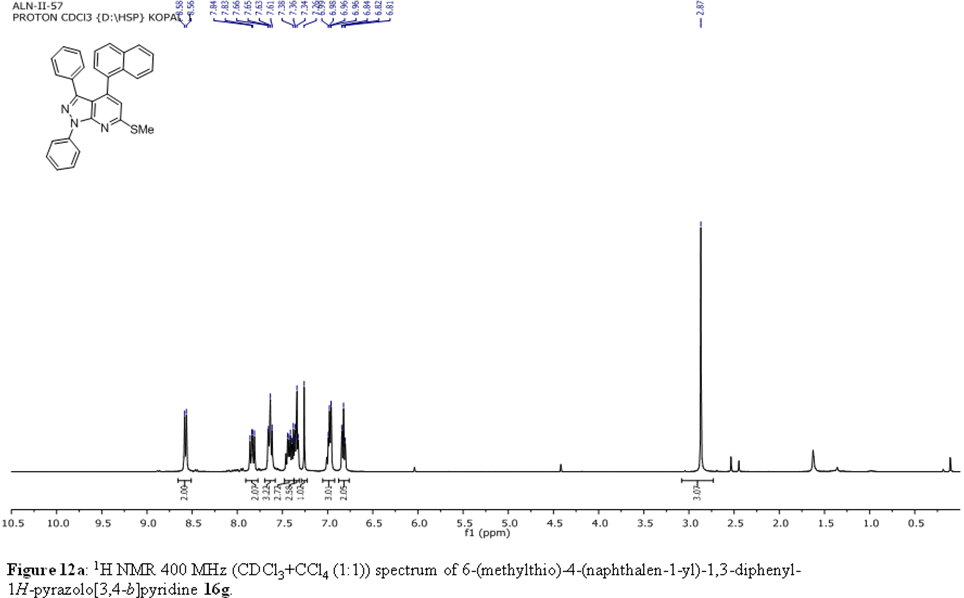

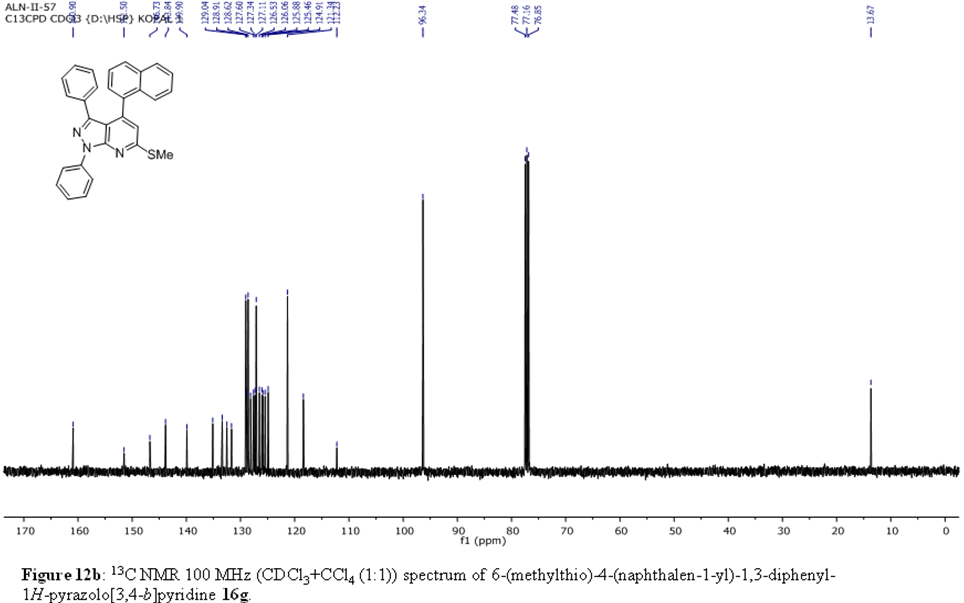

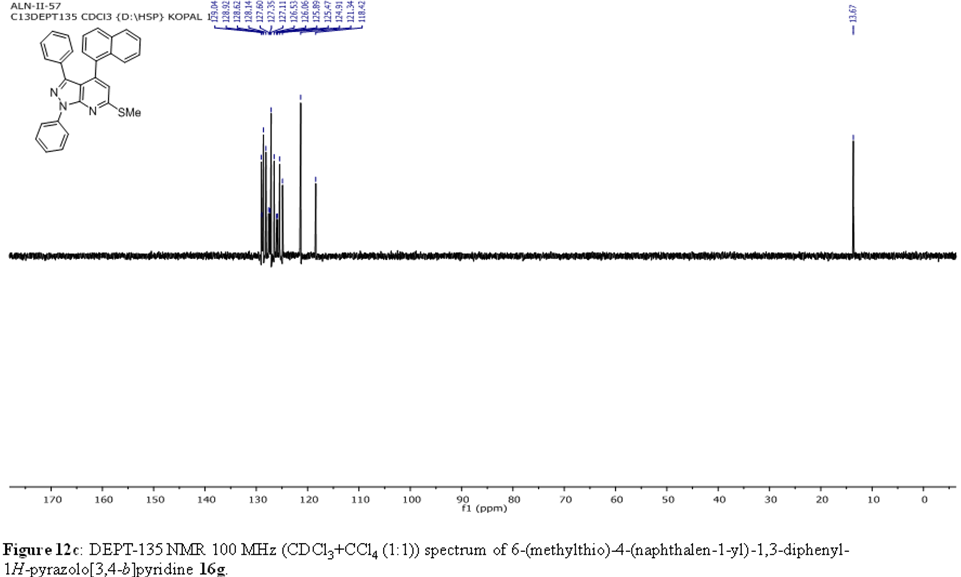

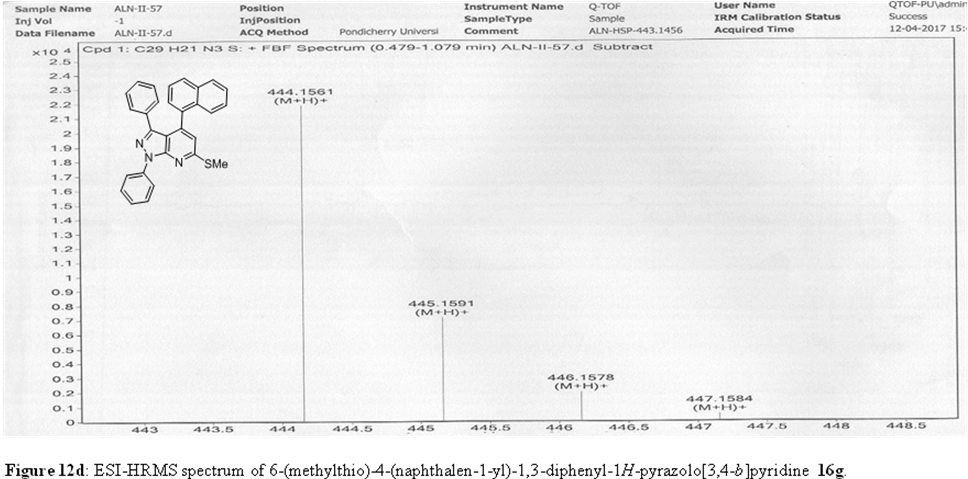

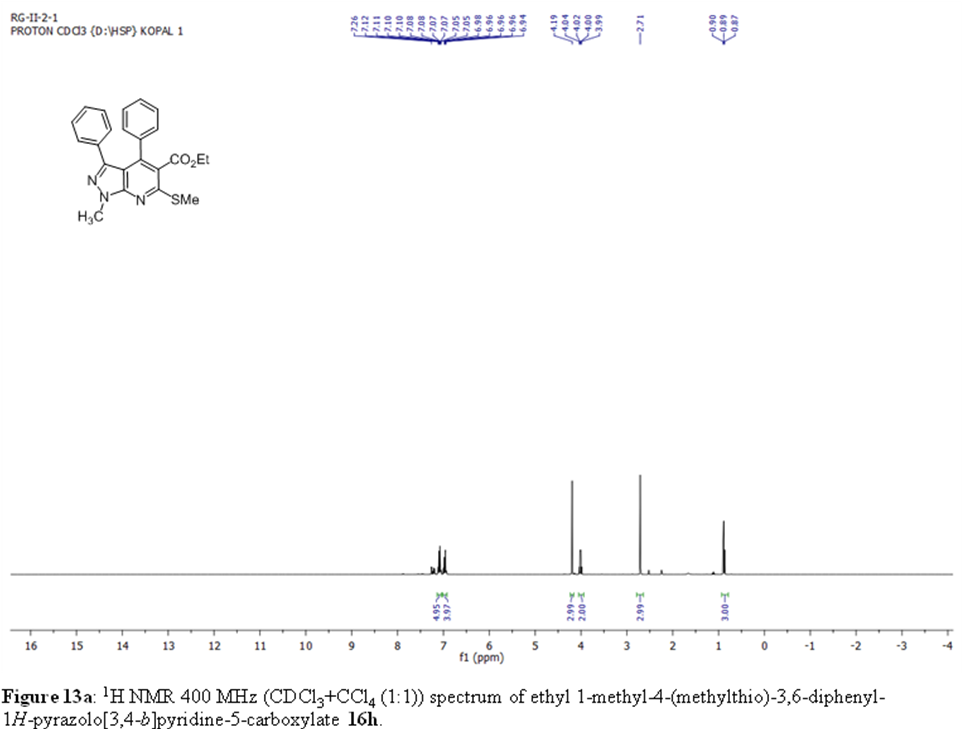

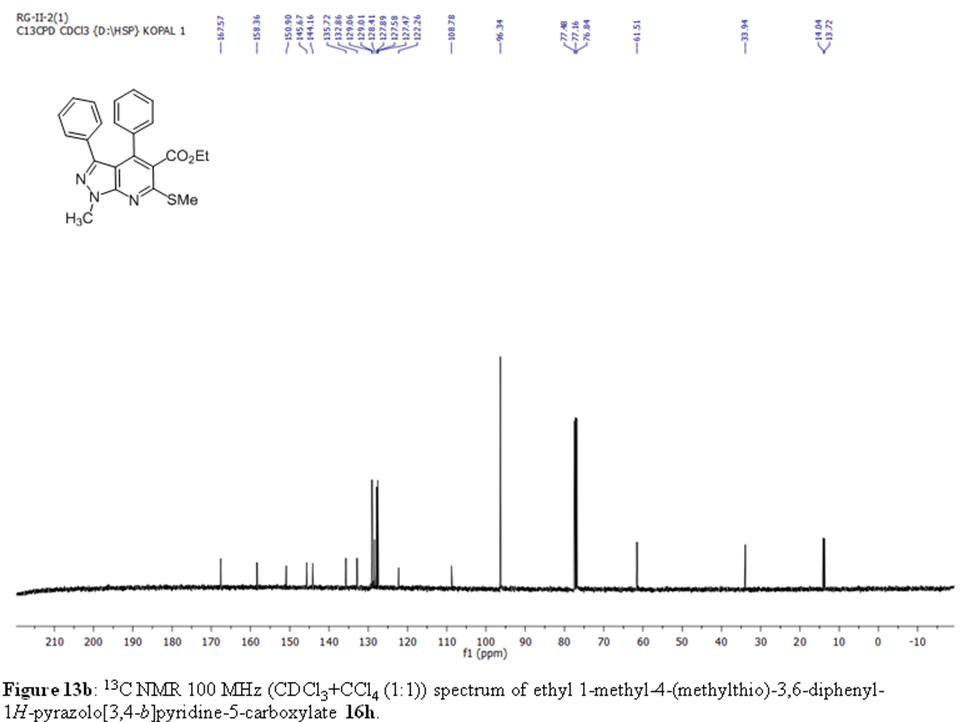

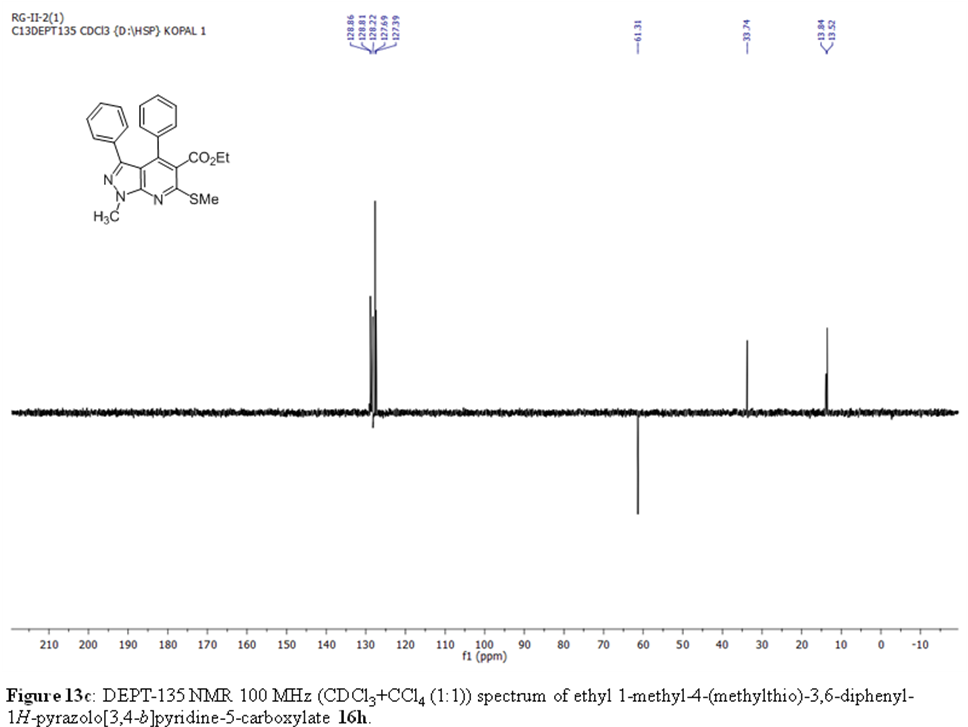

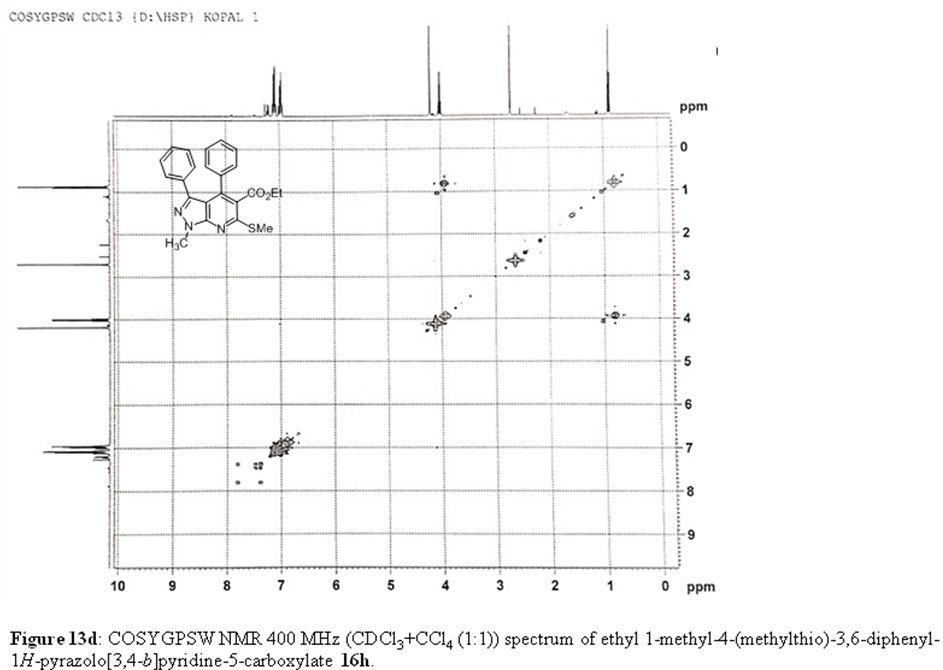

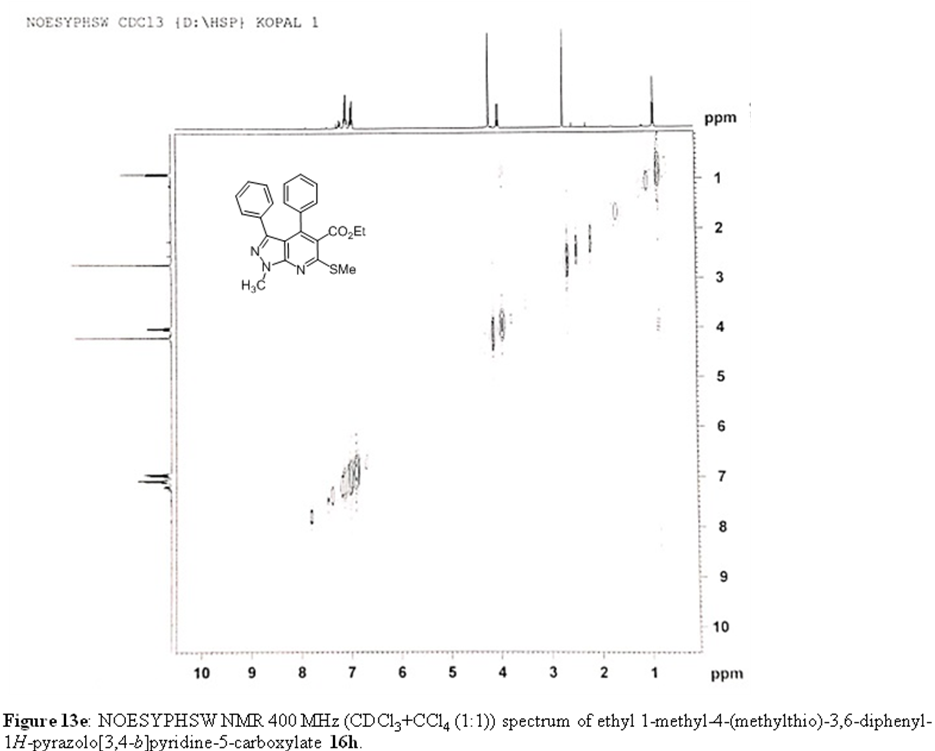

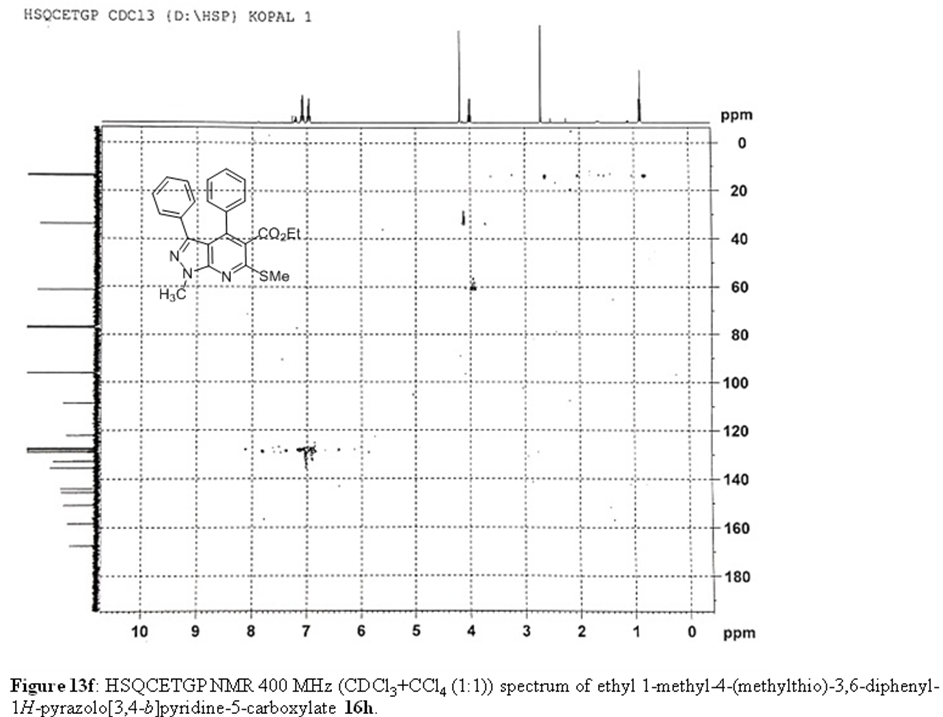

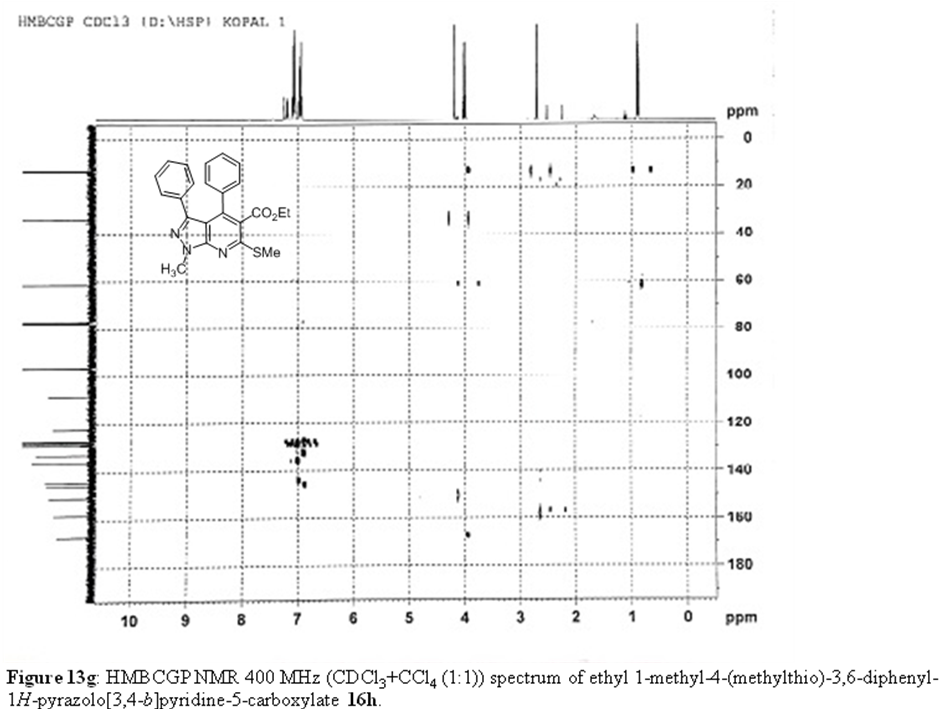

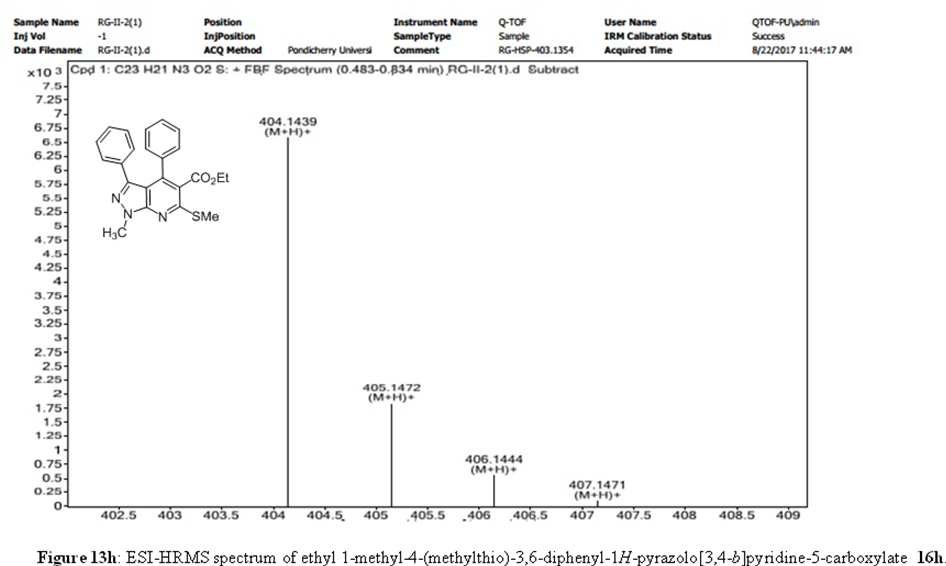

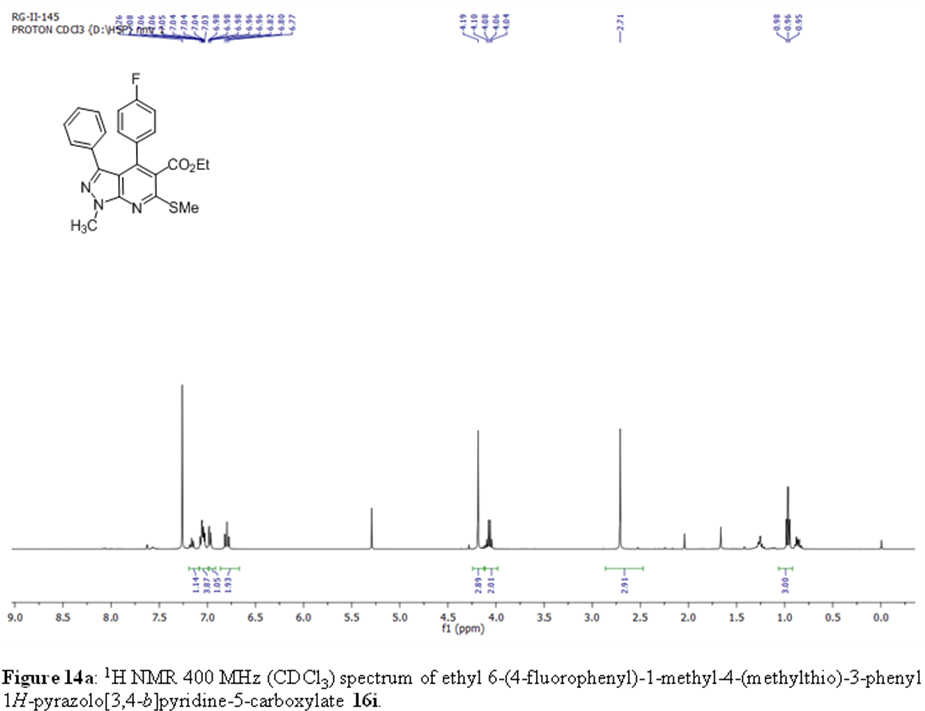

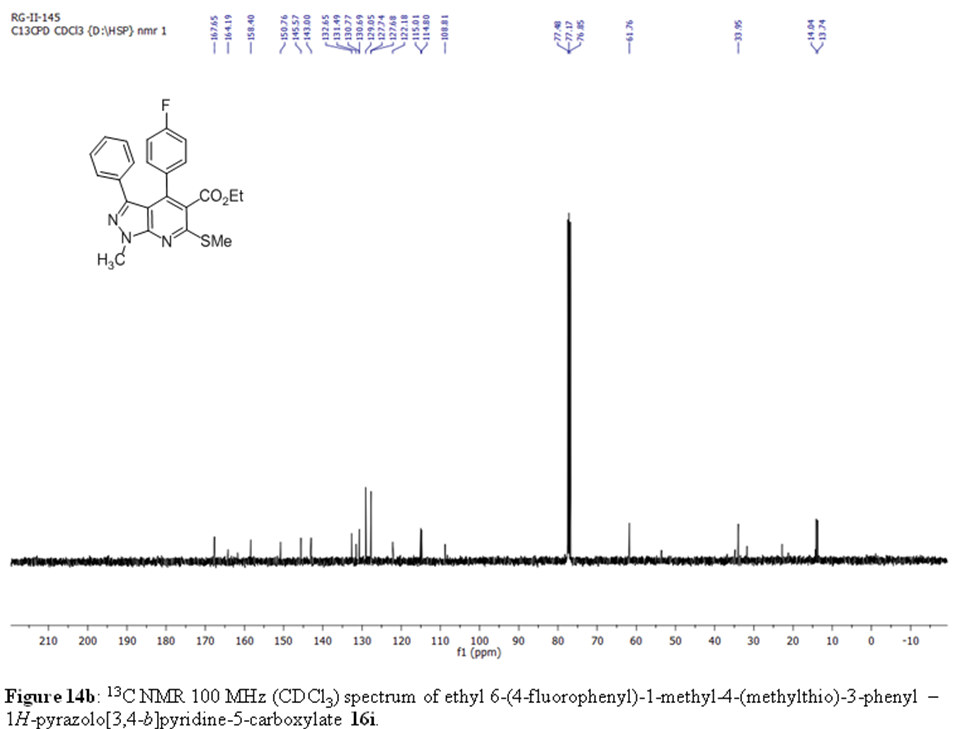

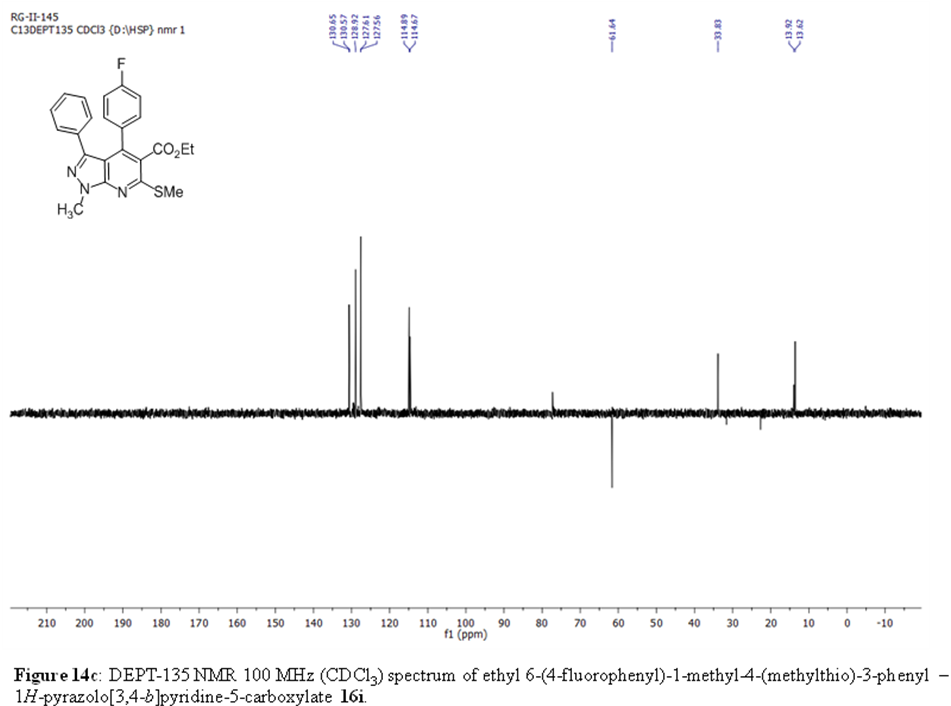

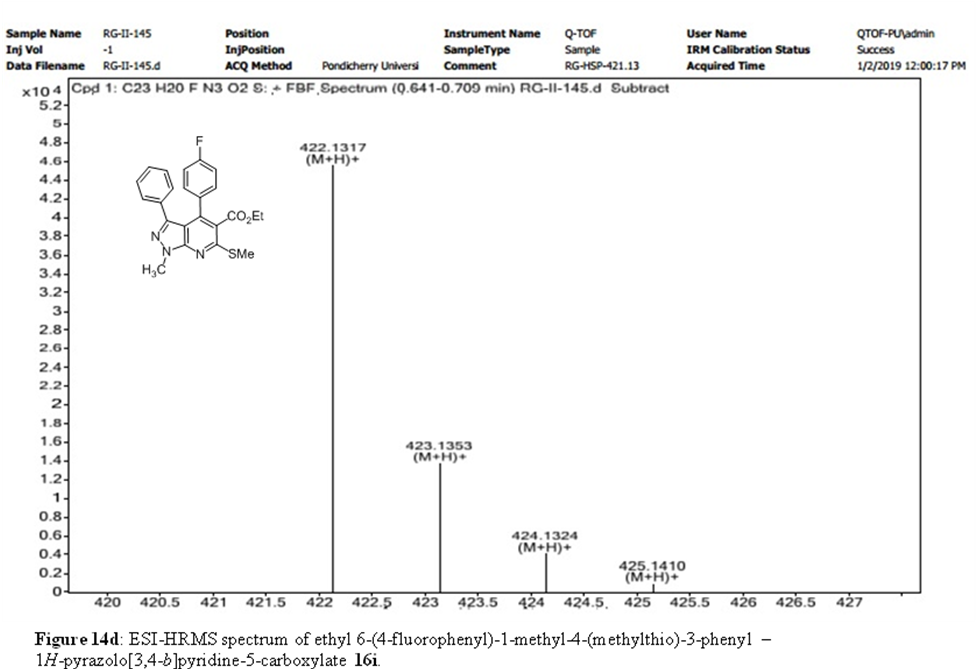

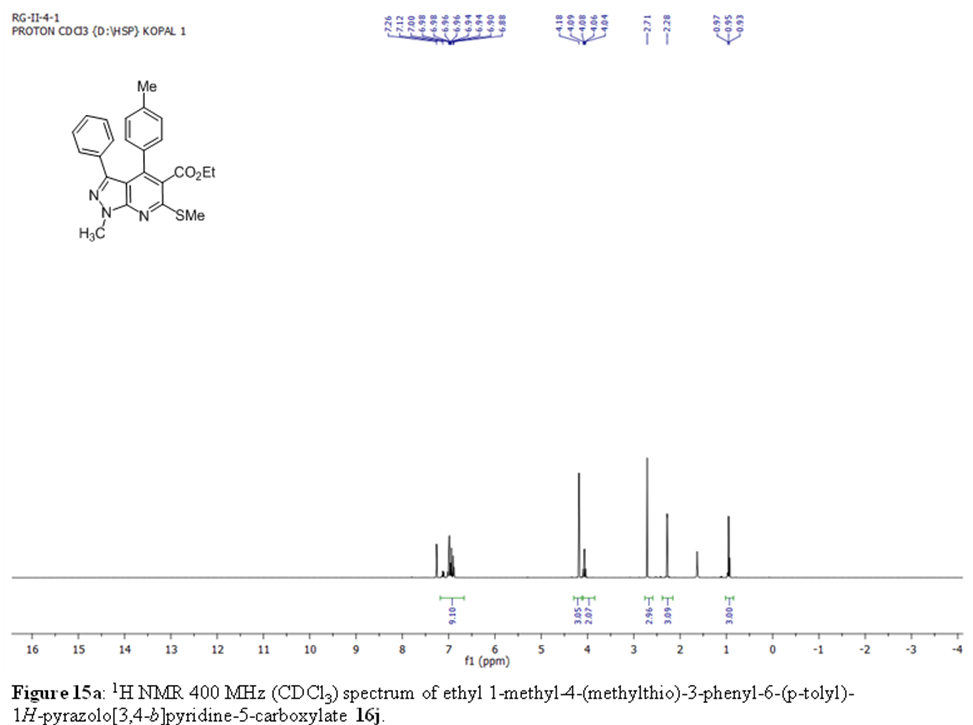

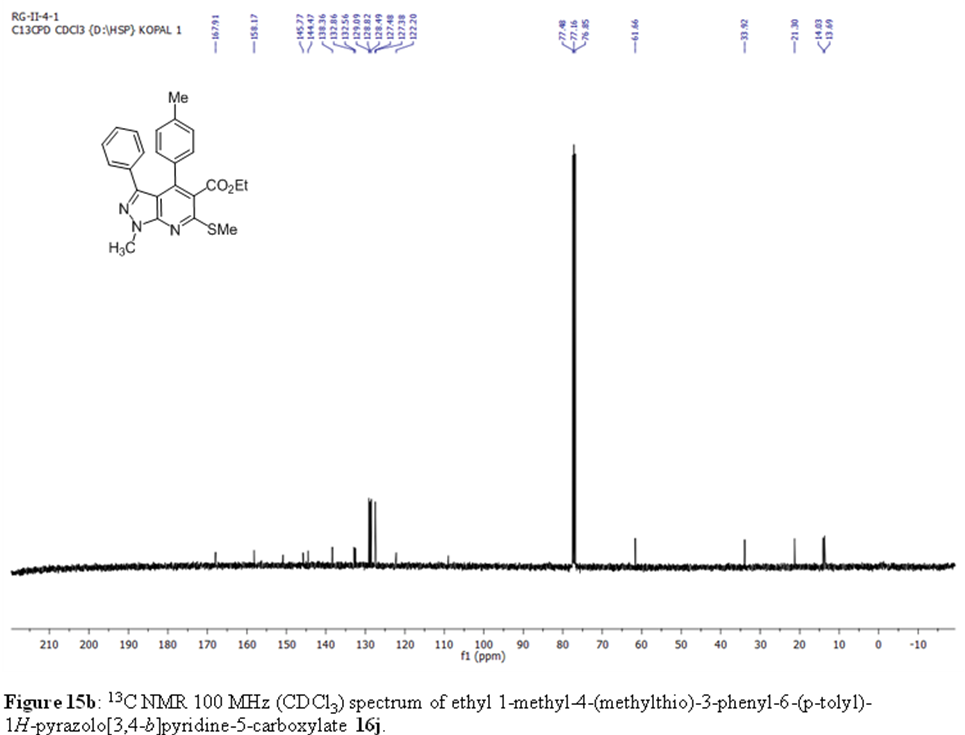

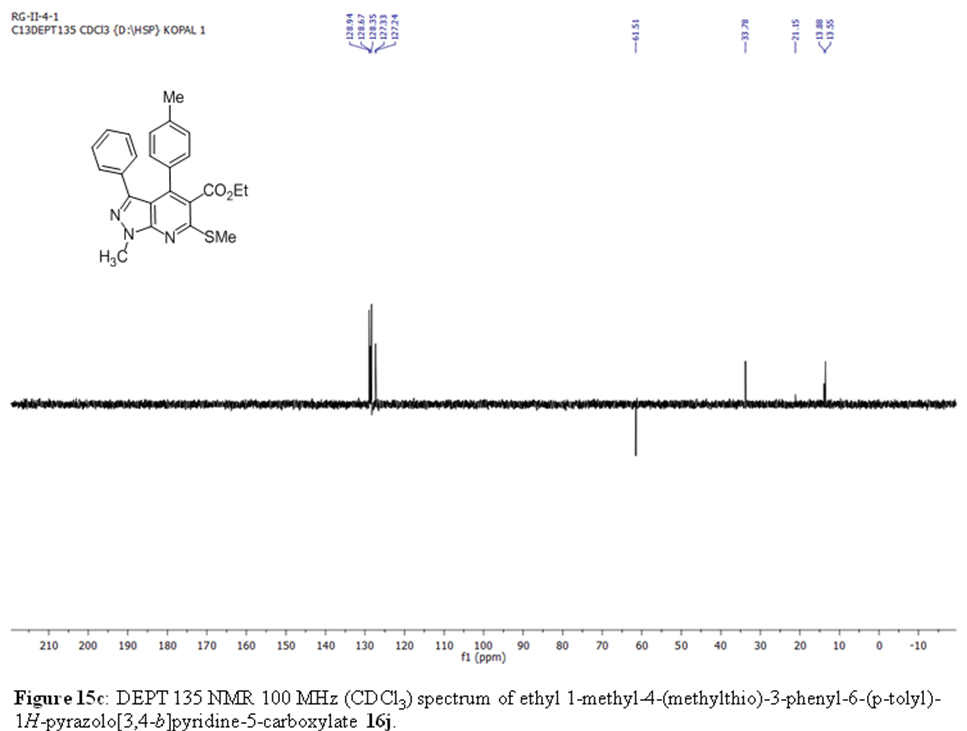

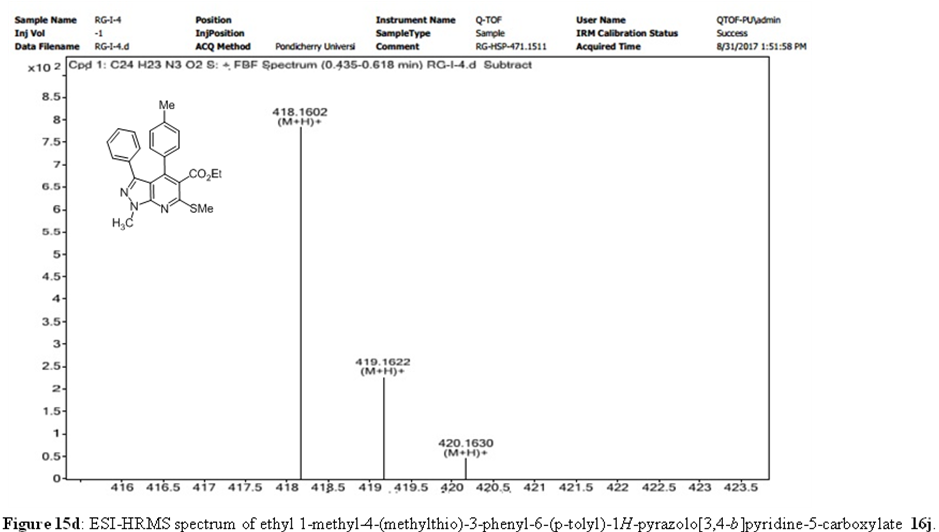

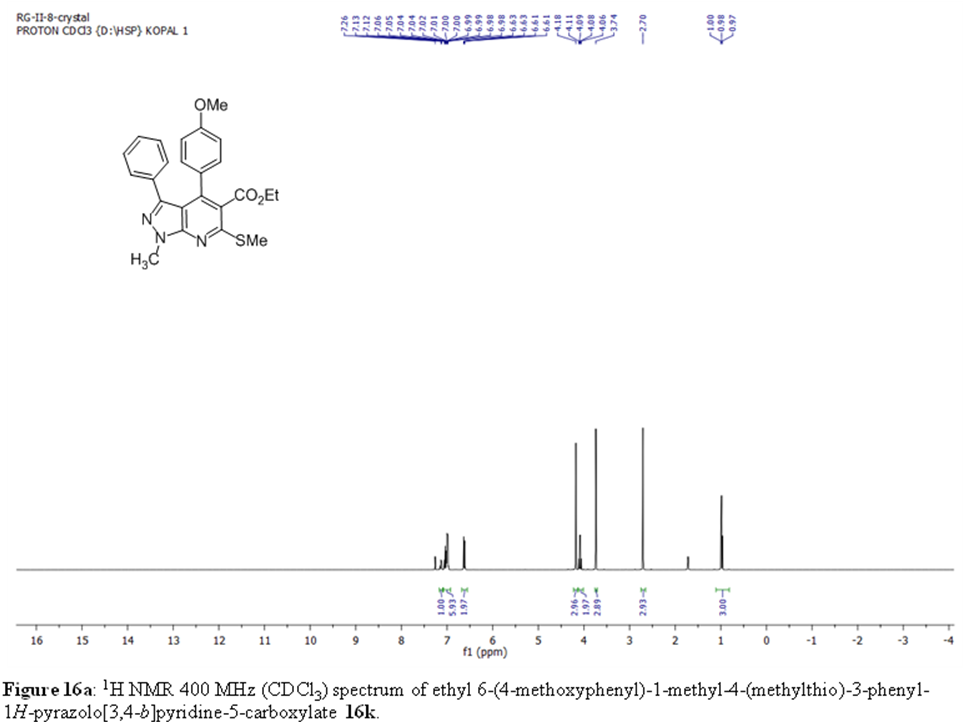

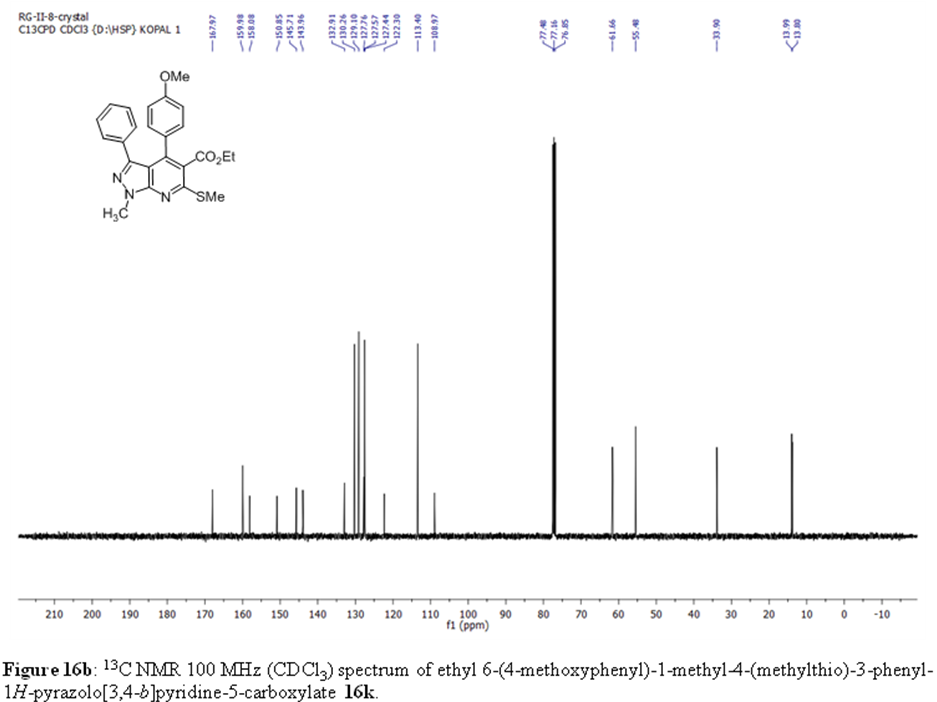

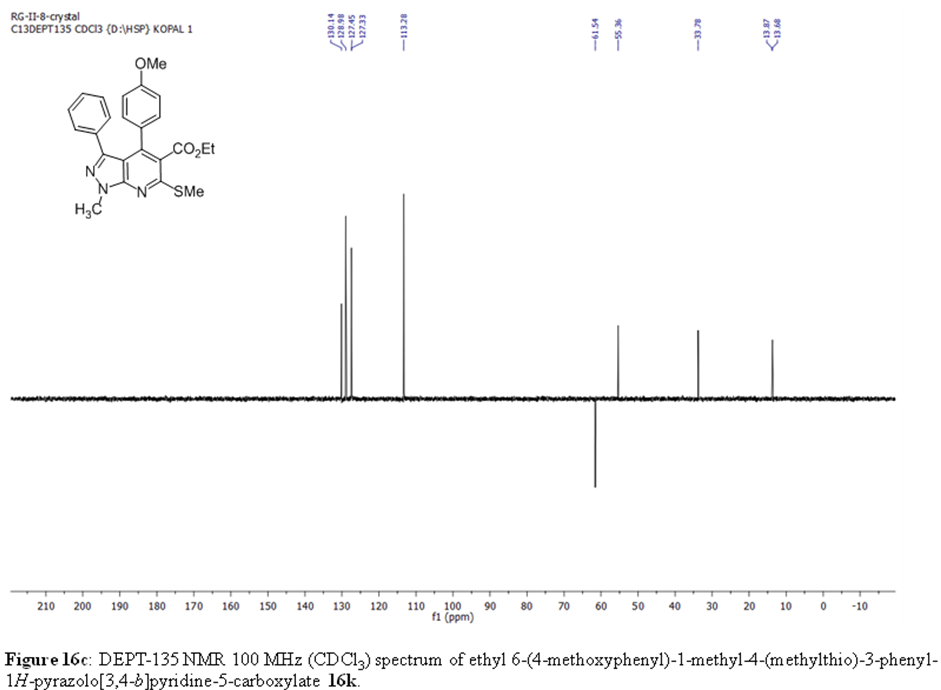

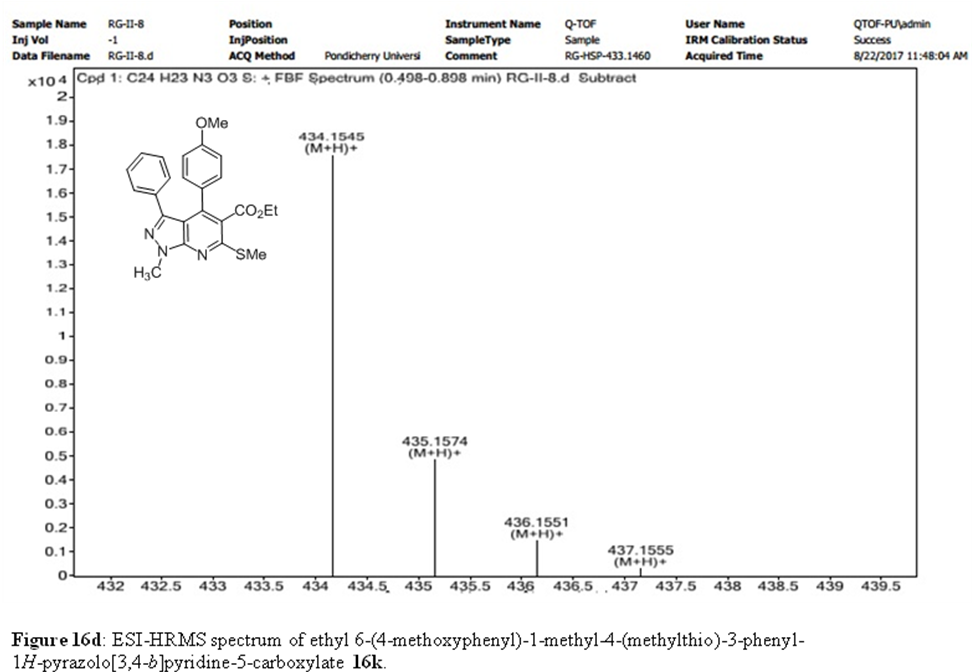

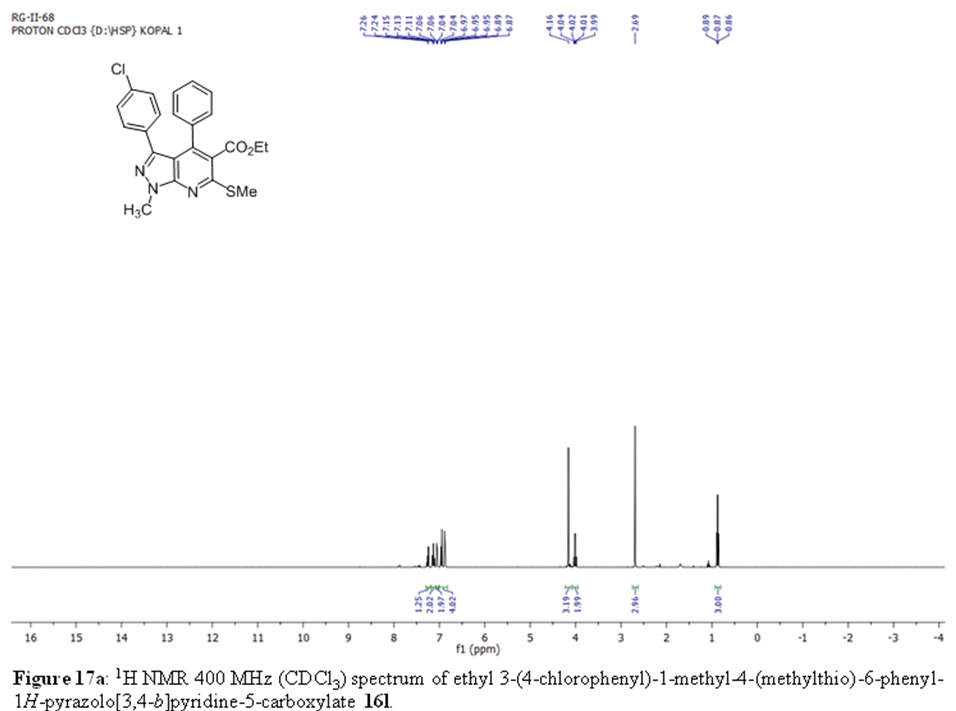

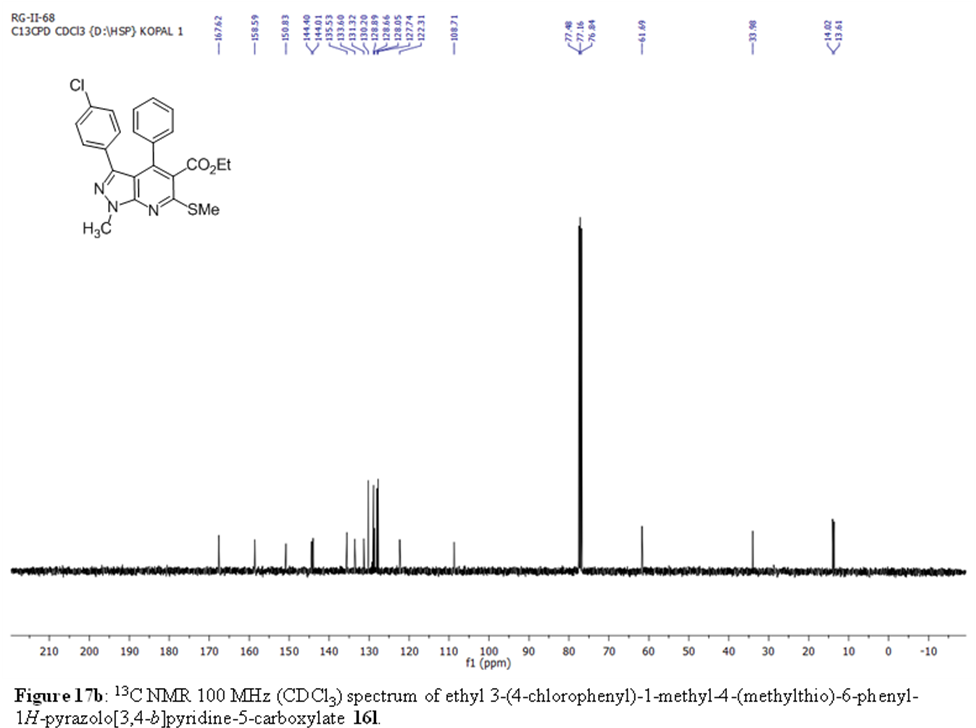

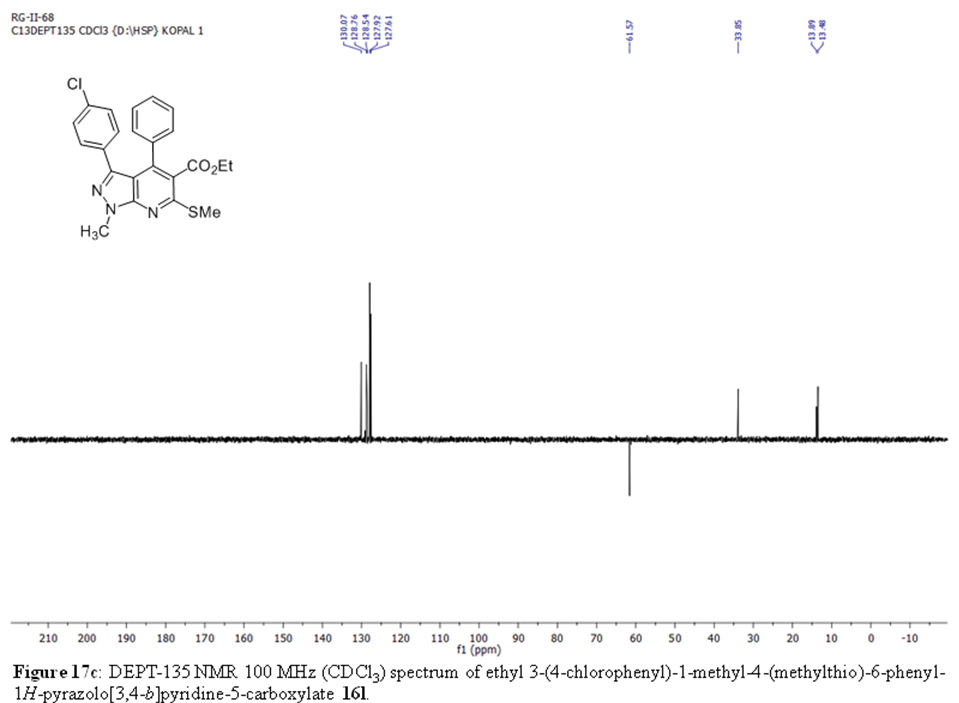

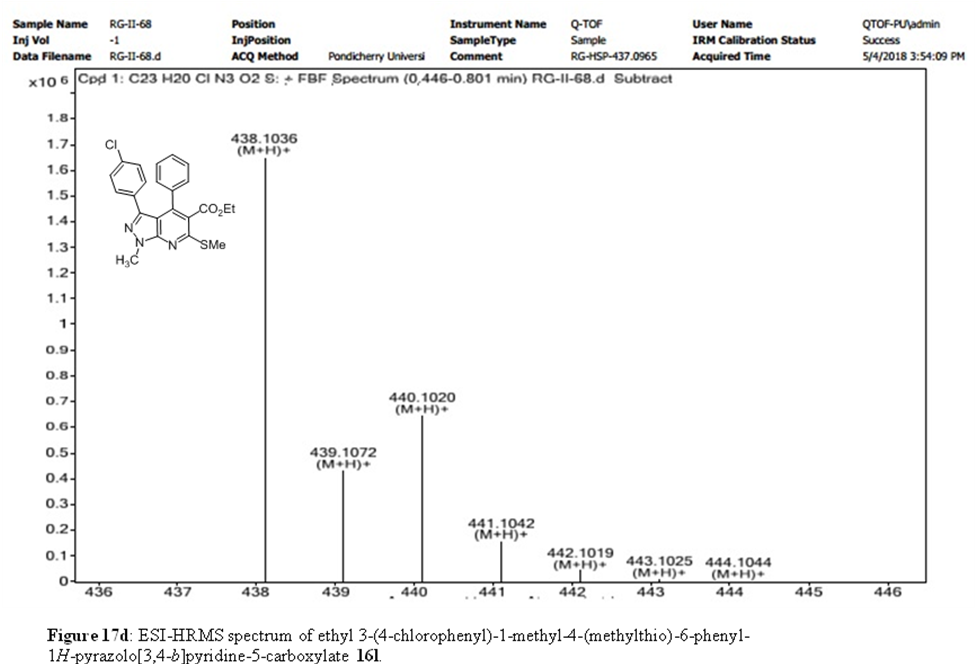

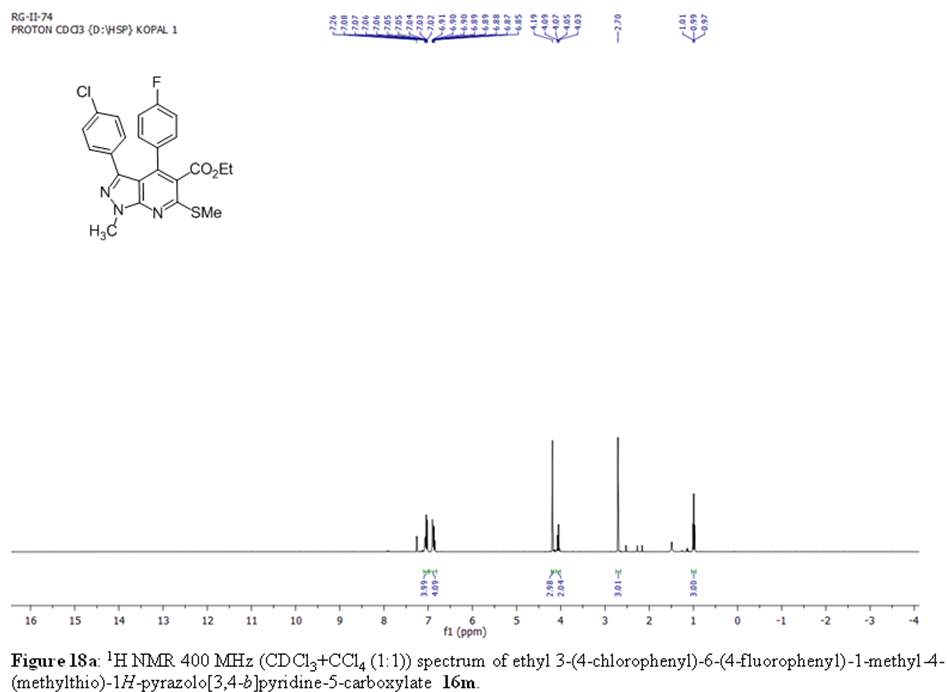

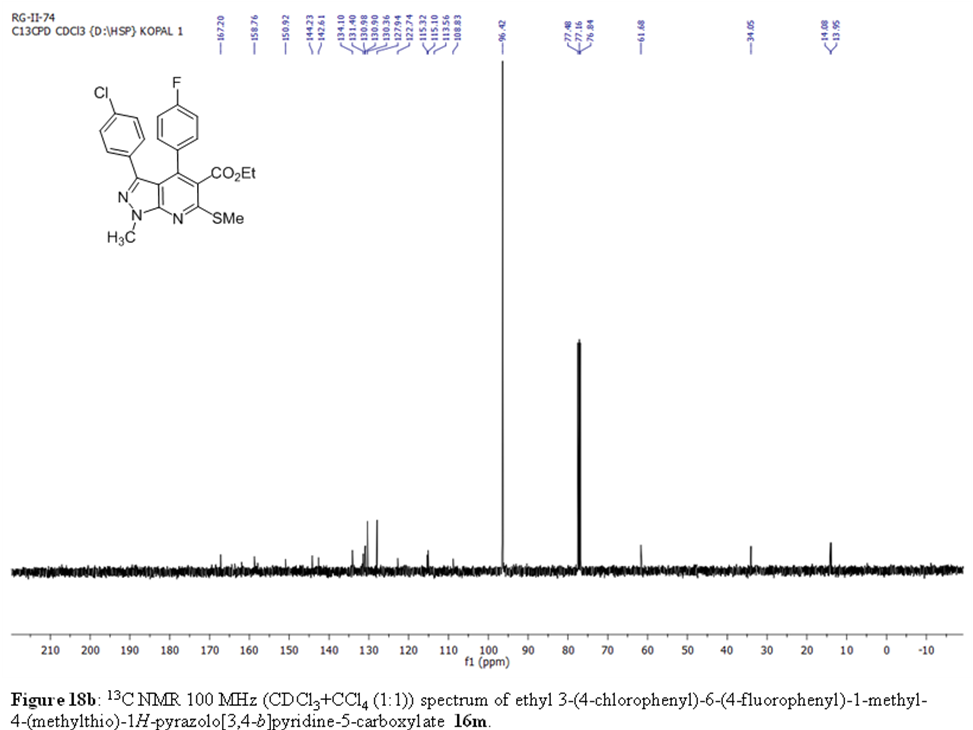

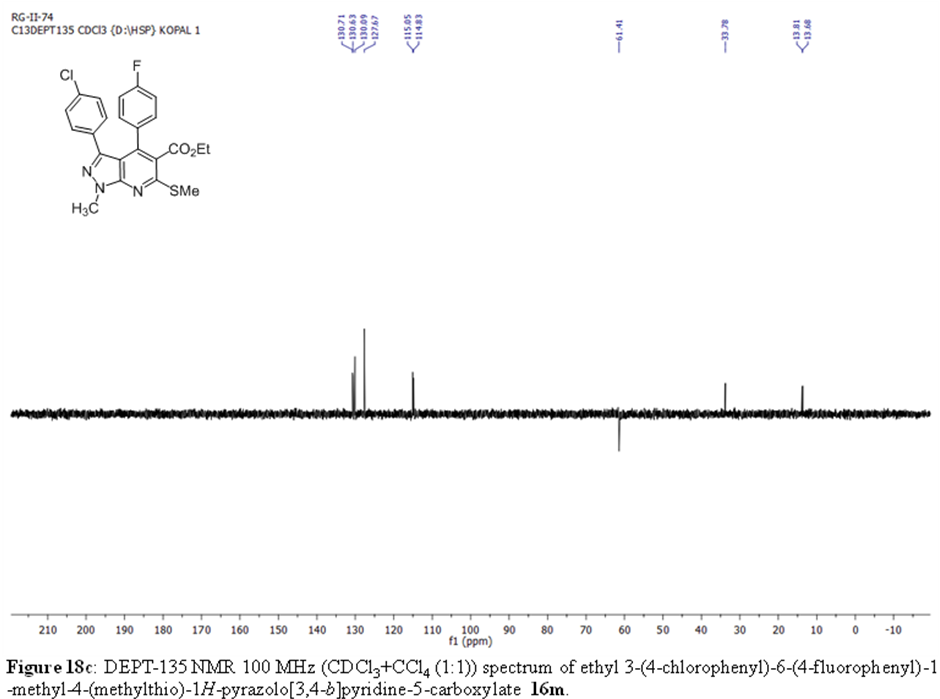

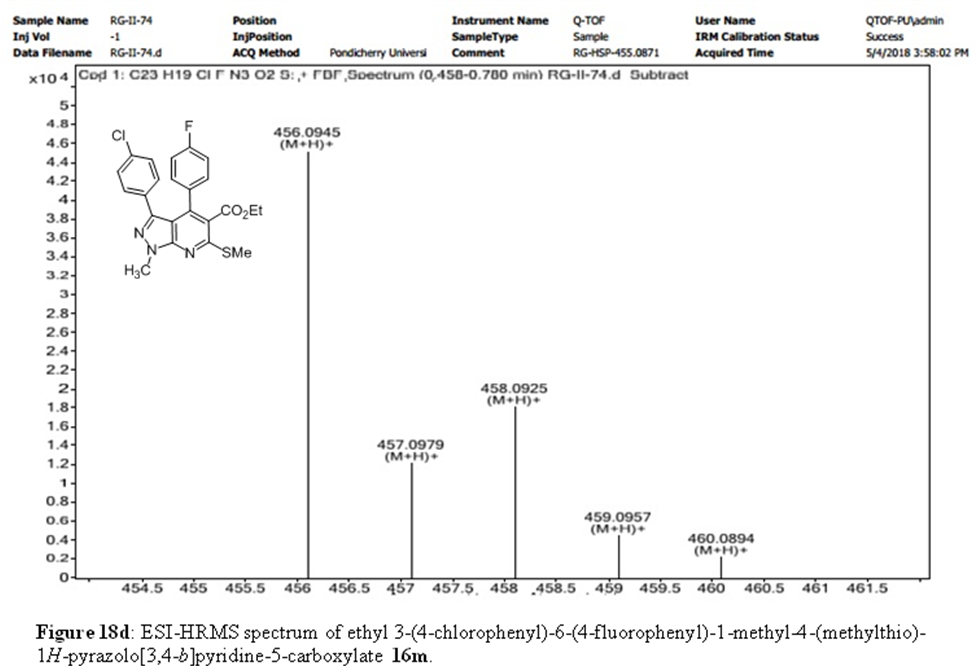

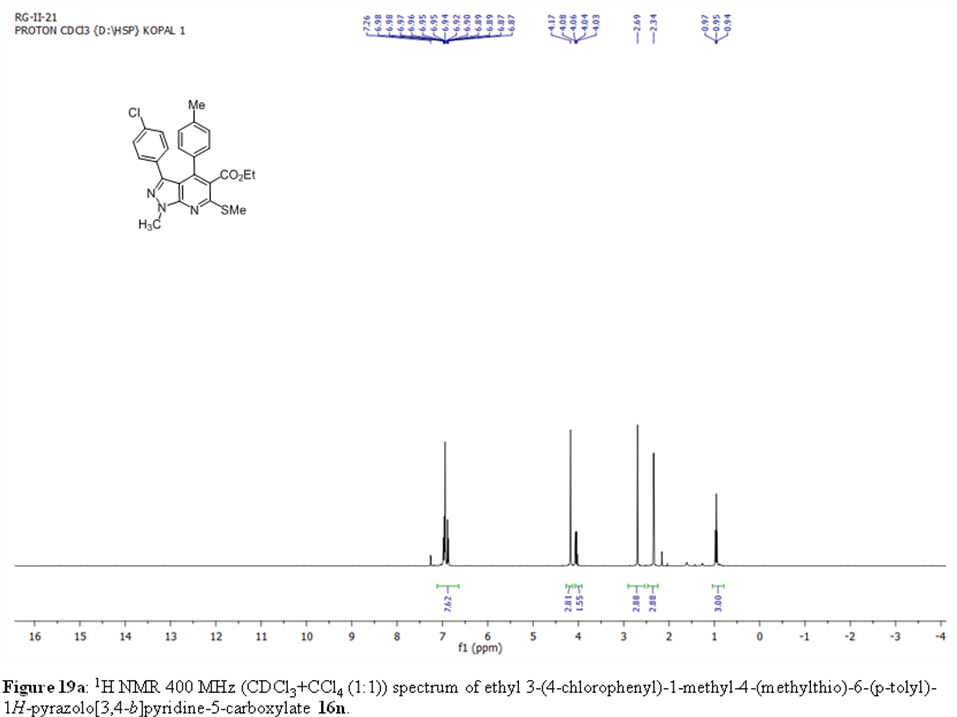

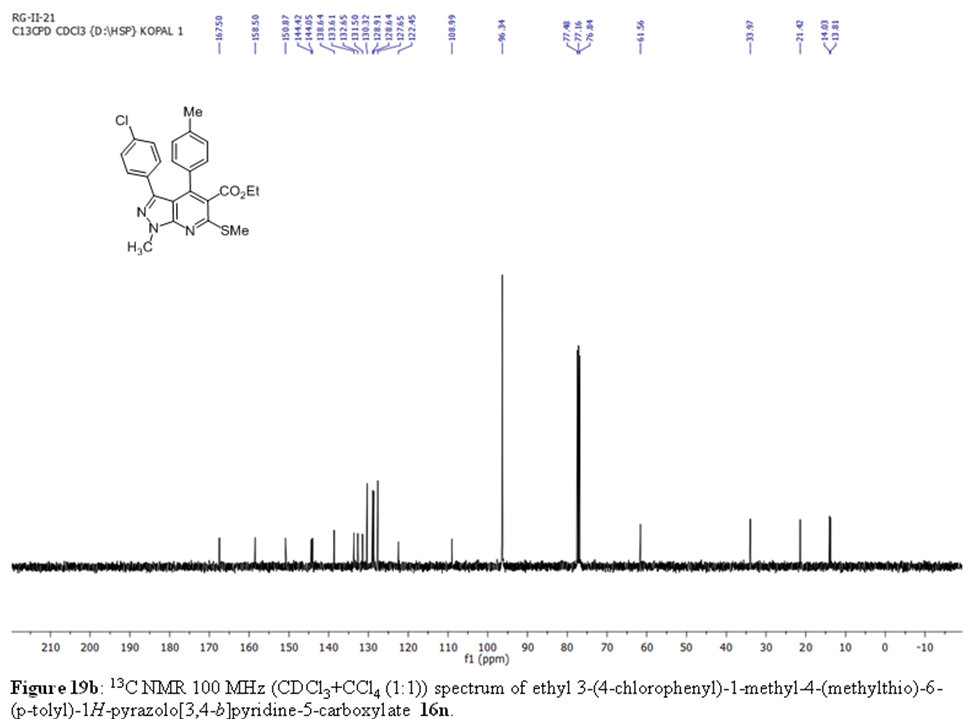

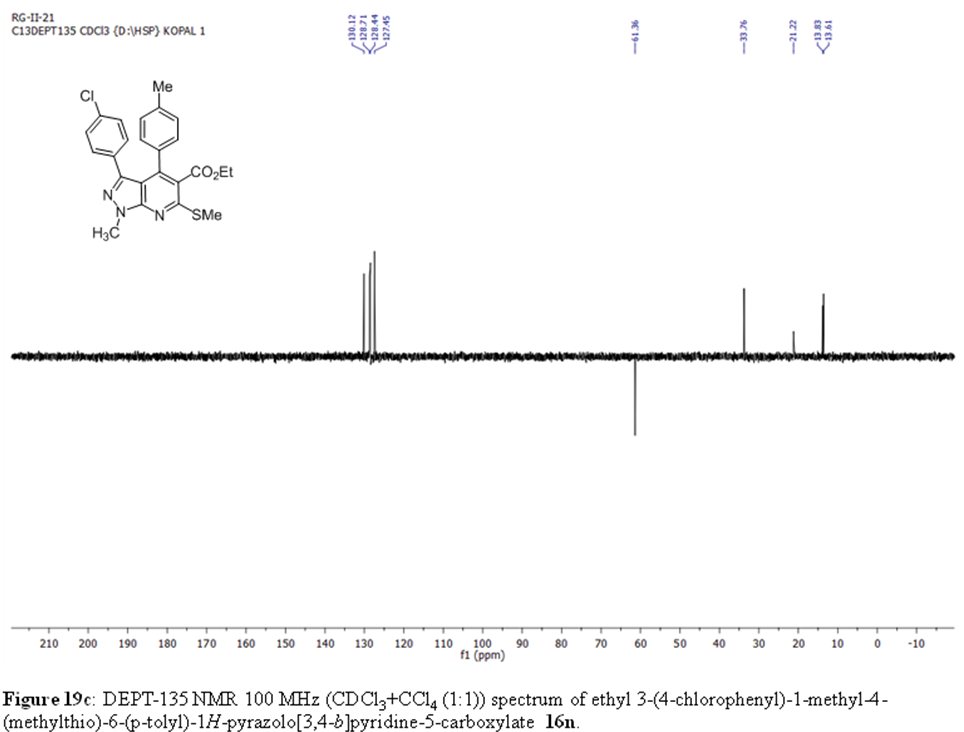

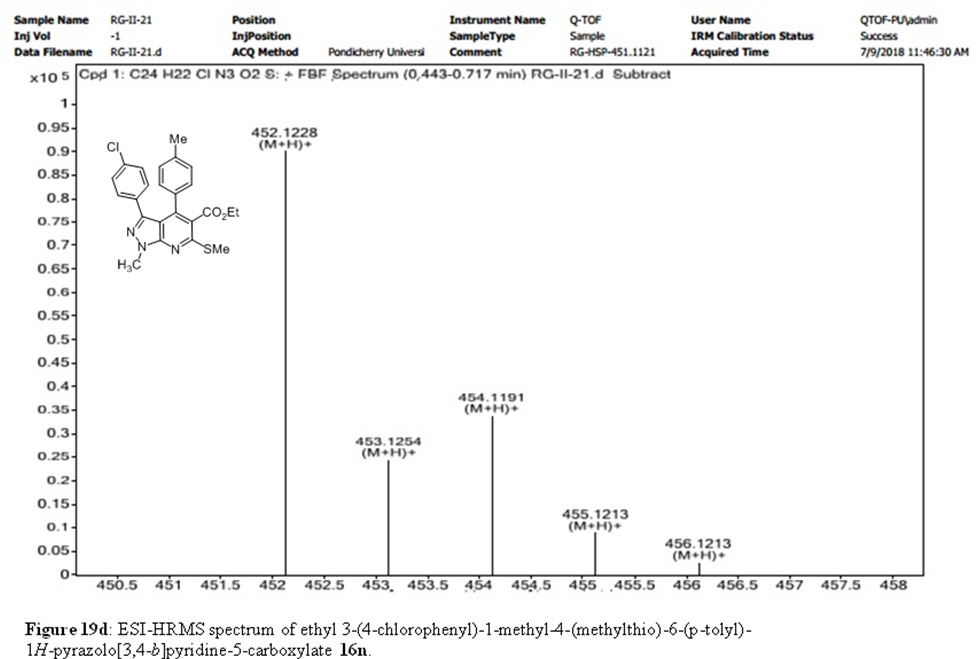

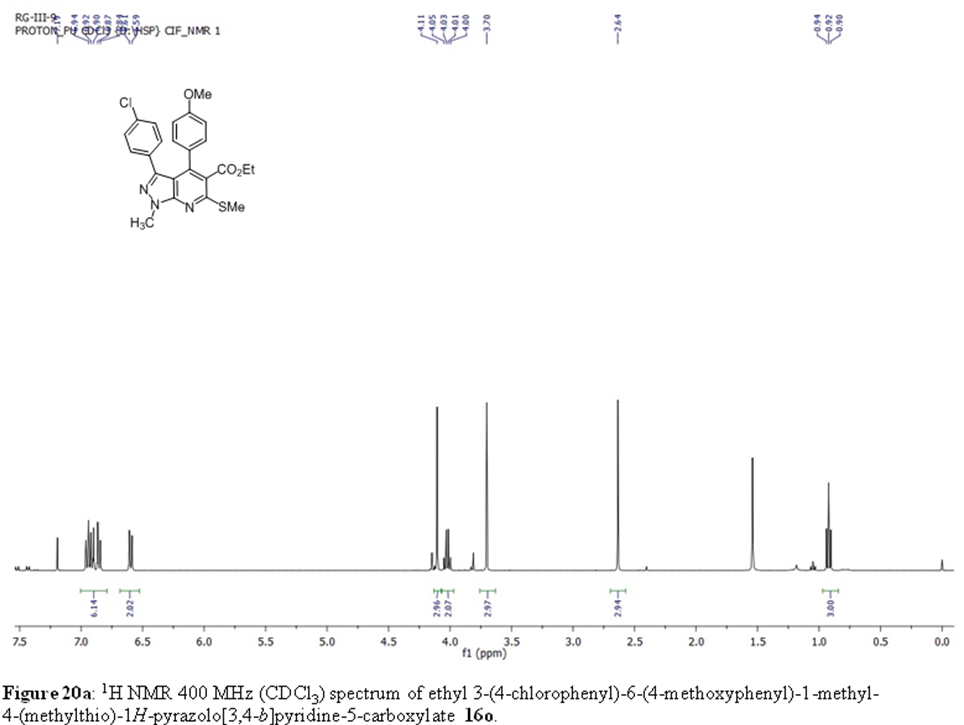

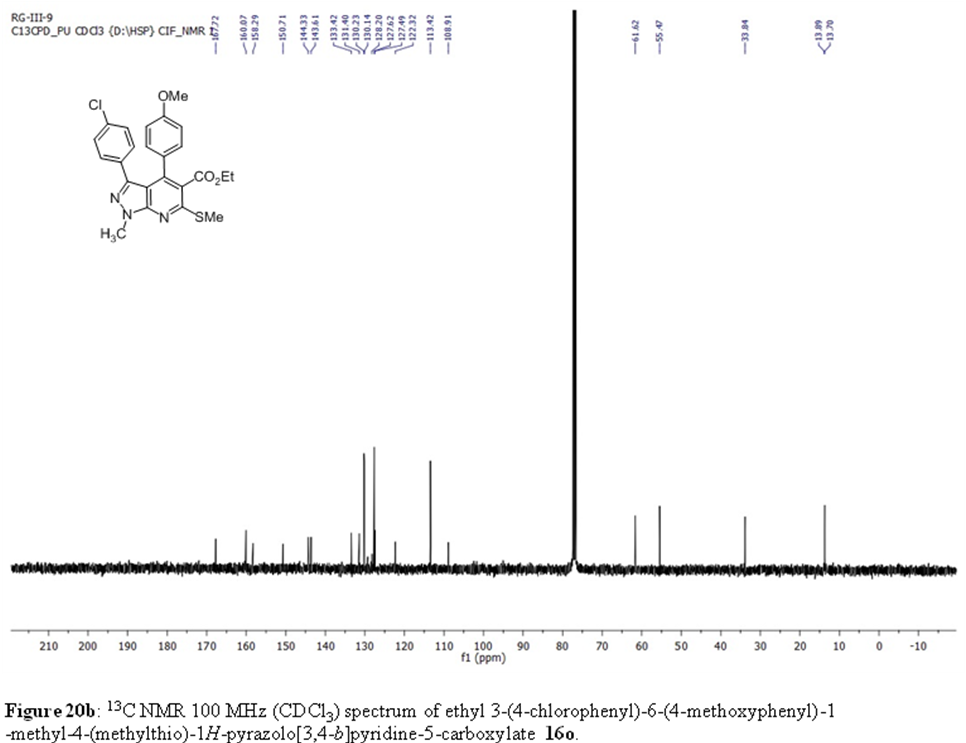

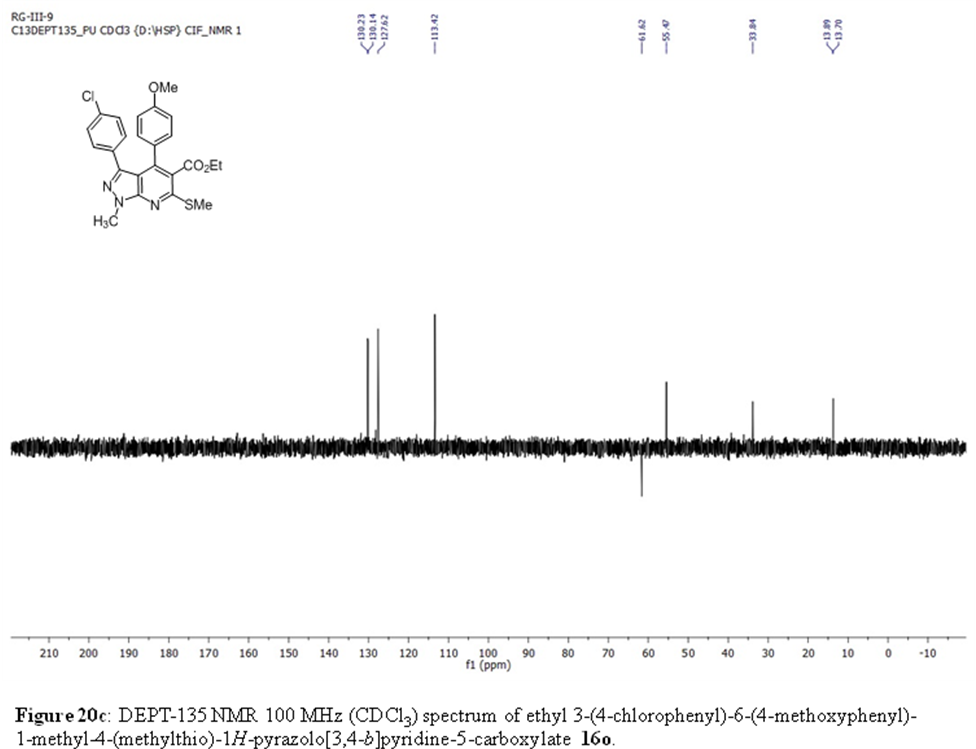

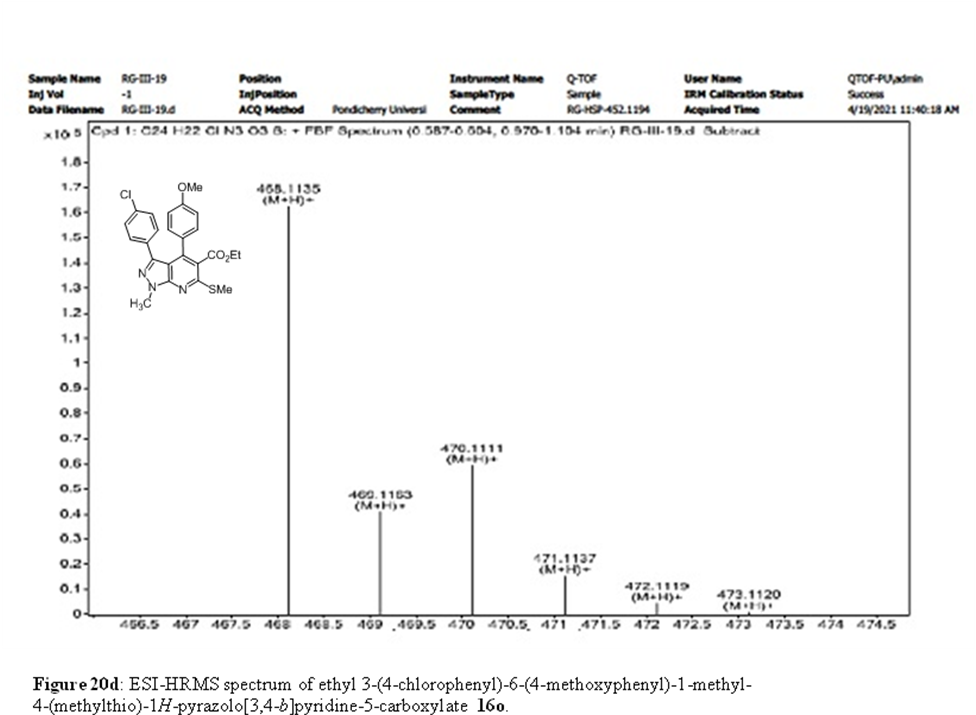

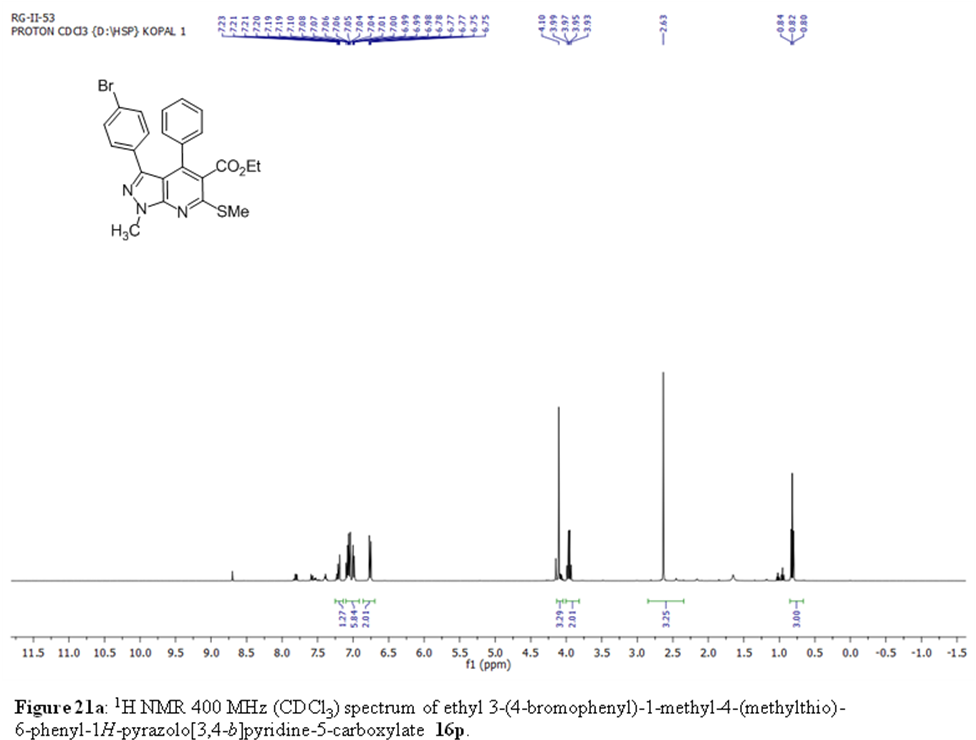

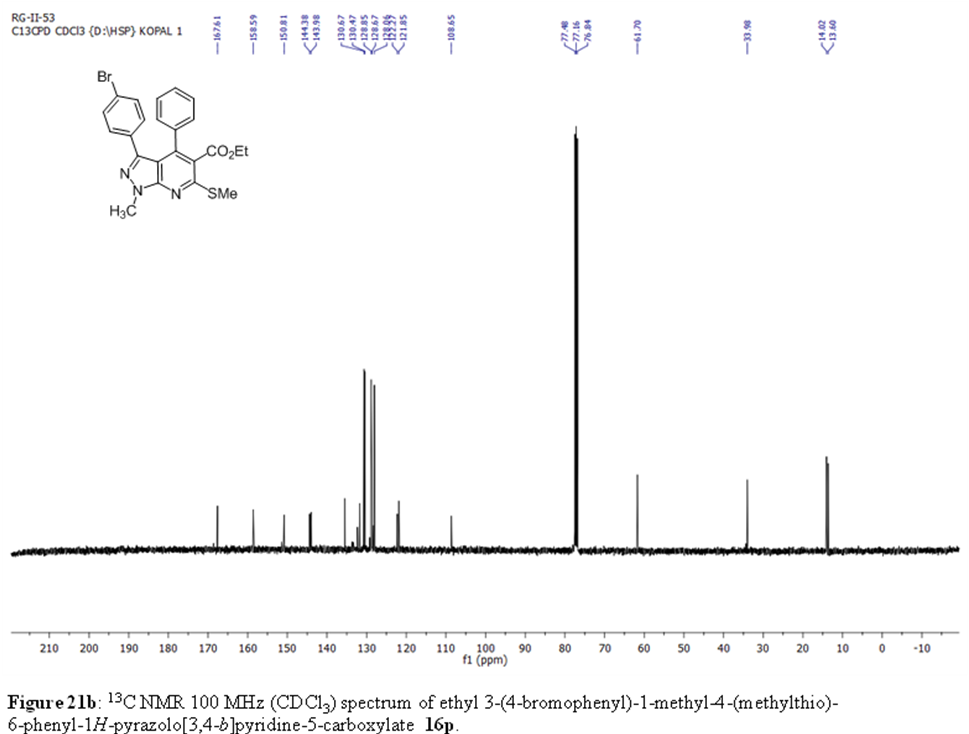

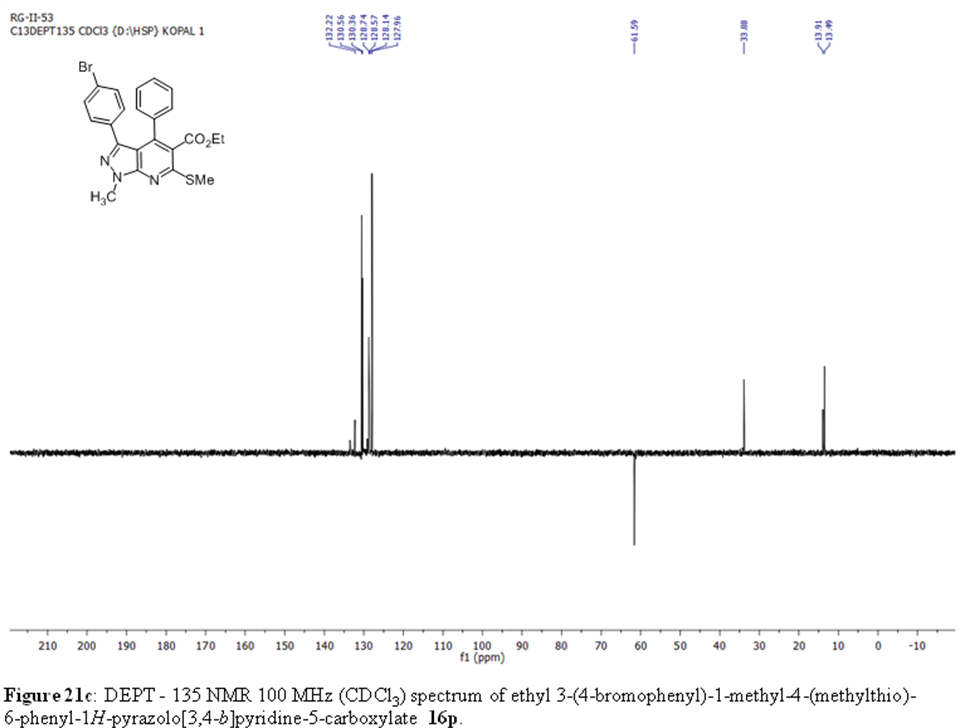

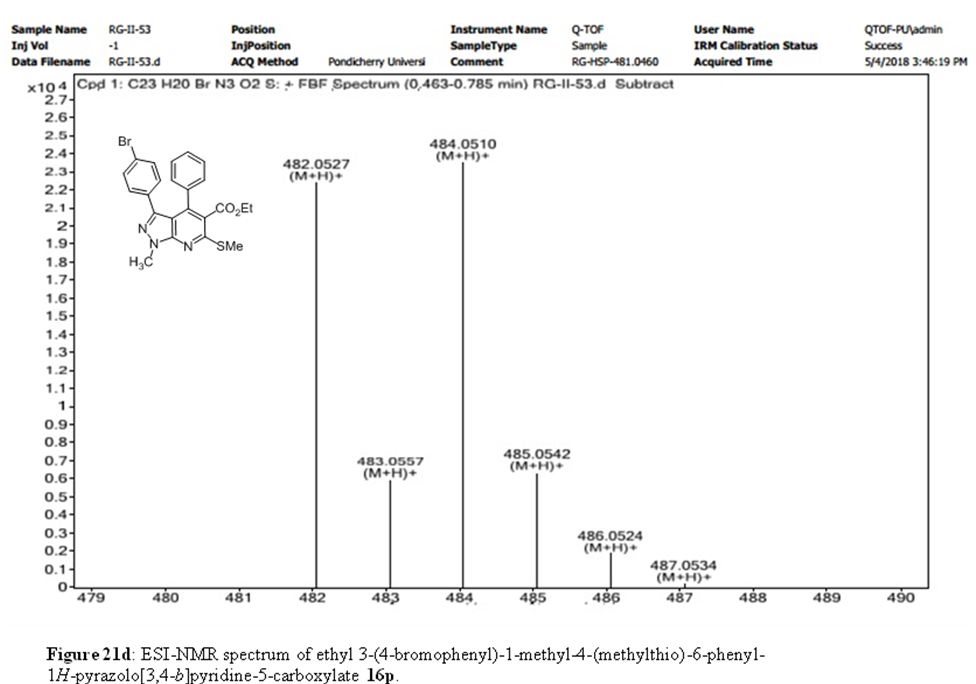

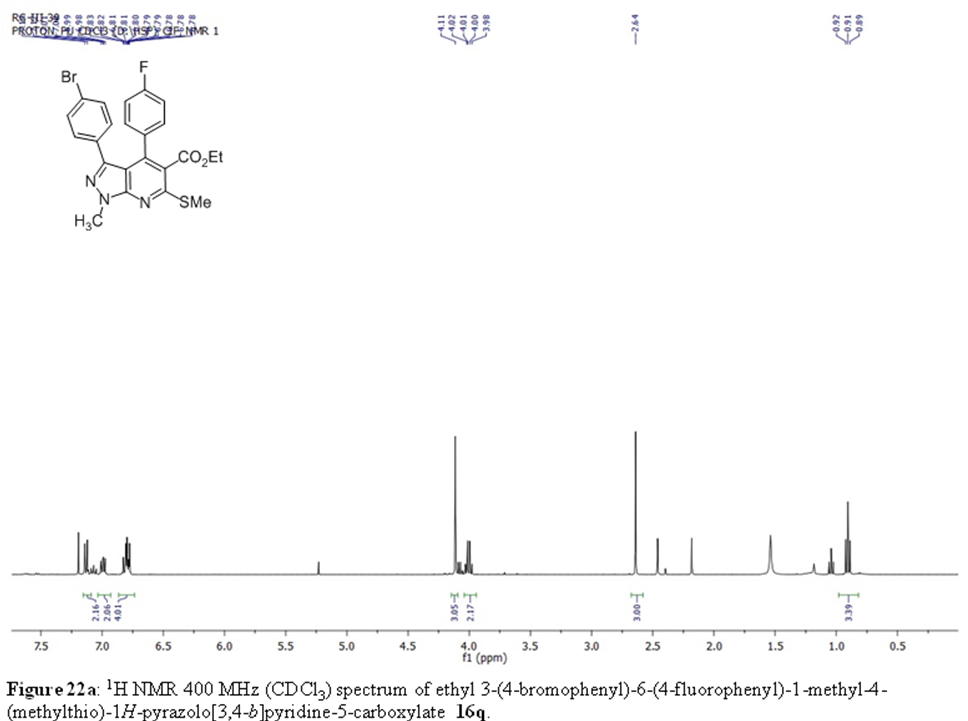

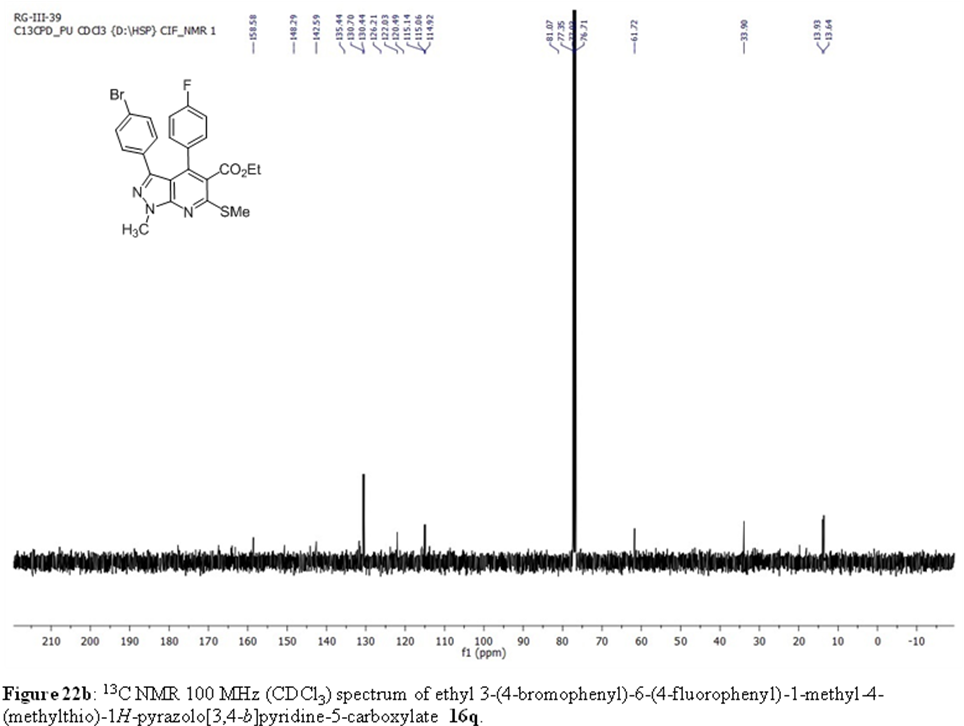

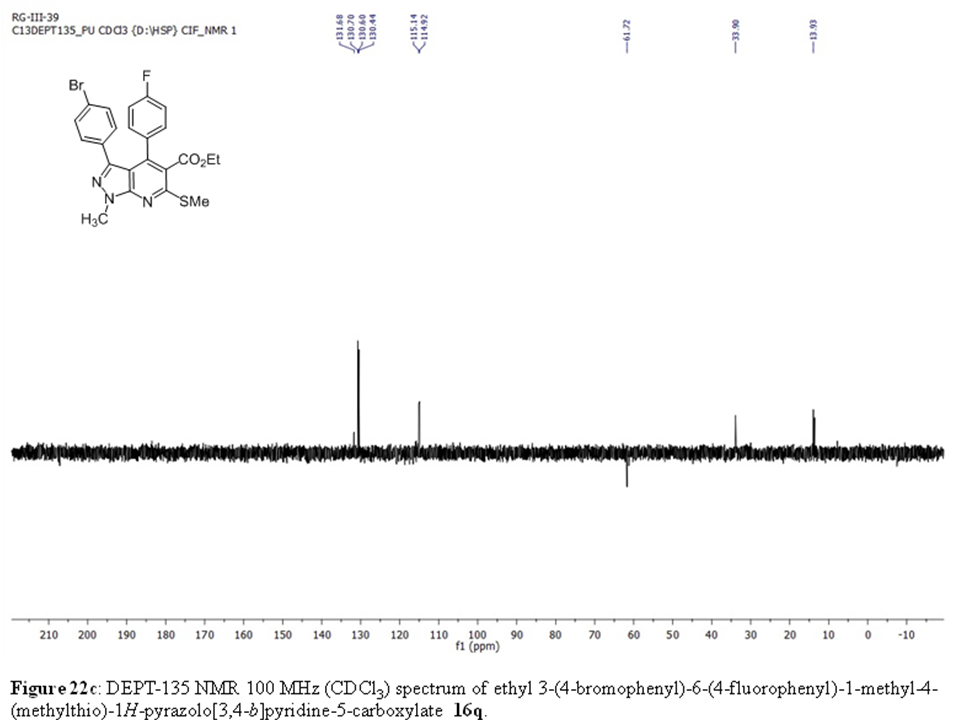

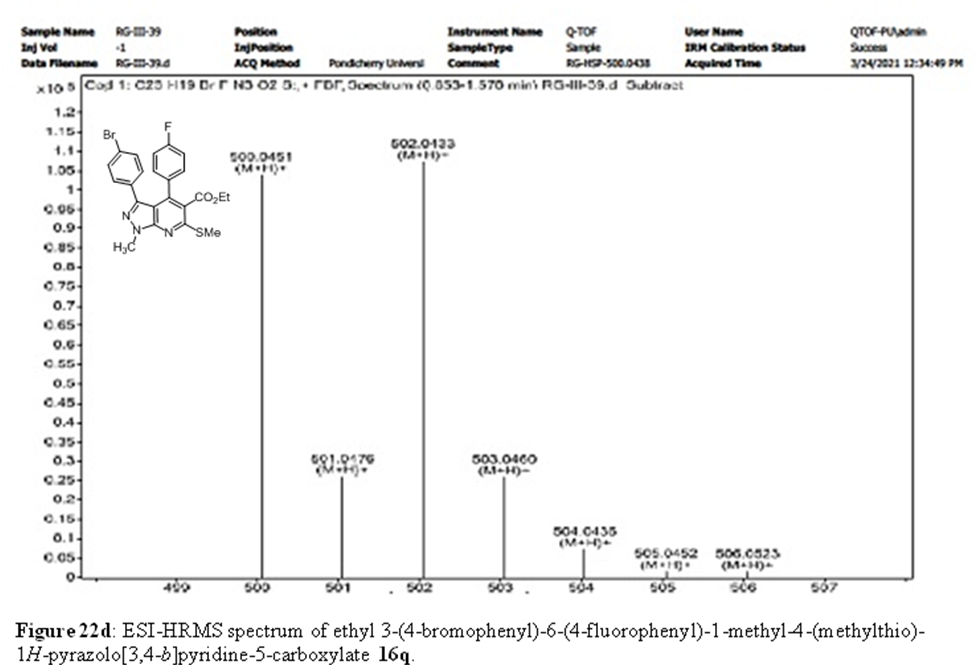

1. Berman, H.M., Westbrook, J., Feng, Z., Gilliland, G., Bhat, T.N., Weissig, H., Shindyalov, I.N. and Bourne, P.E., **2000**. The Protein Data Bank. *Nucleic Acids Research*, *28*, 235-242.doi: 10.1093/nar/28.1.235.
2. Forli, S., Huey, R., Pique, M.E., Sanner, M.F., Goodsell, D.S. and Olson, A.J., **2016**. Computational Protein–ligand Docking and Virtual Drug Screening with the AutoDock suite. *Nature Protocols*, *11*, 905-919.
3. Mooers, B.H., **2020**. Shortcuts for Faster Image Creation in PyMOL. *Protein Science*, *29*, 268-276.
4. Laskowski, R.A., **2011**.Swindells, M. B. LigPlot+: Multiple Ligand-Protein Interaction Diagrams For Drug Discovery. *J. Chem. Inf. Model*,*51*, 2778-2786.
